# Discovery of Tankyrase scaffolding inhibitor specifically targeting the ARC4 peptide binding domain

**DOI:** 10.1101/2025.03.31.646301

**Authors:** Chiara Bosetti, Albert Galera-Prat, Sven T. Sowa, Alexandra Gade, Cláudia Braga, Shoshy A. Brinch, Faranak Nami, Johan Pääkkönen, Veeti Pulju, Jo Waaler, Mads H. Clausen, Lari Lehtiö

**Affiliations:** Faculty of Biochemistry and Molecular Medicine & Biocenter Oulu, University of Oulu, Oulu, Finland; Norwegian Centre for Molecular Biosciences and Medicine, University of Oslo, Oslo, Norway; Department of Chemistry, Technical University of Denmark, Kgs. Lyngby, Denmark; Department of Immunology, Oslo University Hospital, Oslo, Norway; Hybrid Technology Hub-Centre of Excellence, Institute of Basic Medical Sciences, Faculty of Medicine, University of Oslo, Oslo, Norway

**Keywords:** Tankyrase, Ankyrin-repeat cluster, ARC, Inhibitor, Protein-protein interactions, pyrrolone, drug discovery

## Abstract

In the past, development of tankyrase inhibitors has focused on the ADP-ribosyltransferase domain. Targeting tankyrases ability to interact with protein substrates through their ARC domains represents an alternative strategy to be explored as a therapeutic approach against specific protein-protein interactions. In this paper, we employed a FRET-based assay to identify ARC4-binding compounds by screening the EU-OPENSCREEN Pilot and Commercials Diversity libraries. We discovered an effective series of compounds with the same scaffold and through chemical synthesis we obtained the compound **S8** (**ARCher-142**), which binds selectively to ARC4 with potency of 8 µM. NMR analysis and X-ray crystallography allowed us to identify the binding site in ARC4 and to rationalize the observed selectivity. Despite binding exclusively to ARC4, the inhibitor can attenuate the WNT/β-catenin signaling pathway in cells. Our work demonstrates that targeting single ARC domains is possible, offering an inhibition approach tailored to tankyrase ARC4 inhibition.

**Significance:** Tankyrases impact a variety of cellular processes by binding proteins through their ARC domains and the inhibition of these scaffolding functions represents an alternative therapeutic approach to catalytic inhibitors. With a FRET-based high-throughput screening of the EU-OPENSCREEN Pilot and Commercials Diversity libraries we discovered a pyrrolone-based scaffold that is interestingly selective towards ARC4, despite the high conservation of the ARC binding site. Our synthesized compound **S8** (**ARCher-142**) displays an 8 µM potency for TNKS2 ARC4. With NMR and X-ray crystallography we demonstrate that **S8** (**ARCher-142**) competes with the peptide optimized for binding and extends to a unique hydrophobic sub-pocket of ARC4. The compound attenuates the WNT/β-catenin signaling pathway in cells and interestingly offers the possibility to target specific protein-protein interactions mediated by ARC4, paving the way for the development of a pyrrolone-based class of tankyrase scaffolding inhibitors.

## Introduction

Ankyrin repeat (ANK) domains are ubiquitous protein modules designed to mediate protein-protein interactions. They are characterized by an ANK motif (30–34 residues), which is tandemly repeated and which structurally organizes into a helix-loop-helix architecture.^1,2^ Among the human enzyme family of Diphtheria toxin-like ADP-ribosyltransferases (ARTDs), tankyrases (paralogs TNKS1 and TNKS2) are the only members containing ANK domains, also defined as ankyrin repeat cluster (ARC) domains. Both tankyrases have five ARCs (ARC1/2/3/4/5), each of them characterized by five repetitions of the ANK motif.^3^ Once bound to the ARC domains, the protein partners can undergo poly-ADP-ribosylation (PARylation), a reaction that consists in the covalent attachment of ADP-ribose moieties from the substrate β-nicotinamide adenine dinucleotide (NAD^+^) into an elongated PAR chain. Tankyrases perform the PARylation reaction through their C-terminal catalytic domain (ART), whose function is possible only upon polymerization of multiple tankyrase molecules through their sterile alpha motif (SAM) domains (**Figure 1**).^4–7^ Targets that are subjected to this post-translational modification are subsequently ubiquitylated by E3 ubiquitin-protein ligase RNF146 and directed to proteasomal degradation.^8–12^

**Figure 1.**
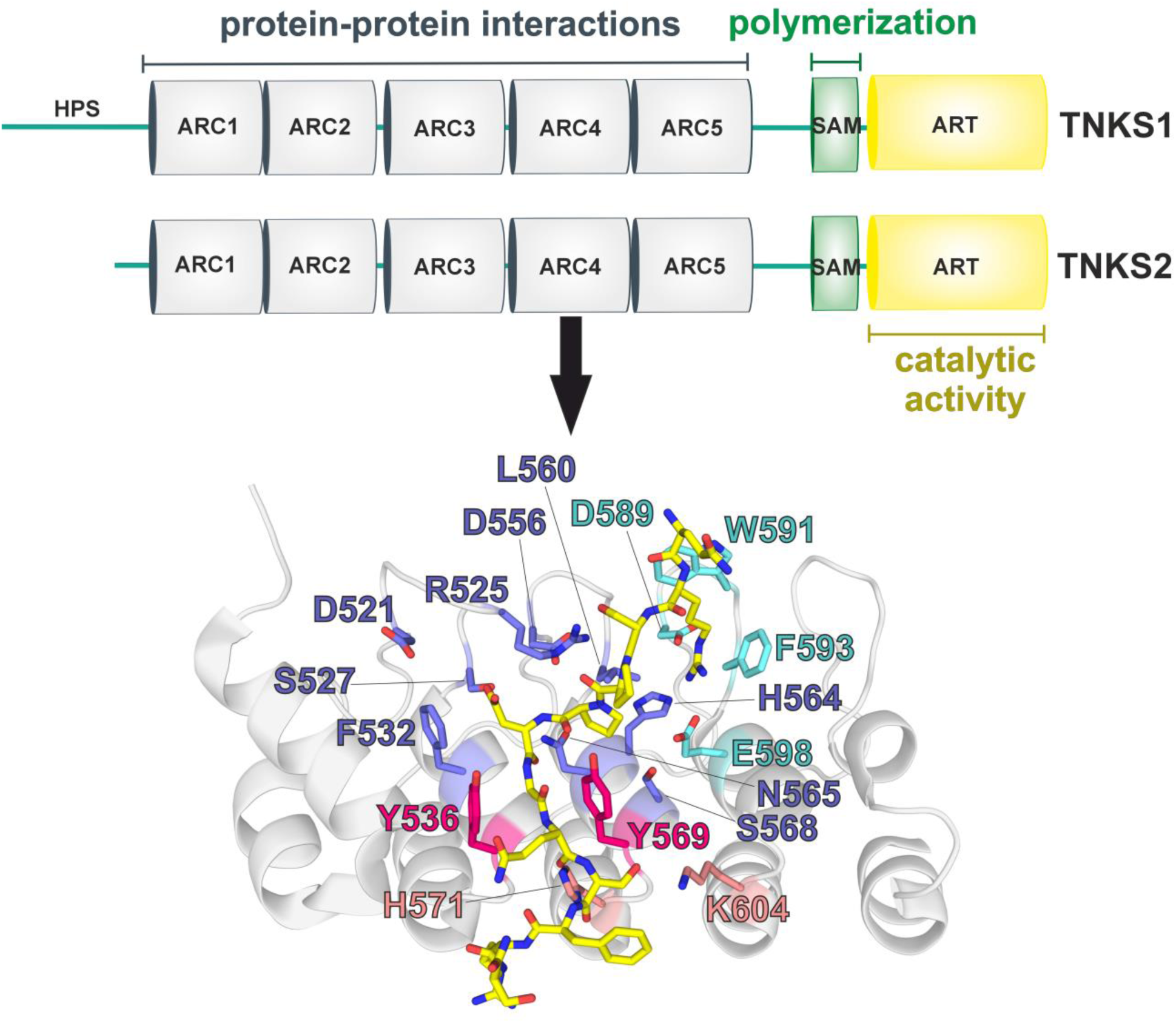
Domain organization of tankyrases and TNKS2 ARC4 peptide binding pocket. Tankyrases contain a C-terminal catalytic domain (ART), a SAM domain that mediates polymerization and five ARC domains responsible for protein-protein interactions. TNKS1 has an additional N-terminal histidine, proline and serine-rich region (HPS). The crystal structure of TNKS2 ARC4 in complex with the TBM of 3BP2 (PDB: 3TWR^3^) allowed to identify different regions of the binding pocket: the “arginine cradle” (residues in cyan), the “central patch” (blue), the “aromatic glycine sandwich” (magenta) and residues that offer additional contacts with the peptide (pink). The bound 3BP2 peptide is yellow.

Human tankyrase ARCs are highly similar and display a strong conservation throughout evolution that is particularly evident in the pocket for interaction with protein partners.^3,13^ This pocket can bind proteins provided with a specific 6–8 residue pattern defined as tankyrase-binding motif (TBM). The consensus rules of the TBM have been defined (R-X-X-[small hydrophobic or G]-[D/E/I/P]-G-[no P]-[D/E]) and indicate that arginine at position 1 and glycine at position 6 are essential for the binding to take place.^3^ Crystal structures of ARCs in complex with binding TBMs provided insights into the anatomy of the pocket (**Figure 1**).^3,14–18^ Notably, the mandatory arginine at position 1 of the TBM fits into an “arginine cradle”, while glycine at position 6 is accommodated into an “aromatic glycine sandwich”, which encompasses two tyrosine residues that form a gate crossed by the peptide. The central residues of the binding peptide occupy the “central patch”, and residues of the ARC domain can also provide additional interactions with the terminal residues of the TBM. Interestingly, ARC domains show different aptitudes for protein binding. ARC3 has no binding capacity and this behavior is reflected by its scarce conservation in comparison to other ARCs,^3,13^ leading to the hypothesis that ARC3 may play a structural role.^3,14,16^ ARC2 and ARC5 display stronger affinity towards TBMs in comparison to ARC1 and ARC4, where ARC1 is the weakest binder.^3^ Collectively, the five ARCs act as a multivalent platform that can adapt to binding partners.^16^

The ability to interact with and modify a variety of proteins allows tankyrases to orchestrate a vast range of cellular processes.^19–21^ By binding and PARylating paralogs AXIN1/2, tankyrases modulate the WNT/β-catenin pathway,^22^ a cascade of signaling events that affects processes like cell proliferation and differentiation.^23^ Deregulation of the WNT/β-catenin pathway contributes to the development of several diseases,^24^ encompassing diverse types of cancer^25^ and tissue fibrosis.^26^ Therefore, the regulating action of tankyrases on WNT/β-catenin signaling events can be potentially exploited as therapeutic strategy to re-tune the pathway^22,27^ and, in the past years, a wide range of catalytic inhibitors has been developed for this purpose.^28^ Inhibition of tankyrases has also proven to suppress YAP oncogenic activity^29–31^ and to potentiate cell sensitivity to DNA-damaging drugs by targeting MERIT40 PARylation.^32^ Tankyrases also regulate telomere length,^33–35^ apoptosis,^36^ mitotic spindle assembly,^37–39^ vesicular transport^38,40^ and glucose metabolism.^40–43^

In the past, research efforts have focused on the discovery of catalytic inhibitors. Although there are reports of dose-limiting intestinal toxicity in mice,^44–46^ optimized lead shows no substantial toxicity and a clear therapeutic window.^47^ The development of scaffolding-targeting compounds can provide new routes to explore tankyrase inhibition, with the possibility to impact distinct biological activities. While inhibitors of SAM-mediated multimerization have not been reported yet, macrocyclized peptides and some ARC inhibitors have already been developed.^48–51^

In this study, we screened the EU-OPENSCREEN Pilot Library and Commercials Diversity collection of compounds by employing a fluorescence resonance energy transfer (FRET) based assay designed for the discovery of novel compounds targeting scaffolding functions of tankyrases.^52^ We aimed at the identification of compounds that target the TBM pocket by inhibiting the interaction between TNKS2 ARC4 and the peptide optimized for ARC binding (REAGDGEE).^3^ The same libraries were tested in a counter screen for inhibition of SAM domain dimer formation to identify non-specific compounds, while simultaneously assessing possible hit compounds that would interfere with the SAM-SAM interaction. We identified a common scaffold in the best hits and defined their binding site to TNKS2 ARC4 with NMR titration experiments. Subsequently, analogous compounds were acquired commercially or synthesized and analyzed for the study of structure-activity relationships (SARs). One synthesized analogue **S8** displayed increased potency, and we solved its co-crystal structure with TNKS2 ARC4. We reveal that the compound occupies the TBM pocket and it extends towards a region that is unique of TNKS2 ARC4. Profiling of the scaffold with ARC domains of TNKS1 and TNKS2 explained the compound selectivity towards TNKS1/2 ARC4. In addition, **S8** is active in cells as it attenuates the WNT/β-catenin signaling pathway. These findings indicate that it is possible to develop inhibitors tailored towards the inhibition of specific ARC-mediated interactions and affecting potentially distinct cellular pathways. Overall, this study provides novel insights into the inhibition of tankyrase scaffolding functions and opens the route to the development of a new class of tankyrase inhibitors.

## Results

### Discovery of TNKS2 ARC4 binders through FRET-based high-throughput screening

To discover novel compounds targeting the TBM pocket in ARC domains, we performed a high-throughput screening based on a FRET assay designed for the purpose.^52^ In the assay, we employed human recombinant TNKS2 ARC4 fused with mCerulean (CFP) and the 8-residues peptide optimized for the binding of ARCs (REAGDGEE) fused with mCitrine (YFP).^3^ The interaction is evaluated by calculating the ratiometric FRET (rFRET), which is the ratio of the fluorescent intensity of the acceptor at the emission wavelength to the fluorescent intensity of the donor at emission wavelength. The control indicating total inhibition of the interaction (positive control) is represented by the same FRET pair in presence of 1 M guanidine hydrochloride. At the same time, we employed the FRET assay developed for the discovery of compounds inhibiting SAM-SAM interaction to screen the same libraries and to identify overlapping, hence non-selective, hits.

We first tested the compounds of the EU-OPENSCREEN Pilot Library, which encompasses 4928 compounds including bioactive compounds and representative members of the Commercials Diversity library. This pilot screen was used to set a threshold of 20% activity for hit determination. The screening resulted in the identification of 50 primary hits (1% hit rate), active at a concentration of 30 µM, which were later analyzed with DSF assay. As the assay performed well, the larger EU-OPENSCREEN Commercials Diversity collection of compounds was screened. A total of 96076 compounds were screened, identifying 772 primary hits (0.8% hit rate). Out of 772 primary hits, 12 were found to overlap with hits of the counter screen for the SAM-SAM interaction, so they did not undergo further validation. As a result, 760 hits were subjected to validation through potency measurements, where compounds were studied at different concentrations ranging from 0.5 µM to 100 µM. A primary hit was considered validated when the calculated IC_50_ value fell in the 0.5–100 µM range and when its activity change was at least 25%. Out of 760 compounds, 114 were validated and were further tested with DSF assay. Complete data of the screening is available at the European Chemical Biology Database (ECBD). ^53,54^

### Pyrrolone-based compounds inhibit and are selective towards TNKS1/2 ARC4

We used DSF assay to validate the binding of the primary hits to TNKS2 ARC4. A total of 50 primary hits from the Pilot Library and 114 compounds from the Commercials Diversity Library were tested (**Figure S1, S2**). Two compounds from the Pilot Library were found to mildly stabilize TNKS2 ARC4 with ΔTm equal to 0.45 °C and 0.41 °C (**Figure S1**). Validation of primary hits from the Commercials Diversity collection led to the identification of nine molecules that share a common pyrrolone-based scaffold and that remarkably stabilized the protein up to 2.25 °C (**Figure S2**).

Given the notable stabilization of the pyrrolone-based compounds from the Commercials Diversity collection, we decided to focus on their common scaffold for compound optimization. Pyrrolone is indeed a versatile moiety that is largely employed for the design of compounds with a variety of biological effects.^55^ The nine primary hits were acquired from commercial suppliers, they underwent quality control, additional DSF validation and determination of potency through FRET-based IC_50_ measurements (**Table 1**). All the tested compounds include a benzodioxane moiety (R^1^), a phenyl group with substitutions in the *para* position (R^2^), and an R^3^ group with an aliphatic character and a tertiary amine. All the compounds studied significantly stabilized TNKS2 ARC4, with ΔTm values ranging from 1.40 °C to 4.58 °C. In addition, all molecules, except for **7**, have IC_50_ values below 100 µM. Among the nine molecules, **1** performed the best in terms of stabilization of TNKS2 ARC4 and potency. Direct comparison of **1** with **2** indicates that a polar hydrogen bond donor group (-OH) in the *para* position of the phenyl at R^2^ is preferred in comparison to a fluorine substituent. In the same position, the presence of a hydrophobic methyl group (**3, 4**), other halogen substituents (**5**) and alkoxy groups (**6**–**9**) is also tolerated, even if not optimal. Focusing on the R^3^ group, all the nine primary hits have a common 3-carbon chain and comparisons between **3**–**4**, and **8**–**9**, show that the morpholine-based moiety performs slightly worse than the dimethylamino-based substituent.

**Table 1.**
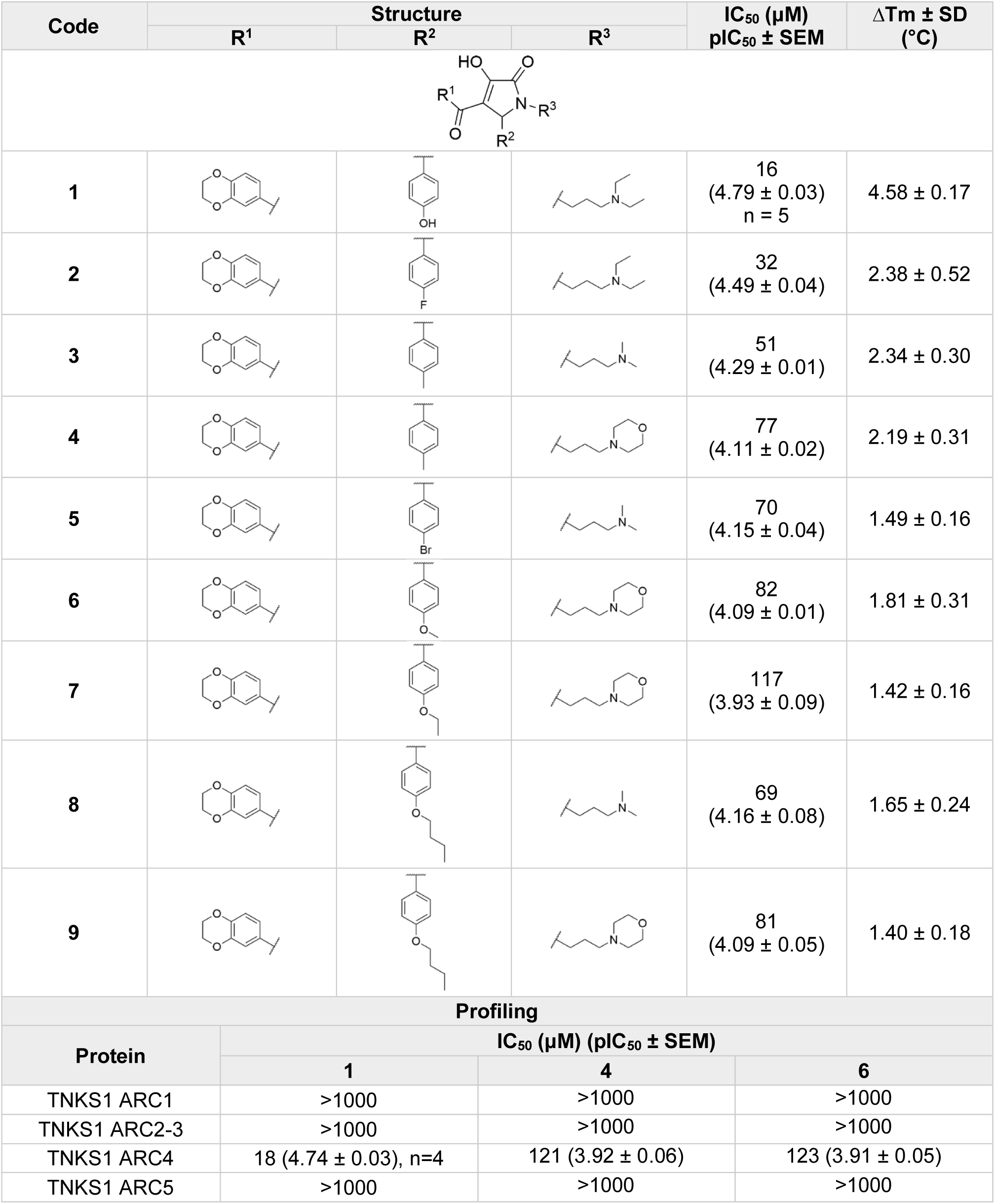
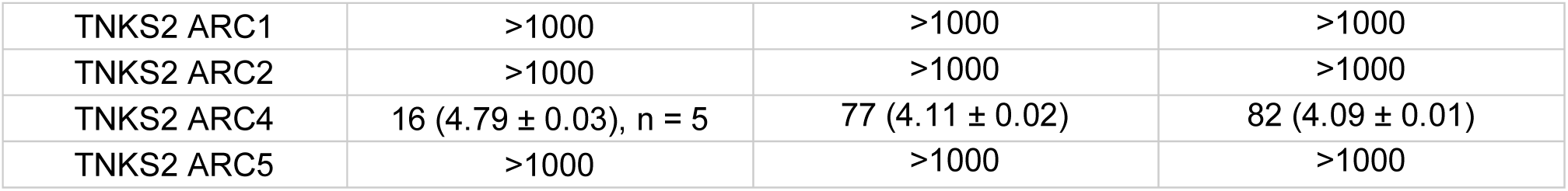
The nine primary hit compounds from the Commercials Diversity collection share a common pyrrolone scaffold and profiling of compounds **1**, **4**, **6** against TNKS1/2 ARC domains reveals selectivity of the scaffold towards TNKS1/2 ARC4 domains. FRET-based potency measurements are indicated as IC_50_ (μM) and pIC_50_ ± SEM, with number of repetitions n=3 unless indicated otherwise (n=1 for measurements with IC_50_ >1000 μM). Thermal stabilization (ΔTm ± SD) is also reported (n=4).

| Code | Structure |  |  | IC<sub>50</sub> (μM)<br>pIC<sub>50</sub> ± SEM | ΔT<sub>m</sub> ± SD<br>(°C) |
| --- | --- | --- | --- | --- | --- |
|  | R<sup>1</sup> | R<sup>2</sup> | R<sup>3</sup> |  |  |
| **1** |  |  |  | 16<br>(4.79 ± 0.03)<br>n = 5 | 4.58 ± 0.17 |
| **2** |  |  |  | 32<br>(4.49 ± 0.04) | 2.38 ± 0.52 |
| **3** |  |  |  | 51<br>(4.29 ± 0.01) | 2.34 ± 0.30 |
| **4** |  |  |  | 77<br>(4.11 ± 0.02) | 2.19 ± 0.31 |
| **5** |  |  |  | 70<br>(4.15 ± 0.04) | 1.49 ± 0.16 |
| **6** |  |  |  | 82<br>(4.09 ± 0.01) | 1.81 ± 0.31 |
| **7** |  |  |  | 117<br>(3.93 ± 0.09) | 1.42 ± 0.16 |
| **8** |  |  |  | 69<br>(4.16 ± 0.08) | 1.65 ± 0.24 |
| **9** |  |  |  | 81<br>(4.09 ± 0.05) | 1.40 ± 0.18 |
| **Profiling** |  |  |  |  |  |
| Protein | IC<sub>50</sub> (μM) (pIC<sub>50</sub> ± SEM) |  |  |  |  |
|  | **1** | **4** | **6** |  |  |
| TNKS1 ARC1 | >1000 | >1000 | >1000 |  |  |
| TNKS1 ARC2-3 | >1000 | >1000 | >1000 |  |  |
| TNKS1 ARC4 | 18 (4.74 ± 0.03), n=4 | 121 (3.92 ± 0.06) | 123 (3.91 ± 0.05) |  |  |
| TNKS1 ARC5 | >1000 | >1000 | >1000 |  |  |

|  |  |  |  |
| --- | --- | --- | --- |
| TNKS2 ARC1 | >1000 | >1000 | >1000 |
| TNKS2 ARC2 | >1000 | >1000 | >1000 |
| TNKS2 ARC4 | 16 (4.79 ± 0.03), n = 5 | 77 (4.11 ± 0.02) | 82 (4.09 ± 0.01) |
| TNKS2 ARC5 | >1000 | >1000 | >1000 |

Profiling of **1** against a panel of ARC domains from TNKS1 and TNKS2 interestingly indicates that the compound inhibits the interaction of the optimized peptide only with TNKS1/2 ARC4 and not with other ARCs (**Table 1**). Profiling of other primary hits **4** and **6** also shows selectivity towards TNKS1/2 ARC4, confirming the selectivity of the scaffold for ARC4 inhibition (**Table 1**).

### Pyrrolone-based compounds bind to a common site in TNKS2 ARC4 binding pocket

To identify the binding site of the compounds and the reason behind the observed selectivity, we performed ^1^H-^15^N HSQC NMR titration experiments with ^15^N-labelled TNKS2 ARC4 and compounds **1**, **4** and **6**. Compounds appear to bind with intermediate exchange rate to a common binding site (**Figure 2**). This site overlaps with the TBM pocket and intriguingly extends to a sub-pocket with a hydrophobic character (**Figure 3A**-**B**). The bottom of this sub-pocket is indeed decorated with a leucine and a phenylalanine residue (L497, F532) that may provide the correct hydrophobic environment to accommodate the aliphatic R^3^ group of the compounds, while the tertiary amine at R^3^ may extend towards other residues that surround the sub-pocket. Notably, residue F532 is characterized by a significantly high chemical shift perturbation (CFP), possibly due to the vicinity of the pyrrolone core, which is likely located at the center of the identified binding area. The alignment of the sequences of ARC domains indicates that phenylalanine at this position is conserved only in TNKS1/2 ARC4 and TNKS1/2 ARC1, where other ARCs contain leucine instead (**Figure 3C**). It is necessary to highlight that ARC1 domains show an overall lower conservation of the residues lining the binding pocket and are weaker peptide binders in comparison to ARC2/4/5. This is mainly due to the presence of phenylalanine residues instead of the two tyrosines at the “aromatic glycine sandwich” present in other ARCs. Since the “aromatic glycine sandwich” and neighboring residues form the binding pocket of the analyzed compounds, the described absence of tyrosines may explain the lack of binding of the compounds to ARC1 domains. Additional differences between TNKS1/2 ARC4 and other ARC domains in the binding pocket of the compounds are provided by the loop lining the sub-pocket (R520 to T528). Specifically, positions occupied by residues I522 and E523 in TNKS2 ARC4 (L673 and E674 in TNKS1 ARC4) vary among other ARCs (**Figure 3C**).

**Figure 2.**
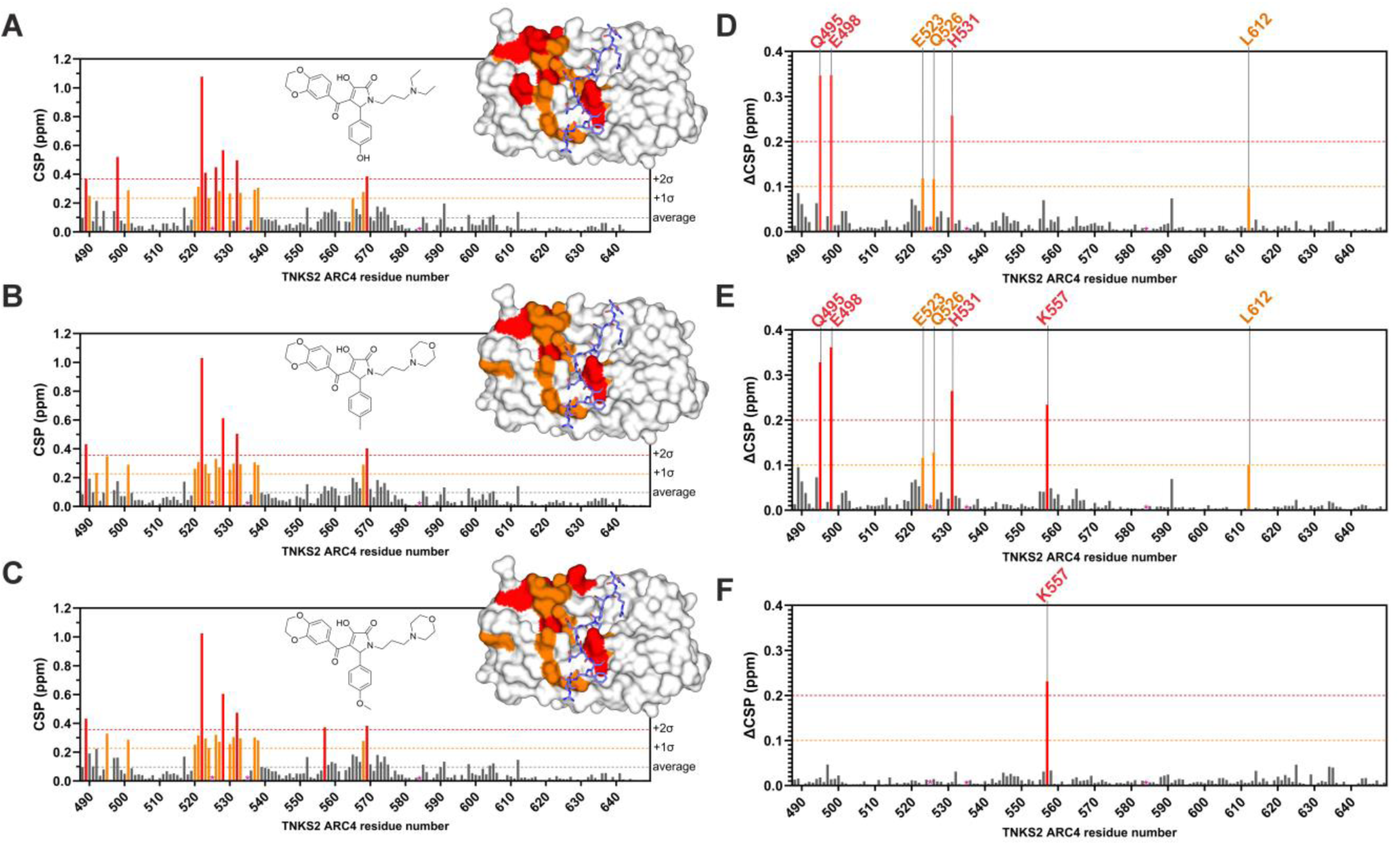
Analysis of NMR titration experiments. CSPs of TNKS2 ARC4 induced by binding of compounds **1** (**A**), **4** (**B**) and **6** (**C**). Each plot is accompanied by the structure of the titrated compound and the crystal structure of TNKS2 ARC4 bound to the TBM of 3BP2 (PDB: 3TWR^3^). On the surface of the protein, residues significantly perturbed are indicated in orange (2σ < CSP ≤ 1σ) and red (CSP ≥ 2σ). The TBM peptide of 3BP2 is shown (blue) to highlight the overlapping between its binding site and the binding site of the compounds. To identify significant differences in the binding sites of **1**, **4** and **6**, the Euclidean distances between the residues peaks at saturation have been calculated (ΔCSP) and plotted (**D**, **E**, **F**). This approach highlights peaks that underwent different changes in the environment, including peaks that covered different distances and peaks that shifted to different locations. Relevant residues are indicated and colored in orange (ΔCSP ≥ 0.1 ppm) and red (ΔCSP ≥ 0.2 ppm). (**D**) Plot indicating ΔCSPs induced by compounds **1** and **4**. (**E**) Plot indicating ΔCSPs induced by compounds **1** and **6**. (**F**) Plot indicating ΔCSPs induced by compounds **4** and **6**. In all plots, residues that could not be assigned (R525, G535, A584) are shown with * (color magenta). Proline residues spots on the plots are left empty.

**Figure 3.**
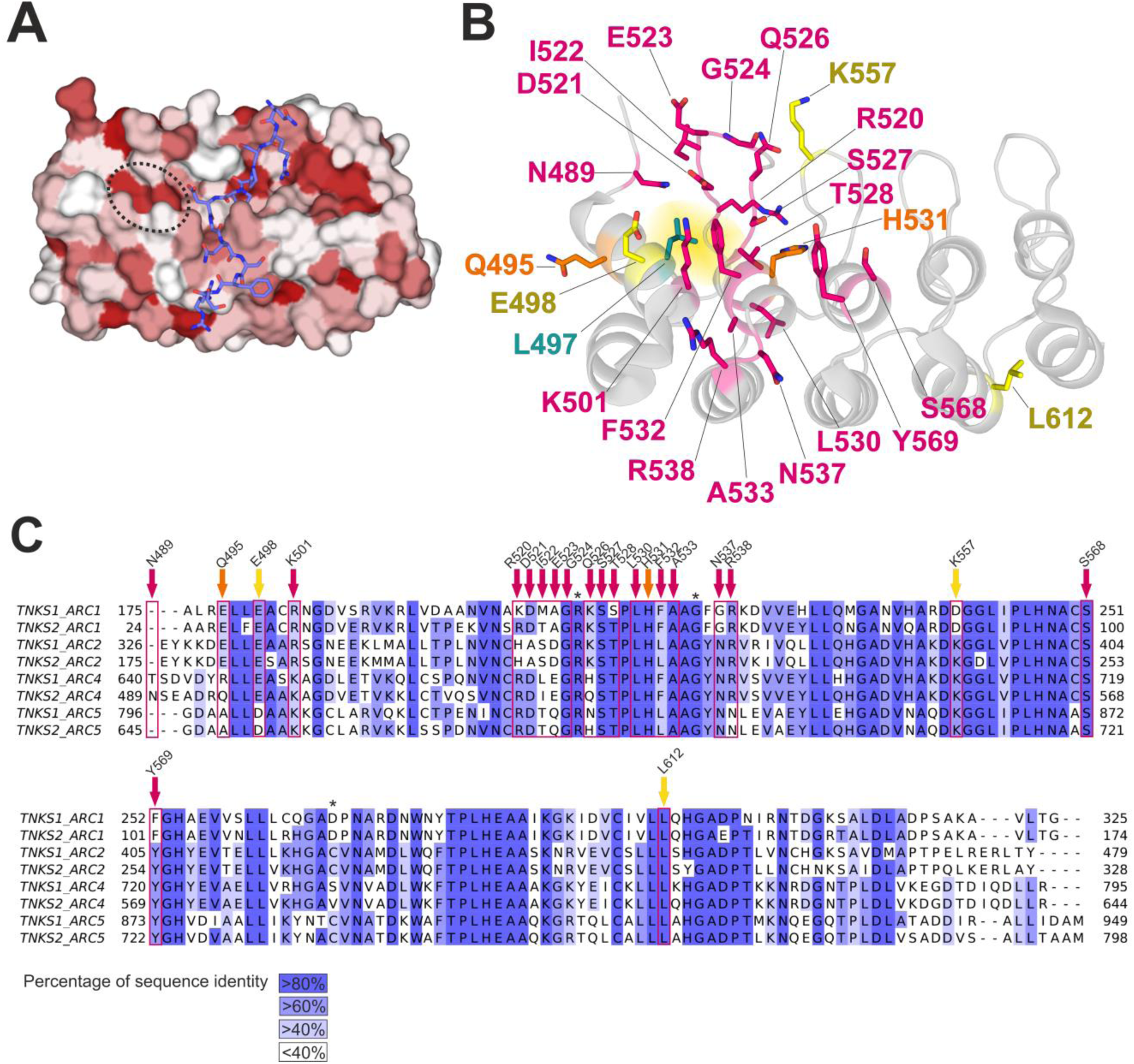
Identification and characterization of the compounds binding site in TNKS2 ARC4. (**A**) Crystal structure of TNKS2 ARC4 (PDB: 3TWR^3^) with bound TBM of 3BP2 (blue). The surface of the protein is colored in a gradient from white to red according to the Kyte-Doolittle hydrophobicity scale, where red indicates residues with high hydrophobic score. The hydrophobic pocket lined by L497 and F532 is indicated with a dotted line. (**B**) Crystal structure of TNKS2 ARC4 (PDB: 3TWR^3^) with residues that were significantly perturbed upon compound titration shown as sticks. Residues that likely form the shared binding site of the compounds, that is residues with CSP ≥ 1 σ in all titrations, are shown in magenta. Residues Q495 and H531 appear to undergo significant environmental changes with compounds **4** and **6** only and are colored orange. Residues that display different behavior with one compound titration only are shown in yellow. Residue E498 has significantly high CSP induced only by **1**, while residue L612 moves to a different location with **1**. Residue K557 appears instead to be perturbed significantly only upon compound **6** titration. Residue L497 is shown in green, and it forms the floor of the hydrophobic pocket (yellow halo) with H532. (**C**) Sequence alignment of TNKS1/2 ARCs. Residues are highlighted with a blue color gradient according to the sequence identity. Residues that appear to be perturbed upon compound binding are indicated with arrows in the same color scheme as in (**B**). TNKS1/2 ARC3 sequences are excluded from the alignment since they are reported not to bind TBMs of protein partners. Residues that were not assigned in NMR titrations are indicated with *.

The NMR titration experiments reveal that **1**, **4** and **6** bind TNKS2 ARC4 in the same fashion and only some residues appear to have different roles in the binding sites. Residues with significantly different behavior were identified by calculating the Euclidean distances between the peak positions at compound saturation (ΔCSP) (**Figure 2D**-**F**). The compounds differ only in their R^2^ and R^3^ groups and share the core and the benzodioxane moiety (R^1^). Compounds **4** and **6** share a morpholine-based R^3^ group, which provides a heterocyclic ring in comparison to the R^3^ group of **1**. All three compounds have a three-carbon aliphatic chain in the R^3^ group that can be accommodated in the hydrophobic sub-pocket. Consequently, the presence of the morpholine moiety may explain the involvement of residue Q495 with **4** and **6**, while E498 is affected only by **1** binding, indicating a possible ion-pair interaction with the positively charged tertiary amine. In addition, residue H531 displays significantly higher CSPs in **4** and **6** titrations, while its CSP in titration with **1** appears to be below the calculated average. Analogue **6** has a bulkier methoxy substitution on the *para* position of the phenyl ring at R^2^ and the high CSP of K557 may indicate the substituent pointing in the direction of the residue. Other minor differences in CSPs involve E523 and Q526, which are more perturbed in the presence of **1**. In addition, residue L612 has comparable CSPs in the titrations with the different compounds, but its peak appears to shift to a different location with **1** only.

### Synthesis of pyrrolone-based analogues

To explore the potential of the pyrrolone scaffold, analogues of **1** with different R^1^, R^2^ and R^3^ substituents (**Table S1**, compounds **S1**–**S12**) were synthesized using a two-step procedure, as depicted in **Scheme 1**. Aryl methyl ketones **1a**–**c** were treated with a dialkyl oxalate under basic conditions, yielding 2,4-diketoesters **2a**–**c** via a crossed Claisen condensation.^56^ Pyrrolone analogues were synthesized using commercially available substituted benzaldehydes, substituted primary amines and 2,4-diketoesters **2a**–**c**, through a one pot three-component reaction,^57,58^ yielding the desired products in relatively low yields (3–22%). In addition to pyrrolone ring cyclization, an undesired competing side reaction was observed, in which 2,4-diketoesters underwent aminolysis with primary amines. Compound **S9** was synthesized by reducing the commercially available *para*-nitro analogue with zinc dust and ammonium chloride by heating a H₂O/EtOH solution to 80 °C. ^1^H NMR spectra of the synthesized analogues show characteristic singlet signals for the CH proton at the 5-position at 5.2–5.4 ppm (**Figure S3**-**S31**). Additionally, the CH_2_ protons from the alkyl chain substituent at R^3^, positioned close to the chiral center are diastereotopic and exhibit distinct chemical shifts.

**Scheme 1.**
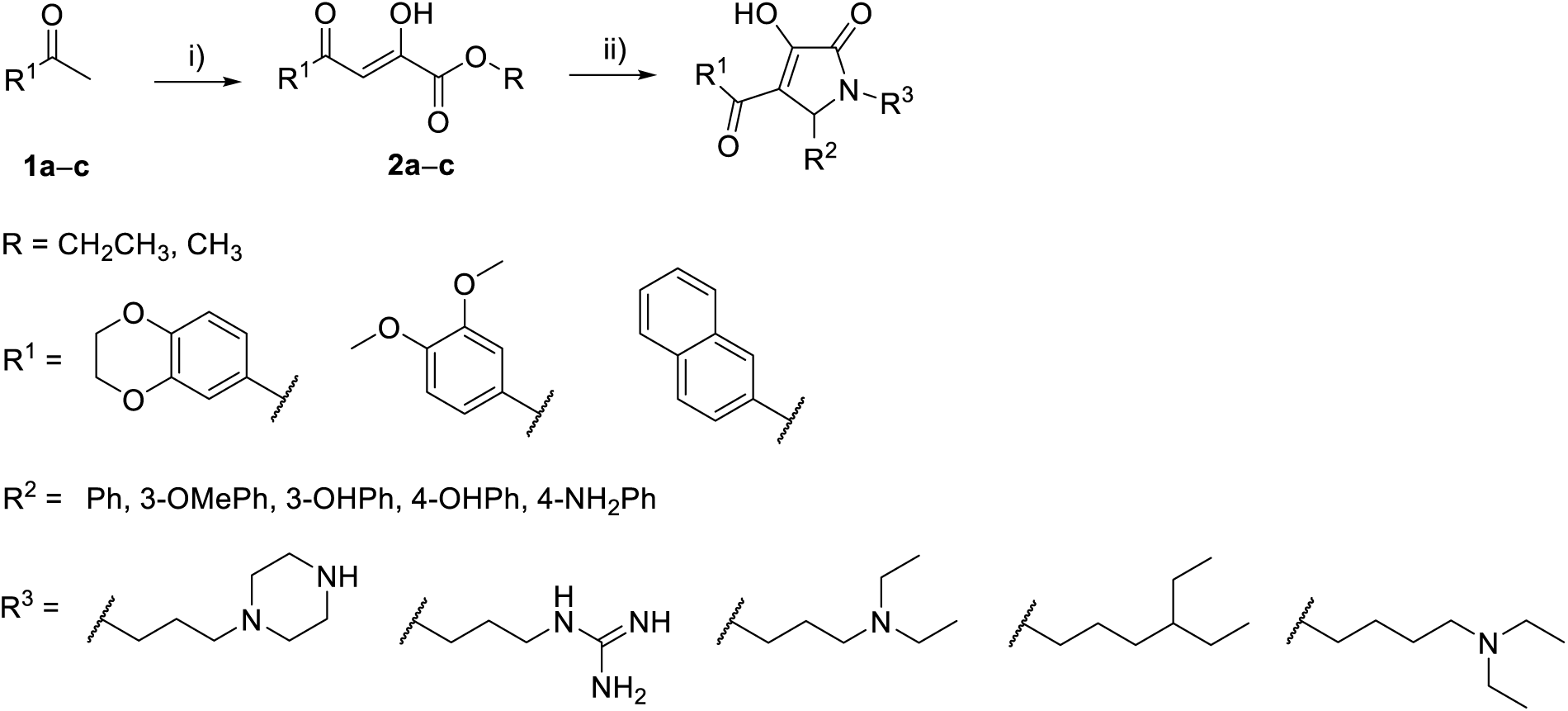
General synthetic route for the synthesis of pyrrolone analogues. Reagents and conditions: i) dialkyl oxalate, NaH, THF, Δ; ii) R^2^CHO, R^3^NH_2_, THF, EtOH/AcOH or AcOH/NH_4_Ac, rt or Δ.

### SAR investigation

A SAR investigation was conducted with purchased and synthesized analogues of **1** through FRET-based potency measurements and DSF assay with TNKS2 ARC4 (**Table 2**, **Table S2**). All the compounds commercially acquired underwent quality control. Modifications at R^1^ indicate that the presence of a double ring system with at least one oxygen is essential for maintaining high inhibitory activity and stabilization of the protein (**10** IC_50_ 31 µM, **11**/**S6**/**S7** IC_50_ > 500 µM). Diverse variations of the substituents on the phenyl ring at R^2^ appear to be tolerated, including substituents in the *meta* position and large alkoxy substituents as seen in **Table 1** (**8** IC_50_ 69 µM, **9** IC_50_ 81 µM). Notably, the methoxy group in the *meta* position and the dimethylamino group in the *para* position moderately improve potency and allow a stabilization of the protein ΔTm > 4 °C (**12** IC_50_ 11 µM, **19** IC_50_ 10 µM). Changes to the length of the aliphatic chain at R^3^ greatly impacts both potency and protein stabilization. Comparison of compounds **13** (IC_50_ 24 µM) with **14** (IC_50_ 139 µM) and **16** (IC_50_ 31 µM) with **17** (IC_50_ 274 µM) indicates indeed that a 3-carbon aliphatic chain is superior to a 2-carbon chain. At the same time, a longer 4-carbon chain has a negative impact on the measured values (**S12** IC_50_ 74 µM). The diethylamino-based substituent at R^3^ (**13** IC_50_ 24 µM) has slightly improved potency and a greater TNKS2 ARC4 stabilization in comparison to the dimethylamino-based substituent (**16** IC_50_ 31 µM). In addition, at R^3^ the piperazine (**S1** IC_50_ 250 µM), guanidine (**S4** IC_50_ 142 µM), methoxypropyl (**15** IC_50_ 368 µM), morpholine (**18** IC_50_ 85 µM, **23** IC_50_ 411 µM), pyridyl (**22** IC_50_ 325 µM) and phenyl (**25** IC_50_ 56 µM, **26** IC_50_ 30 µM) groups decrease potency towards TNKS2 ARC4. On the contrary, compound **S8**, characterized by a fully aliphatic R^3^ substituent, is the best tested analogue with an IC_50_ value of 8 µM and a 1.9-fold improved potency in comparison to **1**. Compounds **25** and **26** further indicate that the presence of tertiary amine at R^3^ is not necessary to achieve binding. Profiling of **S8** against TNKS1/2 ARCs indicates that the analogue maintains selectivity towards TNKS1/2 ARC4 (**Table 2**).

**Table 2.**
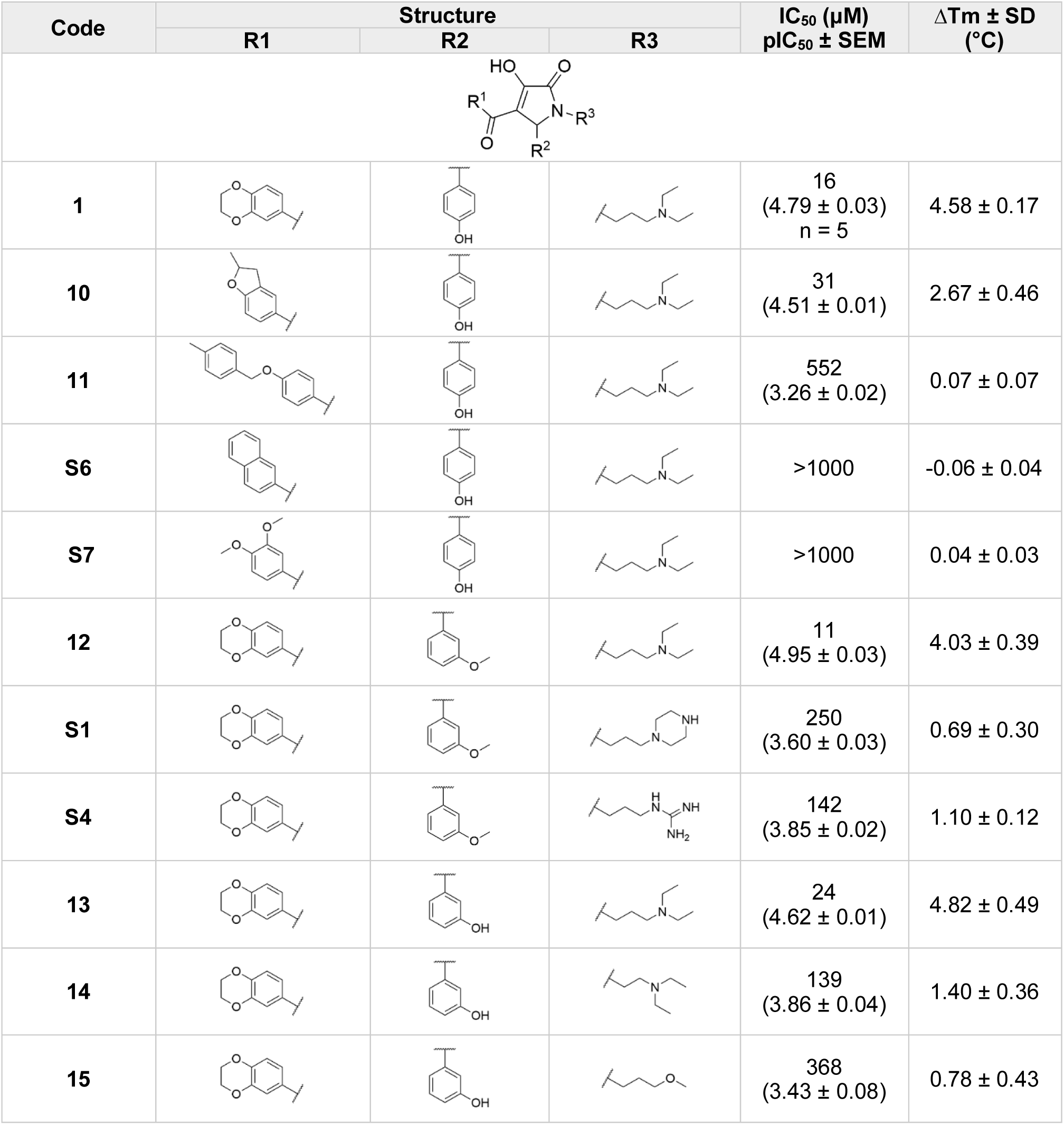
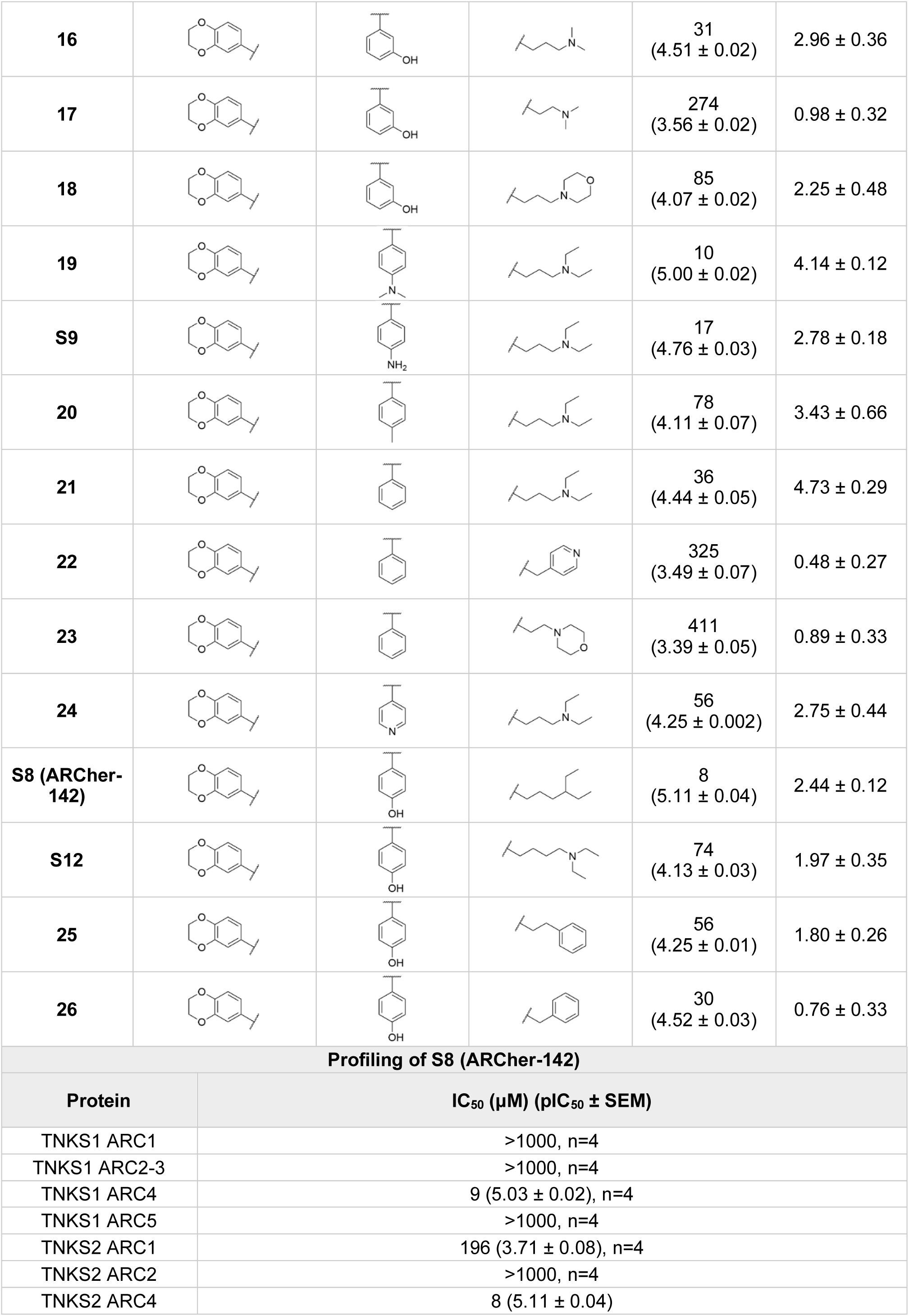
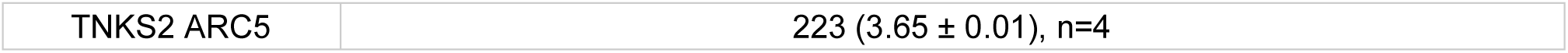
Structures of **1** and its derivatives for investigation of SARs and profiling of compound **S8** against TNKS1/2 ARCs. FRET-based potency measurements are indicated as IC_50_ (μM) and pIC_50_ ± SEM, with number of repetitions n=3 unless indicated otherwise (n=1 for measurements with IC_50_ >1000 μM unless differently indicated). Thermal stabilization (ΔTm ± SD) is indicated (n=4). The full list of all analyzed compounds is available in **Table S2**.

### Crystal structure of S8 in complex with TNKS2 ARC4 reveals its binding mode

To study the binding mode of the scaffold, we co-crystallized the compound displaying the best potency (**S8**) with TNKS2 ARC4 (**Figure 4**) (PDB: 9QFC). The crystal structure confirms that the compound binds to the TBM pocket and interestingly extends towards the hydrophobic sub-pocket, as indicated by our NMR titration experiments. The hydrophobic sub-pocket accommodates the 4-ethylhexan-1-yl substituent (R^3^) of the compound. Specifically, R^3^ forms hydrophobic contacts with F532 and L497, packing against the nonpolar component of R494, E498 and K501 side chains. The pyrrolone core is located between F532 and Y536 and it forms hydrogen bonds with K501 and Y536 side chains. In this way, the hydroxybenzene substituent (R^2^) packs against R525 and is directed towards the solvent, with the formation of a hydrogen bond with a water molecule. Below R^2^, the carbonyl group that connects the pyrrolone to the benzodioxane moiety (R^1^) forms an hydrogen bond with S527. This group also forms a hydrogen bond with a water molecule, which bridges a hydrogen bond interaction with D521. The residue F532 forms a π-π stacking interaction with the R^1^ benzene ring and directs R^1^ between Y536 and Y569, the tyrosine gate encasing the irreplaceable glycine of a TBM. In this way, the benzodioxane moiety is cozily surrounded by three aromatic ring side chains (F532, Y536, Y569). In addition, one of the oxygens of the dioxane ring could form a weak hydrogen bond with the hydroxyl group of Y536 (average distance 3.3 Å). The placement of the R^1^ moiety in the aromatic glycine sandwich explains the ability of the compound to inhibit the binding of the optimized peptide in the FRET assay, since the paramount glycine cannot be placed between the two tyrosine residues in the presence of the compound (**Figure 4C**).

**Figure 4.**
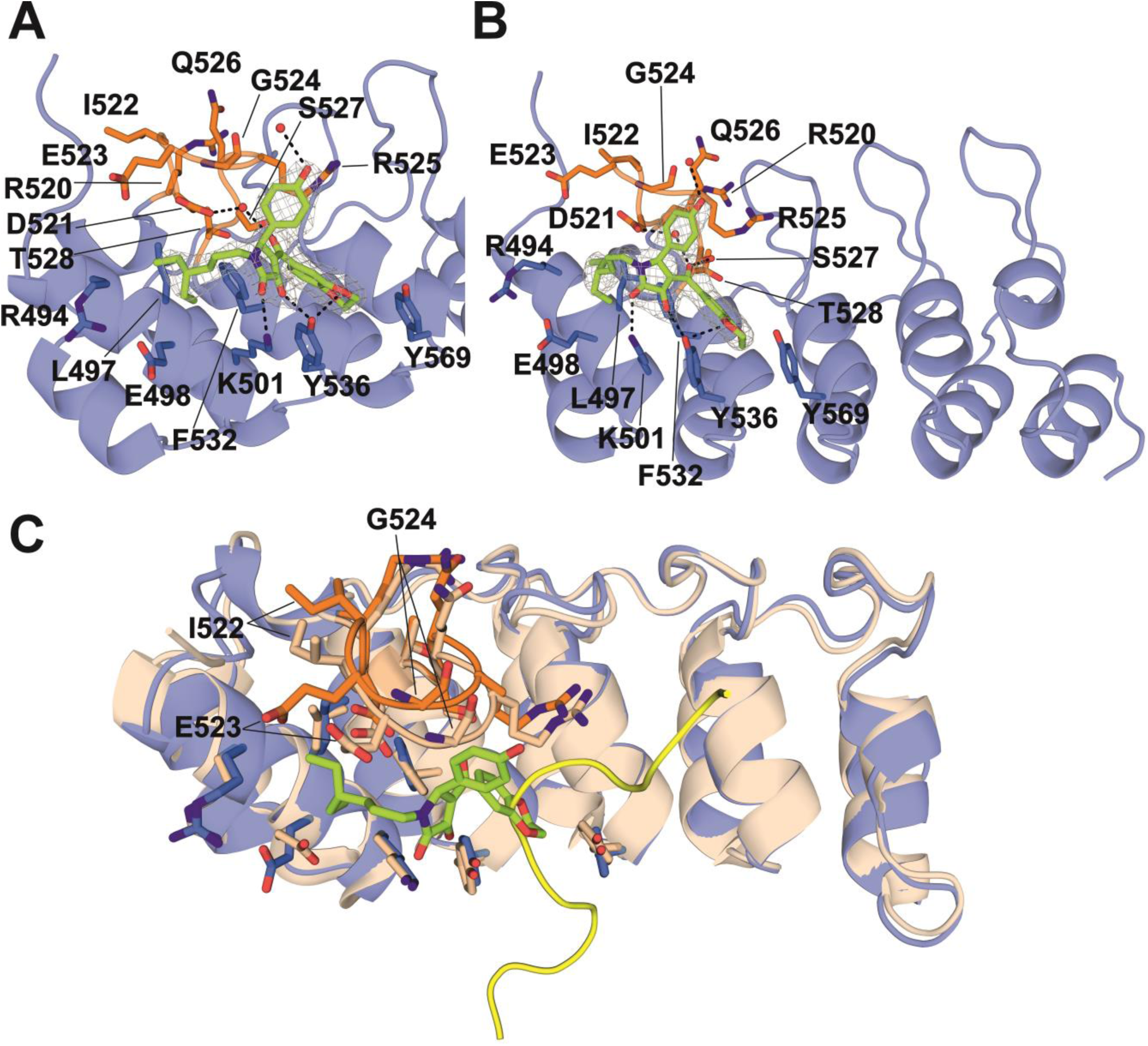
Crystal structure TNKS2 ARC4 in complex with S8. Panels (**A**) and (**B**) show two different perspectives of compound **S8** (green) in complex with TNKS2 ARC4 (blue) (PDB 9QFC). The mobile loop that packs against the compound is shown in orange. The sigma-A weighted 2Fo-Fc electron density map is grey and contoured at 1.0 σ. Hydrogen bonds are represented with black dashed lines. (**C**) Superimposed crystal structures of TNKS2 ARC4 in complex with 3BP2 TBM (PDB: 3TWR^3^, protein in color sand and TBM in yellow) and TNKS2 ARC4 in complex with **S8** (blue with mobile loop in orange). It is evident that the mobile loop, and especially I522, E523, G524, move to accommodate **S8**. The 3BP2 peptide is shown as cartoon and it overlaps with the compound in the binding pocket.

The crystal structure confirms the binding mode hypothesized with the our NMR experiments. Particularly evident is the perturbation of the loop formed by residues R520 to T528, against which the compound packs. Structural superimposition of our crystal structure with the crystal structure of TNKS2 ARC4 bound to the TBM of 3BP2 (PDB: 3TWR^3^) shows indeed that the loop moves to make space for the binding of **S8** (**Figure 4C**). The major displacement is observed for I522, whose α-carbon moves 2.3 Å, E523 (α-carbon movement of 2.6 Å) and G524 (α-carbon movement of 2.4 Å). This loop faces directly the R^2^ substituent of the compound, and its flexibility may explain the reason behind the tolerance towards different R^2^ moieties highlighted by our SAR analysis.

### S8 affects the WNT/β-catenin signaling pathway in cells

To test the effect of **S8** on the WNT/β-catenin pathway in cellular context, we employed a luciferase-based WNT/β-catenin reporter assay with the use of SuperTOP-Luciferase/Renilla HEK-293 cells.^59^ In the assay, the canonical WNT/β-catenin signaling events are activated by the presence of WNT3a in the cellular medium, leading to the binding of β-catenin to TCF/LEF transcription factors and to transcription of downstream genes of the WNT pathway. At the same time, firefly luciferase expression is increased, because it is under the control of a promoter with 7 TCF binding sites.^60^ Incubation of HEK293 cells with **S8** results in an attenuation of the WNT/β-catenin pathway with IC_50_ value of 45 ± 11 µM (n=3). This is in line with previous studies in which inhibition of ARCs leads to suppression of the WNT/β-catenin pathway.^48–50^

## Discussion

In this stufy, we employed a FRET-based high-throughput screening of the EU-OPENSCREEN Pilot and Commercials Diversity libraries (101004 compounds screened in total) to discover a new pyrrolone-based scaffold that specifically targets TNKS2 ARC4. The screening results are disclosed and available for consultation on ECBD.^53,54^ NMR analysis of the best hit compound **1** and its analogues **4** and **6** revealed a common binding site in TNKS2 ARC4. This binding site overlaps with the binding pocket of TBMs and extends towards a hydrophobic sub-pocket lined by a leucine residue (L497) and a phenylalanine (F532). The co-crystal structure of the analogue **S8** with TNKS2 ARC4 reveals that this site is indeed occupied by the aliphatic R^3^ substituent. SAR investigation indicates that a 3-carbon chain is ideal to achieve optimal potency, while longer or shorter aliphatic chains are detrimental. The positive charge provided by the tertiary amine present at R^3^ of most compounds is beneficial, as it could form an ion-pair interaction with E498, but it is not necessary to achieve high potency. The co-crystal structure also highlights the importance of the benzodioxane moiety (R^1^), as confirmed by the SAR investigation. Notably, F532 guides the double ring system inside the tyrosine gate (the “aromatic glycine sandwich”) of the peptide binding pocket, inhibiting the interaction of the optimized peptide. The phenylalanine residue at this position is present only in ARC4 and ARC1 domains, where ARC1 domains are devoid of the tyrosine gate and cannot form the hydrogen bond that instead anchors the pyrrolone core to ARC4, as evident from the crystal structure. Several substituents on the benzene ring at R^2^ are tolerated, since this moiety packs against a flexible loop, variable across ARCs, and is directed towards the solvent. Compound **S8** is the best analogue from our SARs analysis: it maintains selectivity towards TNKS1/2 ARC4 and it displays a potency of 8 μM against TNKS2 ARC4 with optimal protein stabilization in the DSF assay. The compound attenuates the WNT/β-catenin signaling in cells, indicating that the compound can enter the cellular environment and affect the tankyrase-modulated signaling pathway despite only affecting one ARC. At present, tankyrase partners able to bind exclusively to ARC4 have not been discovered, but ARC-selective inhibitors such as **S8** can provide the tools to identify specific proteins and relative pathways controlled by different ARCs. The development of ARC inhibitors further allows the investigation of the interplay between tankyrase scaffolding and catalytic functions, and it may offer an alternative therapeutic approach to catalytic inhibition.

## Materials and Methods

### Cloning

Cloning was performed as previously described.^52^ The method employed to clone the expression constructs into vectors was the one-step sequence– and ligation-independent cloning (SLIC).^61^ The employed expression vectors (pNIC-CFP, pNIC-YFP, pNIC-MBP) originated from the pNIC28-Bsa4 plasmid (26103, Addgene). Specifically, mCerulean (CFP, 29726, Addgene plasmid), mCitrine (YFP, 29724, Addgene plasmid) and *E. coli* maltose-binding protein (Uniprot P0AEX9, residues 27-392) sequences were inserted between the 6xHis-tag and the TEV protease site of the pNIC28-Bsa4 plasmid. Purified PCR products of the constructs were mixed with linearized plasmids (100 ng) in 1:3 molar ratio in presence of T4 polymerase (M0203L, New England Biolabs). The mix was then kept at room temperature for 2.5 minutes and incubated on ice for 10 minutes. After incubation, the reactions were transformed into NEB® 5-alpha Competent *E. coli* (High Efficiency) cells (C2987H, New England BioLabs) according to the high efficiency transformation protocol provided by the manufacturer.^62,63^ Colonies were grown overnight at 37 °C on LB agar with 50 μg/mL kanamycin (KAN0025, Formedium Ltd.) and 5% sucrose for *sacB* counter-selection.^64^ TNKS2 SAM mutants (Y920A and E897K) were obtained through DpnI mediated site-directed mutagenesis.^52,65^ The sequences of the inserted regions were verified through sequencing.

### Protein expression

Expression of YFP and CFP fused proteins was performed as described before.^52^ Plasmids were transformed into BL21(DE3) Competent *E. coli* cells (C2527H, New England BioLabs) according to the manufacturer’s instructions.^66^ Colonies were grown overnight at 37 °C on LB agar supplemented with kanamycin (50 μg/mL) and 5% sucrose. The day after, single colonies were selected for overnight growth of pre-cultures in Terrific Broth (TB, TRB0102, Formedium Ltd.) media with 50 μg/mL kanamycin. Pre-cultures were then added to TB autoinduction media including trace elements (AIMTB0210, Formedium Ltd) with 8 g/L glycerol and 50 μg/ml kanamycin, or LB media with the same antibiotic (for ^15^N TNKS2 ARC4). The cultures were grown at 37 °C in shaking conditions until OD_600_= 1. At this point, the procedure for the expression of the isotopically labelled ^15^N TNKS2 ARC4 required additional steps, including washing the cells with M9 medium without labelled nitrogen sources, centrifugation (5000 x g), resuspension of the pellet in M9 medium with ^15^N source and incubation to allow recovery of cell growth.^67^ Expression of ^15^N TNKS2 ARC4 was induced by addition of 1 mM isopropyl β-D-thiogalactopyranoside (IPTG, IPTG025, Formedium Ltd.). After cell growth, the temperature of the cell cultures was set to 18 °C (17 °C for ^15^N TNKS2 ARC4) for overnight incubation. The morning after, cells were harvested by centrifugation at 4200 × g for 30 minutes at 4°C, and they were resuspended in lysis buffer (50 mM HEPES pH 7.5, 500 mM NaCl, 10% volume/volume glycerol, 0.5 mM TCEP) in the presence of proteinase k inhibitor (Pefabloc® SC, 11585916001, Roche) and DNase I (10104159001, Roche). Resuspended cells were frozen in liquid nitrogen and stored at –20 °C until purification.

### Protein Purification

Purification of YFP and CFP fused proteins was carried out as previously described.^52^ Cells were thawed, lysed by sonication, centrifugated at 16000 x g for 30 minutes and filtered through a 0.45 µm filter. All the proteins underwent a first step of purification through immobilized metal affinity chromatography (IMAC) on Ni^2+^ loaded HiTrap Chelating HP column (17040901, Cytiva). MBP-tagged TNKS2 ARC4 was purified with an additional step of MBP-based affinity chromatography by using MBP-Trap HP column (28918779, Cytiva). TNKS2 ARC4 and ^15^N TNKS2 ARC4 (MBP-tagged) were dialyzed against size exclusion chromatography (SEC) buffer (30 mM HEPES pH 7.5, 350 mM NaCl, 10% volume/volume glycerol, 0.5 mM TCEP) in presence of 6xHis-tagged TEV protease (1:30 molar ratio, 20 hours, 4 °C) to cleave MBP. TNKS2 ARC4 and ^15^N TNKS2 ARC4 were also purified through reverse-MBP chromatography and reverse-IMAC to remove TEV and MBP contaminants. All proteins were further purified through SEC using a HiLoad 16/600 Superdex 75 pg (28989333, Cytiva) and SEC buffer (20 mM HEPES pH 7.5, 100 mM NaCl, 0.5 mM TCEP for TNKS2 ARC4 purification). Fractions containing pure ^15^N TNKS2 ARC4 were pooled together, concentrated and buffer exchanged to the final buffer (20 mM NaPhosphate pH 7.0, 50 mM NaCl). All proteins were aliquoted, frozen in liquid nitrogen and stored at –70 °C. Mass spectrometry analysis of ^15^N TNKS2 ARC4 indicated a labelling efficiency of 92.6%.

### FRET-based high-throughput screening and dose response validation

The FRET assay for discovery of chemical probes targeting ARC and SAM domains of tankyrases was developed in a previous study.^52^ The high-throughput screening of the EU-OPENSCREEN Pilot and Commercials Diversity libraries was performed at the Norwegian Centre for Molecular Biosciences and Medicine (NCMBM, formerly NCMM). For the primary screen, assay-ready plates containing library compound sufficient to reach a screening concentration of 30 µM were prepared on black shallow-well 384-microwell plates (Greiner bio-one small volume, high base, non-binding microplate black, 784900 or Fisherbrand™ 384-Well ShallowWell Polypropylene Microplates, 12-566-617, Fisher Scientific) using 30 nL of 10 mM DMSO compound stocks dispensed with an Echo 550 liquid handler (Labcyte). Each screening plate also contained DMSO controls at equivalent volumes to compounds for positive, negative, and assay buffer blank controls, 16 wells per control type. Stock solutions of CFP-TNKS2 ARC4 and YFP-REAGDGEE or CFP-SAM(E897K) and YFP-SAM(Y920A) were combined and diluted in assay buffer (10 mM BTP pH 7, 3% weight/volume PEG 20000, 0.01% volume/volume Triton, 0.5 mM TCEP) to final concentrations of 100 nM and 200 nM or 150 and 300 nM, respectively. The positive control was prepared by addition of guanidine hydrochloride to the above assay buffer to a final concentration of 1M. A volume of 10 µL of these solutions was added to each well of the assay-ready plates using a BioTek MultiFlo FX dispenser (Agilent) equipped with 5 uL dispensing cassettes for the peristaltic pumps (screening solution and control) and a syringe pump (assay buffer blank). After liquid handling, plates were incubated at RT for 30 minutes before reading fluorescence using a BioTek Synergy Neo2 plate reader (Agilent) with Gen5 software (version 3.02) at four wavelength combinations – 410/477, 410/527, 430/477, 430/527 – each with an excitation bandwidth of 20 nm and an emission bandwidth of 10 nm; a gain setting of 135; a “Normal” read speed with 10 msec delay after plate movement, 50 measurements per data point, low lamp energy, and a “Standard” dynamic range (0 to 99,999); and a read height of 4.5 mm.

Analysis proceeded using a custom workflow created in KNIME version 4.3.4. All plates had to pass a quality filter of Z’ ≥ 0.5 and SSMD ≤ –5. These quality metrics were calculated using the means and standard deviations of the control groups on each plate, as follows: Z’=1-(3×(σ_Positive+σ_Negative))/(|μ_Positive – μ_Negative|), SSMD=(μ_Positive – μ_Negative)/√(〖 σ_Positive〗^2+〖σ_Negative〗^2). The buffer blank was subtracted from each emission measurement and the rFRET was calculated by dividing the fluorescence intensity at 527 nm by the fluorescence intensity at 477 nm for both excitation at 410 nm and 430 nm to give r410 and r430, respectively. These values were then normalized on a plate-wise basis, setting the response of the positive control (screening solution with 1M guanidine hydrochloride and DMSO) to 100% and the response of the negative control (screening solution with DMSO) to 0%. Both donor emission and the r410 value were used to filter fluorescently interfering compounds. Any compounds with donor emission falling outside of 150-50% of the mean of donor emission for the controls on the same plate were flagged as interfering. Any compounds where r410 had a percent difference from r430 greater than 50%, calculated as 100%*(|r430-r410|/0.5*(r430+r410)), were also flagged as interfering. In the early screen this filter was originally a percentage point difference between normalized r430 and r410, excluding compounds where |r410%-r430%| > 50% – changing the filter led to a small amount of previously excluded compounds being reclassified as hits, but these were not rescreened.

For the dose response validation, assay-ready plates containing library compound sufficient to reach concentrations of 0.5, 1.25, 2.5, 5, 10, 25, 50, and 100 µM were prepared on black shallow-well 384-microwell plates (Greiner bio-one small volume, high base, non-binding microplate black, 784900 or Fisherbrand™ 384-Well ShallowWell Polypropylene Microplates, 12-566-617, Fisher Scientific) using 10 mM DMSO compound stocks dispensed with an Echo 550 (Labcyte). Each compound was tested at these concentrations in duplicate. Compound-containing wells were backfilled with DMSO to a total volume of 100 nL, and on each plate 16 wells for each of the positive, negative, and assay blank controls also received 100 nL DMSO. Screening solutions were added to the assay ready plates and fluorescence read as detailed above for the primary screen.

Dose response data were filtered to remove compounds displaying fluorescence interference and normalized as detailed above. Compounds that had at least four doses which passed the fluorescence filtering were further analyzed for dose response. Within KNIME, we used a custom R script calling the function *drm* from the package ‘drc’ with fct=LL.4 to fit the normalized data to a four-parameter log-logistic curve. This function returned the curve upper limit, lower limit, slope and **IC_50_** as parameters. From these parameters and the curve function, derived responses at the maximum and minimum doses used in the curve fit were used to calculate an activity change by subtraction of the latter from the former. For each compound, if the activity change was ≥ 25% and if the **IC_50_** value fell within the tested concentrations that passed the interference filters, then that compound was considered a validated hit.

### FRET-based assay IC_50_ measurements

To measure the potency of compounds against ARC inhibitors, CFP-TNKS1/2 ARCs (100 nM) and YFP-REAGDGEE (200 nM) were mixed with different concentrations of the compounds (100 µM to 3 nM) in FRET buffer (10 mM Bis-Tris Propane pH 7, 3% weight/volume PEG 20000, 0.01% volume/volume Triton X-100, 0.5 mM TCEP). The control for absence of interaction was represented by the FRET pair in absence of compound, while the control for total inhibition was the FRET pair in FRET Inhibiting buffer (10 mM Bis-Tris Propane pH 7, 3% weight/volume PEG 20000, 0.01% volume/volume Triton X-100, 0.5 mM TCEP, 1 M guanidine hydrochloride). The controls were set to be 2 log units apart from the lowest and highest concentrations of compounds tested, respectively. Compounds were dispensed using Echo 650T liquid handler (Labcyte), while protein mixtures and buffer were dispensed using a Formulatrix Mantis liquid dispenser. Each condition was studied with at least four replicates in 384-well plates (Fisherbrand™ 384-Well ShallowWell Polypropylene Microplates, 12-566-617, Fisher Scientific) with a final volume of 10 µl per well. Measurements were carried out by employing 430 nm excitation wavelength and emission wavelengths of 477 nm and 527 nm. Fluorescence readings were performed with the use of a Tecan Spark multimode microplate reader. The blank was subtracted from each emission measurement and the rFRET was calculated by dividing the fluorescence intensity at 527 nm by the fluorescence intensity at 477 nm. Potency values were calculated with GraphPad Prism 10 using a nonlinear regression analysis (sigmoidal dose–response fitting with variable slope).

### Differential scanning fluorimetry assay

This assay was performed by incubating 5 μM TNKS2 ARC4 with 100 μM of compound in buffer (10 mM Bis-Tris-Propane pH 7.0, 3% weight/volume PEG 20000, 0.01% volume/volume Triton X-100, 0.5 mM TCEP) with the presence 5X SYPRO™ Orange dye (S6650, Invitrogen). Three controls were employed: TNKS2 ARC4 with REAGDGEE peptide (100 μM) as positive control; TNKS2 ARC4 with 1% DMSO, which is the same amount of DMSO present when incubating the protein with 100 μM compound (stored in DMSO); TNKS2 ARC4, to measure the melting temperature of the protein only. Each condition was studied in four replicates of 20 μL and measured in a sealed Multiplate™ 96-Well PCR plate (MLL9601, Bio-Rad) with the use of a CFX96 Real-Time PCR Detection System (Bio-Rad). During the experiment, the temperature was gradually raised from 20°C to 95°C (1°C increase per minute) with measurements performed at intervals of 1 minute. After the experiment, data were normalized and analyzed with GraphPad Prism 10 by using a nonlinear regression analysis (Boltzmann sigmoid equation).

### NMR titration experiments and analysis of NMR data

All NMR data for the human TNKS2 ARC4 domain were measured using the Bruker Avance III HD NMR spectrometer, operating at 850 MHz of ^1^H frequency and equipped with a 5 mm ^1^H/^13^C/^15^N triple-resonance TCI cryoprobe. Mapping of the inhibitor binding site on the TNKS2 ARC4 domain was carried out by measuring two-dimensional ^1^H, ^15^N correlation spectra (^15^N-HSQC) of 0.2 mM ^15^N-labeled TNKS2 ARC4 in 20 mM sodium phosphate buffer pH 7, 5% DMSO, 50 mM NaCl without inhibitor and in the presence 0.25, 0.5, 1, 2, 4, 8 and 16 molar excess of unlabelled compounds **4** and **6**. For compound **1**, the inhibitor: protein ratios of 1:4, 1:2, 1:1, 2:1, 4:1 and 12:1 were used. Given that ligand stocks were dissolved in DMSO, the titration series were prepared in such a way that at each titration point the NMR sample contained 0.2 mM ARC domain and the desired molar ratio of inhibitor in 5% DMSO to eliminate chemical shift perturbations stemming from DMSO-buffer mixture. The ^15^N-HSQC data were collected at 20 °C using 128 and 1024 complex points in ^15^N (F_1_) and ^1^H (F_2_) dimensions, respectively. This corresponds to acquisition time of 48 and 75 ms in F_1_ and F_2_ dimensions, respectively. Spectra were collected using 4-64 scans per FID with the repetition rate of 1.1 seconds. Resolution in F_1_ dimension was improved by linear prediction and zero-filled 4x prior to Fourier transform. Spectra were processed using Topspin 3.5 pl7 software package (Bruker). NMR data were analyzed with CcpNmr Analysis Version 3 software.^68^ Full backbone and partial side chain assignment of the human protein TNKS2 ARC4 has been previously reported by Zaleska et al.^69^ (BMRB Entry 27747). This assignment allowed us to identify backbone peaks, which were assigned manually in all spectra of the titration experiments. Chemical shift perturbations were calculated for each residue backbone signals as Euclidean distances, with weighting of ^15^N signals by a factor of 0.14, between the positions at compound saturation and in the control spectrum (DMSO only). Residues that displayed CSPs higher than 1σ or 2σ (σ being the standard deviation for all residues CSPs) were relevant for the determination of the compounds binding sites.^70^ To identify differences between the binding sites of the compounds, Euclidean distances between the residues peaks at saturation (ΔCSP) were calculated with weighting of ^15^N signals by a factor of 0.14. ΔCSP values higher than 0.1 or 2 ppm were considered important for differential binding of the compounds.

### Co-crystallization, data collection, processing and refinement

TNKS2 ARC4 domain (residues 487-649, 8 mg/mL) was incubated with 2 mM **S8** in ice for 1 hour. Protein crystals were grown at 4°C by mixing the protein and compound mixture in a 3:1 ratio (protein: precipitant solution) with 0.1 M phosphate/citrate buffer pH 4.2, 40% volume/volume PEG 300. The crystallization was carried out with the sitting-drop vapor diffusion method in SWISSCI 96-well 3-Lens crystallization plate. Long needles of protein crystals were fully grown after five days. The crystals were cryo-protected by soaking into precipitant solution added with 20% volume/volume glycerol and then frozen in liquid nitrogen. Data were collected at Diamond (beamline I24). Diffraction data were processed using XDS package.^71^ Molecular replacement was performed with PHASER^72^ by using TNKS2 ARC4 structure (PDB: 3TWR^3^) as model. Protein model building was carried out with Coot, while REFMAC5 and Phenix were employed for refinement of the crystal structure.^73–75^ In the crystal structure, each copy of TNKS2 ARC4 is bound to two **S8** molecules, one in the pocket for binding TBMs (*S* enantiomer) and one at crystal contacts (*R* enantiomer). Data collection and refinement statistics for the crystal structure is provided in **Table S3**.

### Cell Culture

The human cell line HEK-293 (CRL-1573, from embryo’s kidney) was obtained from the American Type Culture Collection (ATCC). HEK 293 cells were cultured in high-glucose DMEM (Dulbecco′s Modified Eagle′s Medium, D6429, Sigma-Aldrich). The medium contained 10% Fetal Bovine Serum (FBS, 10270106, Gibco) and 1% Penicillin-Streptomycin (P4333, Sigma-Aldrich). Cells were cultured at 37 °C in a humidified cell incubator with 5% CO_2_. The cell culture was kept below 20 passages (∼10 weeks) and routinely monitored (upon thawing and monthly) for Mycoplasma infections with MycoAlert® Mycoplasma Detection Kit (Lonza). The cell line was authenticated by short tandem repeat profiling to confirm identity (Eurofins).

### Transfection of HEK-293 cells

A stable HEK-293 cell line containing the SuperTOP-Flash plasmid (ST-Luc HEK-293) provided with a 7×TCF binding sites promoter was kindly provided by Dr. Vladimir Korinek. To generate SuperTOP-Luciferase/Renilla HEK-293 cells (ST-Luc/Ren HEK-293), the pRL-TK (Renilla, E2241, Promega) cassette was subcloned into pPUR (Promega), creating the pRL-TK-puro construct. The linearized pRL-TK-puro was transfected into ST-Luc HEK-293 cells using FuGENE® 6 Transfection Reagent (E2691, Promega) before selection with 2.5 μg/mL Puromycin (P9620, Sigma-Aldrich).

### Wnt3a-Conditioned Medium (Wnt3a-CM)

The Wnt3a-expressing cells were purchased from ATCC (male mouse, CRL-2647) and maintained according to the supplier’s recommendations, under the same conditions as HEK-293 cells. Wnt3a-CM was collected following ATCC’s instructions.^76^

### WNT/β-catenin signaling pathway reporter cell assay

A total of 40000 ST-Luc/Renilla HEK-293 cells were seeded in 96-well plates coated with poly-L-lysine (sc-286689, Santa Cruz Biotechnology). The day after, the cells were incubated for a 24 hours treatment with increasing concentrations of **S8** compound in 50% Wnt3a-CM. After compound exposure, the cells were lysed, and firefly luciferase and Renilla activities were measured using a GloMax® Multi Detection System (E7031, Promega) with the Dual-Glo Luciferase Assay System (E1980, Promega).

### Chemistry: General methods

Commercially available reagents were used without further purification and all solvents were of HPLC grade quality. Anhydrous THF and DMF were obtained using PureSolve MD-7 solvent purification system. All reactions were monitored by thin-layer chromatography (TLC) using Merck aluminum sheets precoated with 0.25 mm silica gel, 60 F_254_ plates and/or by reversed-phase ultra-high performance liquid chromatography mass spectrometry (RP-UPLC-MS). Analytical RP-UPLC-MS (ESI) analysis was performed on a S2 Waters AQUITY RP-UPLC system equipped with a diode array detector using an Thermo Accucore C18 column (d 2.6 μm, 2.1 x 50 mm; column temp: 50 °C; flow: 1.0 mL/min). Eluents A (H_2_O + 0.1% formic acid) and B (ACN + 0.1% formic acid) were used in a linear gradient (5% B to 100% B) in 2.4 min and then held for 0.1 min at 100% B (total run time: 2.6 min). The LC system was coupled to a SQD mass spectrometer. Flash column chromatography was performed using Merck Geduran® Si 60 (40–63 µm) silica gel. All new compounds were characterized by ^1^H NMR, ^13^C NMR, HRMS (ESI) and melting point. Preparative HPLC analyses were performed using a Shimadzu LC-20 HPLC system comprised of an LC-20AP pump, an SPD-M20A PDA array detector and a SIL-10AP autosampler (injection volume: 3500 µL). Chromatographic separations were achieved on a Shimadzu column (C18, 10 µm, 20 x 250 mm) placed in a CTO-20AC column oven. Eluents A (H_2_O with 0.1% formic acid) and B (acetonitrile with 0.1% formic acid) were used in a linear gradient (5% B to 60% B) in 23 min, up to 100% B in 2 min, held until 27 min and then column was re-equilibrated at 5% B for 3 min (total run time: 30 min), at a flow rate of 20 mL/min. NMR spectra were acquired at 298 K using either a 400 MHz Bruker AVANCE III HD spectrometer equipped with a Prodigy CryoProbe, a 600 MHz Bruker AVANCE III spectrometer equipped with a Bruker BBFO SmartProbe or an 800 MHz Bruker AVANCE III HD spectrometer equipped with a TCI CryoProbe; chemical shifts (δ) are reported in parts per million (ppm) relative to tetramethylsilane (TMS) as internal standard and coupling constants (*J*) are expressed in hertz (Hz). Melting points were obtained using a Stuart SMP30 melting point apparatus and are uncorrected. HRMS was performed using an Agilent Infinity II 1290 UHPLC system (Agilent Technologies, Santa Clara, CA, USA) for injection directly into the MS. About 90% of the flow was diverted off using a static splitter. MS detection was performed in positive mode on an Agilent 6545 QTOF MS equipped with Agilent Dual Jet Stream electrospray ion source with a drying gas temperature of 200 °C, gas flow of 8 L/min, nebulizer pressure of 35 psi, sheath gas temperature of 200 °C, sheath gas flow of 11 L/min, a fragmentor value of 175 V, skimmer voltage of 65 V and octapole 1 frequency of 750 Vpp. The mass spectra were acquired in the 100–3000 m/z mass range at a rate of 1 spectrum per second, collecting 5890 transients per spectrum. Constant reference mass correction was applied using all the masses defined in the instrument supplier-provided solution (ESI-L Low Concentration Tuning Mix from Agilent Technologies, prod. no. G1969-85000) delivered from the instrument’s reference mass bottle directly to the secondary nebulizer of the ion source. The data was analyzed using the qualitative analysis of Mass Hunter B.07.00. Purity of compounds as determined by HPLC-MS, ^1^H and ^13^C NMR (**Figure S3**-**S32**) was higher than 95%.

### General procedure for the synthesis of 2,4-diketoesters 2a–c

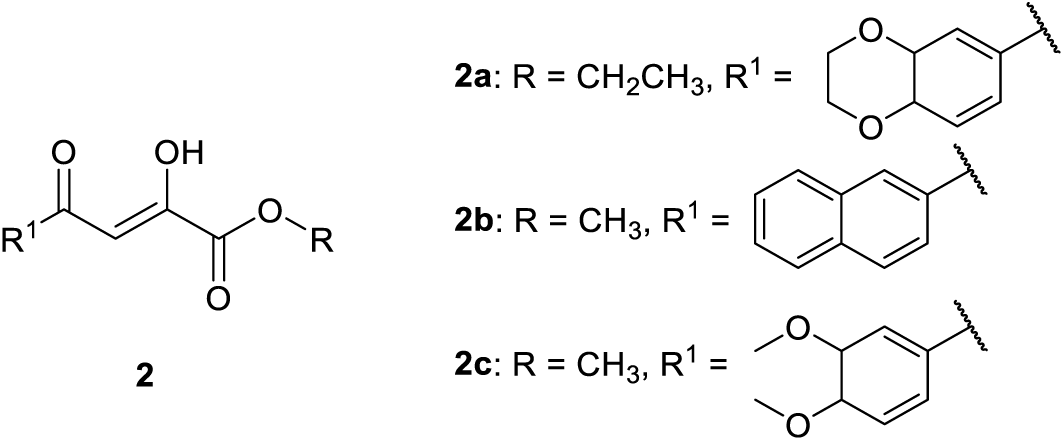

2,4-diketo esters **2a–c** were synthesized *via* a Claisen condensation reaction between an aryl methyl ketone and a dialkyl oxalate, by modifying a previously published method.^56^ Aryl ketone (5.6 mmol, 1.0 equiv.) and dialkyl oxalate (9.5 mmol, 1.7 equiv.) were dissolved in 3 mL of anhydrous THF. This solution was added dropwise to a stirred suspension of 60% sodium hydride (11.2 mmol, 2.0 equiv.) in 6 mL of anhydrous THF at 0 °C. Upon completion of the addition, the ice bath was removed, and the reaction mixture was heated to reflux for 3 h. THF was then removed under reduced pressure and 10 mL of iced water, 6 mL of EtOAc and 560 µL of conc. H_2_SO_4_ were added to the residue. The mixture was stirred and transferred to a separatory funnel. Organic and aqueous phases were separated, and the aqueous layer was extracted with EtOAc (3 x 20 mL). The combined organic phases were washed with brine, dried over anhydrous Na_2_SO_4_ and concentrated under reduced pressure. 2,4-diketo esters were purified by flash column chromatography (gradient elution, 20–70% EtOAc / *n*-heptane) yielding the pure intermediates. ^1^H and ^13^C NMR spectra were in good agreement with the experimental data reported in the literature.^77–79^

### General procedure for the synthesis of pyrrolones

**Method A:** Adapted from a previously published method ^57^. To a stirred solution of the appropriate aldehyde (0.323 mmol, 1.0 equiv.) in dry THF (1 mL) under N_2_, 3-(piperazin-1-yl)propan-1-amine (0.323 mmol, 1.0 equiv.) was added dropwise. After 15 min, a solution of the 2,4-diketo ester **2a** (0.323 mmol, 1.0 equiv.) in THF (2 mL) was added and the mixture was stirred overnight at room temperature. The solvent was removed under vacuum and the resulting crude product was purified by preparative RP-HPLC.

**Method B:** Adapted from a previously published method ^58^. To a solution of anhydrous sodium acetate (0.288 mmol, 1.0 equiv.) in glacial acetic acid (2 mL) were added the 2,4-diketo ester **2a** (0.288 mmol, 1.0 equiv.), the appropriate aldehyde and *N*-(3-aminopropyl)guanidine hydrochloride (0.288 mmol, 1.0 equiv.). The mixture was heated to reflux to ensure complete dissolution of the starting materials and then, it was allowed to cool down to room temperature and stirred overnight. Toluene was added to the mixture and the solvent was removed under reduced pressure. The resulting crude product was purified by preparative RP-HPLC.

**Method C:** To a stirred solution of the appropriate aldehyde (0.390 mmol, 1.0 equiv.) in EtOH (2 mL) and catalytic glacial acetic acid (≈ 5 drops), the appropriate amine (0.390 mmol, 1.0 equiv.) was added dropwise. After 10 min, a suspension of the 2,4-diketo ester (0.390 mmol, 1.0 equiv.) in EtOH (2 mL) was added. The mixture was heated to reflux to ensure complete dissolution of the starting materials and then, it was allowed to cool down to room temperature and stirred overnight. Then, toluene was added to the mixture and the solvent was removed under vacuum. The resulting crude product was purified by preparative RP-HPLC.

### Compound S1

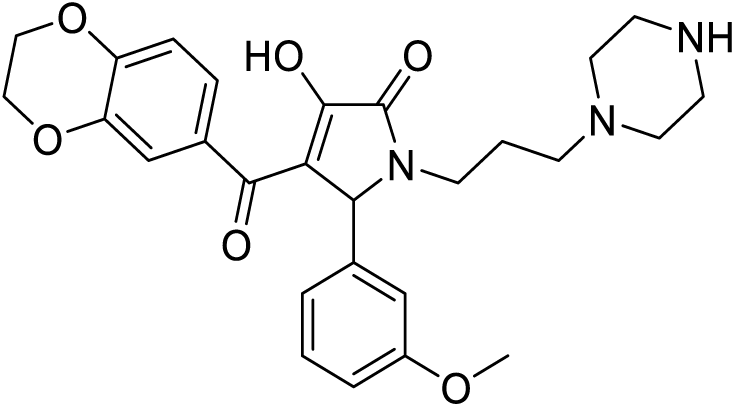

**Method A** was followed using 3-methoxybenzaldehyde, 3-(piperazin-1-yl)propan-1-amine and 2,4-diketo ester **2a**. The crude product was purified by preparative RP-HPLC (H_2_O + 0.1% HCOOH / ACN + 0.1% HCOOH) using linear gradient elution (5–60% ACN (0.1% HCOOH), to afford **S1** as a yellow solid (Yield: 18%); m.p.: 139–141 °C; ^1^H NMR (800 MHz, (CD_3_)_2_SO) δ 8.19 (s, 1H), 7.50 (d, *J* = 2.0 Hz, 1H), 7.35 (dd, *J* = 8.3, 2.0 Hz, 1H), 7.16 (t, *J* = 7.9 Hz, 1H), 6.84–6.89 (m, 2H), 6.77–6.73 (m, 1H), 6.72 (d, *J* = 8.4 Hz, 1H), 5.17 (s, 1H), 4.27–4.16 (m, 4H), 3.69 (s, 3H), 3.54–3.47 (m, 1H), 3.00 (bs, 4H), 2.73–2.63 (m, 1H), 2.42 (bs, 4H), 2.25–2.20 (m, 1H), 2.20–2.16 (m, 1H), 1.58–1.52 (m, 1H), 1.50–1.44 (m, 1H); ^13^C NMR (200 MHz, (CD_3_)_2_SO) δ 183.28, 170.33, 165.34, 163.54, 158.89, 144.82, 143.50, 141.77, 134.45, 128.82, 122.34, 119.69, 118.31, 115.23, 113.68, 111.75, 64.26, 63.87, 60.73, 55.00, 54.87, 49.55, 43.03, 38.19, 24.45; HRMS (ESI): *m/z* [M + H]^+^ calcd for C_27_H_32_N_3_O_6_: 494.2286; found: 494.2285.

### Compound S2

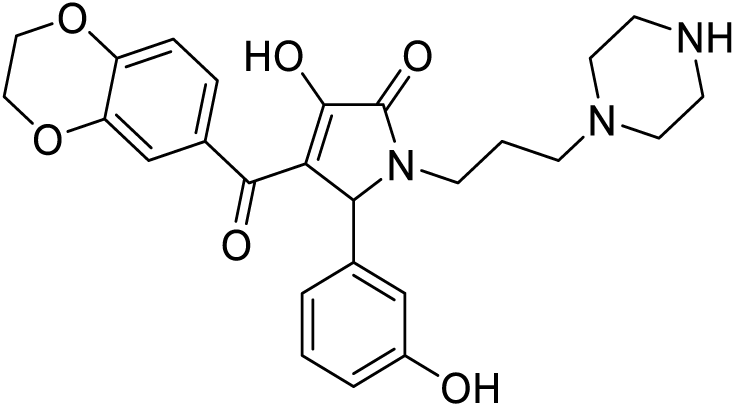

Compound **S2** was synthesized following **Method A** using 3-hydroxybenzaldehyde, 3-(piperazin-1-yl)propan-1-amine and 2,4-diketo ester **2a**. The crude product was purified by preparative RP-HPLC (H_2_O + 0.1% HCOOH / ACN + 0.1% HCOOH) using linear gradient elution (5–60% ACN (0.1% HCOOH), to afford **S2** as a yellow solid (Yield: 11%); m.p.: 195–197 °C; ^1^H NMR (600 MHz, (CD_3_)_2_SO) δ 8.18 (s, 1H), 7.44 (dd, *J* = 15.4, 2.0 Hz, 1H), 7.32 (td, *J* = 8.3, 2.0 Hz, 1H), 7.07–7.02 (m, 1H), 6.78–6.73 (m, 1H), 6.69 (d, *J* = 7.5 Hz, 1H), 6.65 (s, 1H), 6.57 (dd, *J* = 7.9, 2.6 Hz, 1H), 5.17 (s, 1H), 4.24–4.16 (m, 4H), 3.54–3.50 (m, 1H), 3.00 (bs, 4H), 2.70–2.62 (m, 1H), 2.44 (bs, 4H), 2.29–2.18 (m, 2H), 1.68–1.53 (m, 1H), 1.53–1.42 (m, 1H); ^13^C NMR (150 MHz, (CD_3_)_2_SO) δ 184.31, 169.56, 163.56, 160.77, 157.16, 145.42, 142.16, 142.00, 133.85, 128.89, 122.50, 118.55, 118.45, 115.59, 114.28, 114.20, 64.38, 63.94, 60.87, 54.90, 49.34, 42.98, 38.28, 24.46; HRMS (ESI): *m/z* [M + H]^+^ calcd for C_26_H_30_N_3_O_6_: 480.2129; found: 480.2128.

### Compound S3

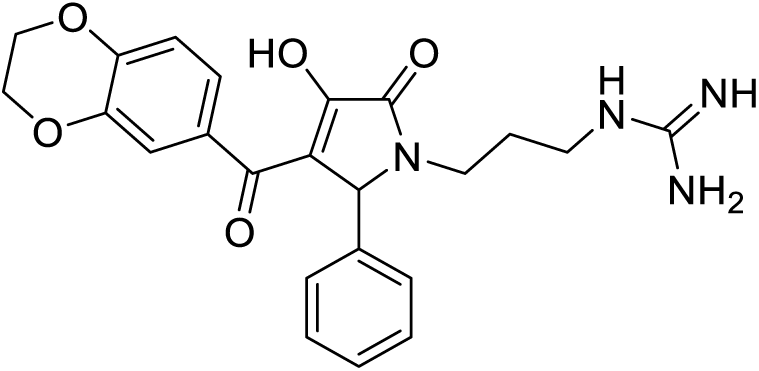

Compound **S3** was synthesized following **Method B** using benzaldehyde, N-(3-aminopropyl)guanidine hydrochloride and 2,4-diketo ester **2a**. The crude product was purified by preparative RP-HPLC (H_2_O + 0.1% HCOOH / ACN + 0.1% HCOOH) using linear gradient elution (5–60% ACN (0.1% HCOOH), to afford **S3** as a pale yellow solid (Yield: 10%); m.p.: 237–239 °C; ^1^H NMR (800 MHz, (CD_3_)_2_SO) δ 7.94 (s, 1H), 7.51–7.15 (m, 7H), 6.69 (d, *J* = 8.4 Hz, 1H), 5.20 (s, 1H), 4.30–4.12 (m, 4H), 3.55–3.44 (m, 1H), 3.04–2.99 (m, 2H), 2.62–2.56 (m, 1H), 1.67–1.51 (m, 2H); ^13^C NMR (200 MHz, (CD_3_)_2_SO) δ 182.94, 170.93, 165.94, 163.53, 156.80, 144.66, 141.72, 134.62, 127.78, 127.53, 126.75, 122.32, 118.19, 115.13, 111.59, 64.23, 63.86, 60.91, 38.34, 37.58, 27.01; HRMS (ESI): *m/z* [M + H]^+^ calcd for C_23_H_25_N_4_O_5_: 437.1819; found: 437.1827.

### Compound S4

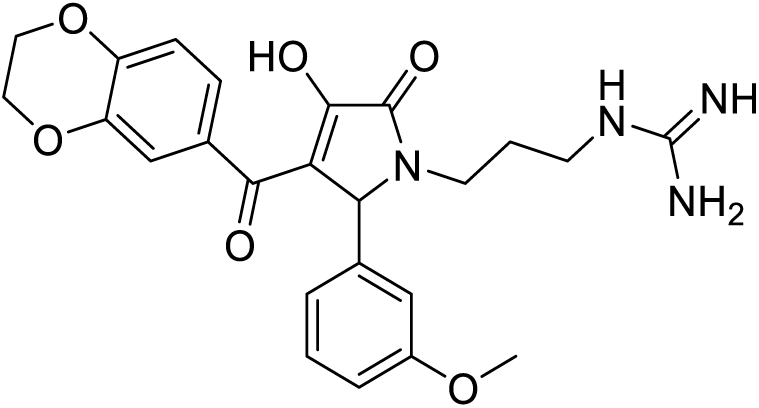

Compound **S4** was synthesized following **Method B** using 3-methoxybenzaldehyde, N-(3-aminopropyl)guanidine hydrochloride and 2,4-diketo ester **2a**. The crude product was purified by preparative RP-HPLC (H_2_O + 0.1% HCOOH / ACN + 0.1% HCOOH) using linear gradient elution (5–60% ACN (0.1% HCOOH), to afford **S4** as a pale orange solid (Yield: 11%); m.p.: 197–199 °C; ^1^H NMR (400 MHz, (CD_3_OD) δ 8.25 (s, 2H), 7.32–7.27 (m, 2H), 7.21 (t, *J* = 7.9 Hz, 1H), 6.89–6.84 (m, 2H), 6.82–6.76 (m, 2H), 5.40 (s, 1H), 4.30–4.18 (m, 4H), 3.74 (s, 3H), 3.66–3.57 (m, 1H), 3.18–3.02 (m, 2H), 3.00–2.91 (m, 1H), 1.74–1.61 (m, 2H); ^13^C NMR (100 MHz, (CD_3_OD) δ 189.63, 170.40, 166.62, 161.37, 158.59, 148.61, 144.31, 140.60, 133.78, 130.75, 124.12, 120.97, 119.54, 118.11, 117.53, 114.67, 114.62, 65.94, 65.42, 63.26, 55.69, 39.82, 39.39, 28.44; HRMS (ESI): *m/z* [M + H]^+^ calcd for C_24_H_27_N_4_O_6_: 467.1925; found: 467.1933.

### Compound S5

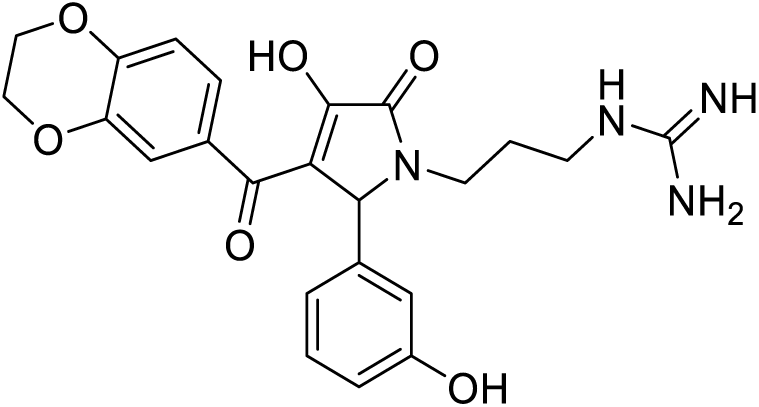

Compound **S5** was synthesized following **Method B** using 3-hydroxybenzaldehyde, *N*-(3-aminopropyl)guanidine hydrochloride and 2,4-diketo ester **2a**. The crude product was purified by preparative RP-HPLC (H_2_O + 0.1% HCOOH / ACN + 0.1% HCOOH) using linear gradient elution (5–60% ACN (0.1% HCOOH), to afford **SCALL0122** as a pale orange solid (Yield: 17%); m.p.: 187-189 °C; ^1^H NMR (400 MHz, CD_3_OD) δ 8.27 (s, 2H), 7.31–7.29 (m, 2H), 7.11 (t, *J* = 7.8 Hz, 1H), 6.81–6.77 (m, 2H), 6.73–6.64 (m, 2H), 5.37 (s, 1H), 4.30–4.20 (m, 4H), 3.63–3.59 (m, 1H), 3.13–3.08 (m, 2H), 3.00–2.96 (m, 1H), 1.69–1.65 (m, 2H); ^13^C NMR (100 MHz, CD_3_OD) δ 189.65, 170.85, 167.44, 167.42, 158.82, 158.59, 148.40, 144.25, 140.92, 134.06, 130.62, 124.08, 120.23, 119.57, 117.44, 116.20, 115.37, 65.93, 65.42, 63.18, 39.83, 39.34, 28.43; HRMS (ESI): *m/z* [M + H]^+^ calcd for C_23_H_25_N_4_O_6_: 453.1769; found: 453.1778.

### Compound S6

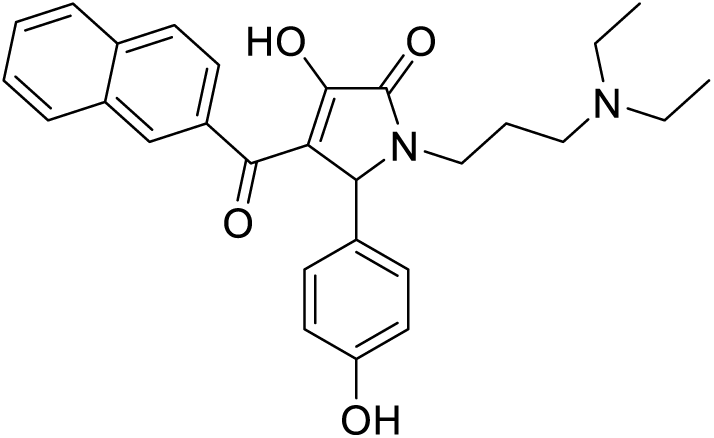

Compound **S6** was synthesized following **Method C** using 4-hydroxybenzaldehyde, *N,N-*diethyl-3-diaminopropane and 2,4-diketo ester **2b**. The crude product was purified by preparative RP-HPLC (H_2_O + 0.1% HCOOH / ACN + 0.1% HCOOH) using linear gradient elution (5–60% ACN (0.1% HCOOH), to afford **S6** as a pale yellow solid (Yield: 22%); m.p.: 213–214 °C; ^1^H NMR (400 MHz, (CD_3_)_2_SO) δ 9.26 (s, 1H), 8.40 (s, 1H), 7.90 (dd, *J* = 18.2, 7.7 Hz, 2H), 7.77 (q, *J* = 8.4 Hz, 2H), 7.58–7.41 (m, 2H), 7.14 (d, *J* = 8.0 Hz, 2H), 6.68 (d, *J* = 7.9 Hz, 2H), 5.23 (s, 1H), 3.46–3.39 (m, 1H), 3.01–2.89 (m, 4H), 2.87–2.75 (m, 3H), 1.84–1.58 (m, 2H), 1.05 (t, *J* = 7.3 Hz, 6H); ^13^C NMR (100 MHz, (CD_3_)_2_SO) δ 185.08, 179.74, 169.84, 156.53, 138.42, 133.79, 132.04, 130.31, 128.89, 128.82, 128.65, 127.35, 126.63, 126.60, 126.27, 126.05, 125.72, 114.81, 60.52, 49.04, 45.84, 37.81, 21.78, 8.70; HRMS (ESI): *m/z* [M + H]^+^ calcd for C_28_H_31_N_2_O_4_: 459.2278; found: 459.2285.

### Compound S7

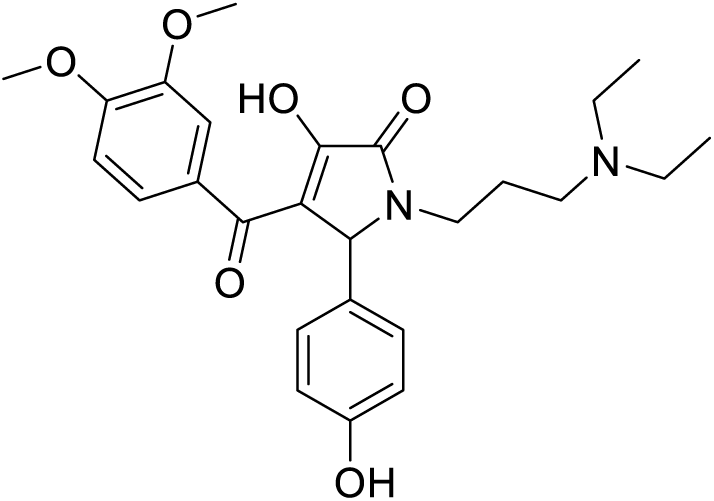

Compound **S7** was synthesized following **Method C** using 4-hydroxybenzaldehyde, *N,N-*diethyl-3-diaminopropane and 2,4-diketo ester **2c**. The crude product was purified by preparative RP-HPLC (H_2_O + 0.1% HCOOH / ACN + 0.1% HCOOH) using linear gradient elution (5–60% ACN (0.1% HCOOH), to afford **S7** as a pale yellow solid (Yield: 12%); m.p.: 243–245 °C; ^1^H NMR (400 MHz, (CD_3_)_2_SO) δ 9.24 (s, 1H), 7.59 (bs, 2H), 7.06 (d, *J* = 8.0 Hz, 2H), 6.86 (d, *J* = 8.7 Hz, 1H), 6.65 (d, *J* = 8.0 Hz, 2H), 5.17 (s, 1H), 3.76 (s, 1H), 3.73 (s, 1H), 3.47–3.43 (m, 1H), 2.91 (q, *J* = 7.3 Hz, 4H), 2.79–2.70 (m, 3H), 1.69–1.64 (m, 2H), 1.09 (t, *J* = 7.1 Hz, 6H); ^13^C NMR (100 MHz, (CD_3_)_2_SO) δ 184.25, 169.75, 156.47, 150.65, 147.27, 133.15, 130.84, 128.62, 122.83, 114.74, 112.71, 109.90, 60.69, 56.02, 55.48, 55.35, 49.05, 45.83, 37.83, 22.09, 18.57, 9.00; HRMS (ESI): *m/z* [M + H]^+^ calcd for C_26_H_33_N_2_O_6_: 469.2333; found: 469.2342.

### Compound S8

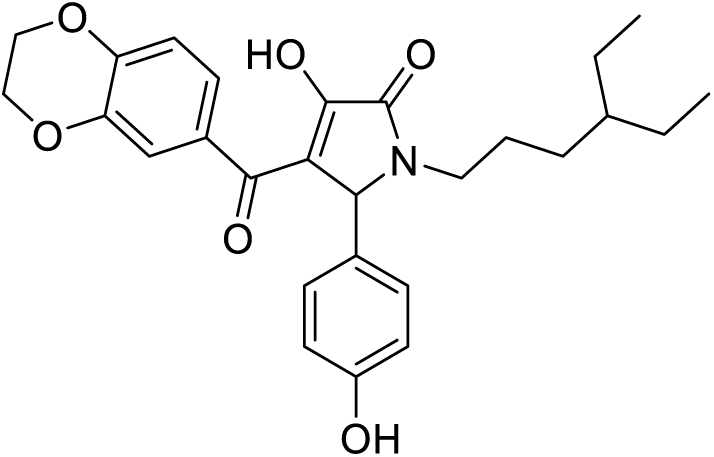

Compound **S8** was synthesized following **Method C** using 4-hydroxybenzaldehyde, 4-ethylhexan-1-amine and 2,4-diketo ester **2a**. The crude product was purified by preparative RP-HPLC (H_2_O + 0.1% HCOOH / ACN + 0.1% HCOOH) using linear gradient elution (5–60% ACN (0.1% HCOOH), to afford **S8** as a beige solid (Yield: 3%); m.p.: 203–205 °C; ^1^H NMR (400 MHz, CD_3_OD) δ 7.33 (dd, *J* = 8.5, 2.1 Hz, 1H), 7.28 (d, *J* = 2.0 Hz, 1H), 7.06 (d, *J* = 8.5 Hz, 2H), 6.84 (d, *J* = 8.4 Hz, 1H), 6.71 (d, *J* = 8.5 Hz, 2H), 5.44 (s, 1H), 4.32–4.20 (m, 4H), 3.61–3.46 (m, 1H), 2.93–2.79 (m, 1H), 1.55–1.44 (m, 1H), 1.41–1.30 (m, 1H), 1.28–1.21 (m, 4H), 1.20–1.08 (m, 3H), 0.82 (td, *J* = 7.4, 1.4 Hz, 6H); ^13^C NMR (100 MHz, CD_3_OD) δ 190.04, 166.90, 159.01, 153.55, 149.73, 144.63, 132.27, 130.12, 126.93, 124.46, 121.53, 119.59, 117.92, 116.63, 66.04, 65.40, 63.12, 42.34, 41.25, 30.77, 26.41, 26.29, 11.22, 11.13; HRMS (ESI): *m/z* [M + H]^+^ calcd for C_27_H_32_NO_6_: 466.2224; found: 466.2232.

### Compound S9

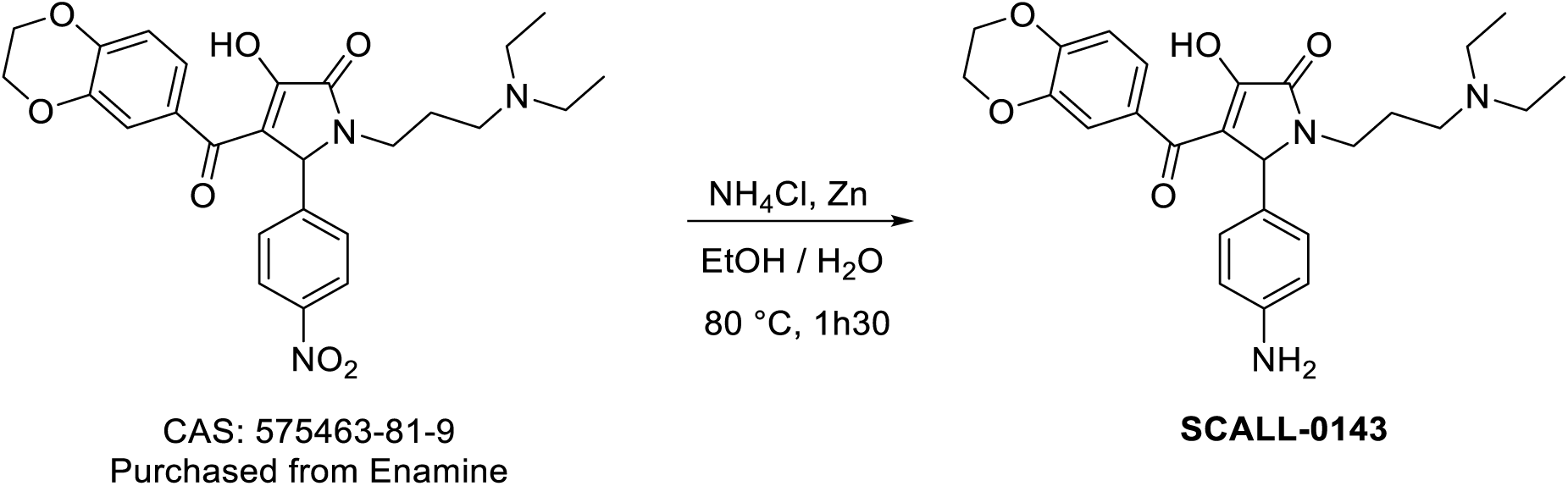

Compound **S9** was synthesized following a modified version of a previously reported procedure ^80^. To a solution of the commercially available 1-(3-(diethylamino)propyl)-4-(2,3-dihydrobenzo[b][1,4]dioxine-6-carbonyl)-3-hydroxy-5-(4-nitrophenyl)-1h-pyrrol-2(5h)-one (0.121 mmol, 1equiv.) in EtOH/H_2_O (1 mL/320 µL) were added ammonium chloride (0.158 equiv., 1.3 equiv.) and zinc powder (0.968, 8 equiv.). The mixture was stirred and heated to 80°C for 1h30. After cooling to room temperature, solvent was removed under reduced pressure, the residue was dispersed in MeOH and zinc powder was removed by filtration. The crude product was then purified by preparative RP-HPLC (H_2_O + 0.1% HCOOH / ACN + 0.1% HCOOH) using linear gradient elution (5–60% ACN (0.1% HCOOH), to afford **S9** as a yellow solid (Yield: 12%); m.p.: 193–196 °C; ^1^H NMR (800 MHz, (CD_3_)_2_SO) δ 8.20 (s, 1H), 7.46 (s, 1H), 7.33 (dd, *J* = 8.3, 2.0 Hz, 1H), 6.86 (d, *J* = 8.2 Hz, 2H), 6.70 (d, *J* = 8.4 Hz, 1H), 6.42 (d, *J* = 8.5 Hz, 2H), 5.32 (t, *J* = 4.8 Hz, NH_2_), 5.03 (s, 1H), 4.31–4.17 (m, 4H), 3.41–3.39 (m, 1H), 2.65–2.60 (m, 1H), 2.43–2.34 (m, 2H), 2.04–1.93 (m, 1H), 1.60–1.54 (m, 1H), 1.49–1.41 (m, 1H), 1.30–1.20 (m, 3H), 0.94 (t, *J* = 7.2 Hz, 6H); ^13^C NMR (200 MHz, (CD_3_)_2_SO) δ 174.65, 147.90, 145.00, 142.21, 135.34, 130.07, 128.68, 128.46, 122.74, 118.80, 115.67, 113.86, 64.66, 64.34, 60.90, 50.06, 46.51, 38.35, 35.65, 29.11, 27.13, 25.67, 24.82, 11.34; HRMS (ESI): *m/z* [M + H]^+^ calcd for C_26_H_32_N_3_O_5_: 466.2336; found: 466.2344.

### Compound S10

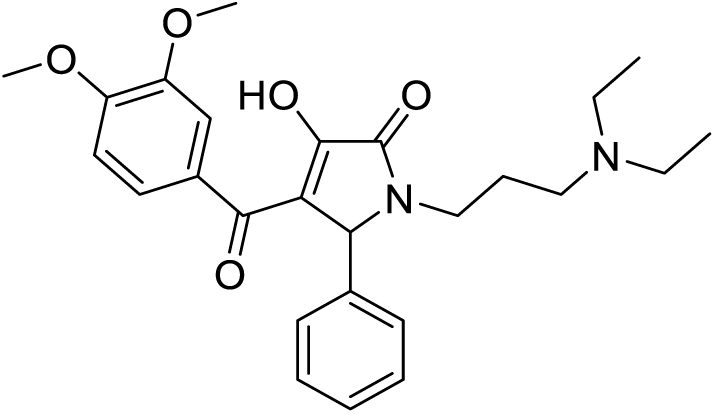

Compound **S10** was synthesized following **Method C** using benzaldehyde, *N,N-*diethyl-3-diaminopropane and 2,4-diketo ester **2c**. The crude product was purified by preparative RP-HPLC (H_2_O + 0.1% HCOOH / ACN + 0.1% HCOOH) using linear gradient elution (5–60% ACN (0.1% HCOOH), to afford **S10** as a pale yellow solid (Yield: 17%); m.p.: 203–205 °C; ^1^H NMR (400 MHz, (CD_3_)_2_SO) δ 7.67–7.56 (m, 2H), 7.30–7.24 (m, 4H), 7.23–7.15 (m, 1H), 6.85 (d, *J* = 8.8 Hz, 1H), 5.27 (s, 1H), 3.76 (s, 3H), 3.73 (s, 3H), 3.48–3.41 (m, 1H), 2.95 (q, *J* = 7.2 Hz 4H), 2.87–2.77 (m, 2H), 2.75–2.63 (m, 1H), 1.80–1.60 (m, 2H), 1.10 (t, *J* = 7.2 Hz, 6H); ^13^C NMR (100 MHz, (CD_3_)_2_SO) δ 184.12, 170.17, 163.61, 150.63, 147.25, 141.17, 133.11, 127.97, 127.64, 127.03, 122.79, 112.88, 112.74, 109.88, 61.19, 55.48, 55.35, 49.00, 45.84, 38.01, 21.85, 8.81; HRMS (ESI): *m/z* [M + H]^+^ calcd for C_26_H_33_N_2_O_5_: 453.2384; found: 453.2392.

### Compound S11

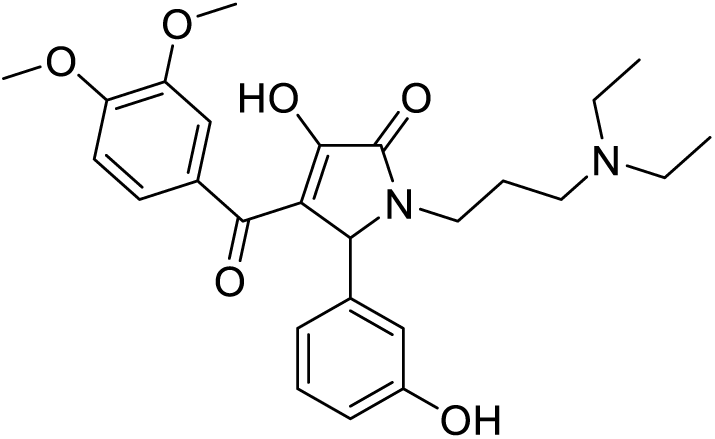

Compound **S11** was synthesized following **Method C** using 3-hydroxybenzaldehyde, *N,N-*diethyl-3-diaminopropane and 2,4-diketo ester **2c**. The crude product was purified by preparative RP-HPLC (H_2_O + 0.1% HCOOH / ACN + 0.1% HCOOH) using linear gradient elution (5–60 % ACN (0.1% HCOOH), to afford **S11** as a yellow solid (Yield: 15%); m.p.: 196–198 °C; ^1^H NMR (400 MHz, (CD_3_)_2_SO) δ 9.30 (s, 1H), 7.66–7.57 (m, 2H), 7.06 (t, *J* = 7.8 Hz, 1H), 6.86 (d, *J* = 8.2 Hz, 1H), 6.74 (d, *J* = 7.9 Hz, 1H), 6.71–6.66 (m, 1H), 6.59 (dd, *J* = 8.0, 2.4 Hz, 1H), 5.18 (s, 1H), 3.76 (s, 3H), 3.73 (s, 3H), 3.47–3.41 (m, 1H), 2.95 (q, *J* = 7.2 Hz, 4H), 2.88–2.80 (m, 2H), 2.79–2.71 (m, 1H), 1.79–1.58 (m, 2H), 1.11 (t, *J* = 7.2 Hz, 6H); ^13^C NMR (100 MHz, (CD_3_)_2_SO) δ 184.60, 70.66, 164.98, 157.39, 150.86, 147.51, 143.00, 133.48, 129.24, 123.12, 119.11, 114.51, 114.37, 113.02, 112.76, 110.22, 61.38, 55.79, 55.67, 49.23, 46.38, 38.24, 22.23, 9.01; HRMS (ESI): *m/z* [M + H]^+^ calcd for C_26_H_33_N_2_O_6_: 469.2333; found: 469.2340.

### Compound S12

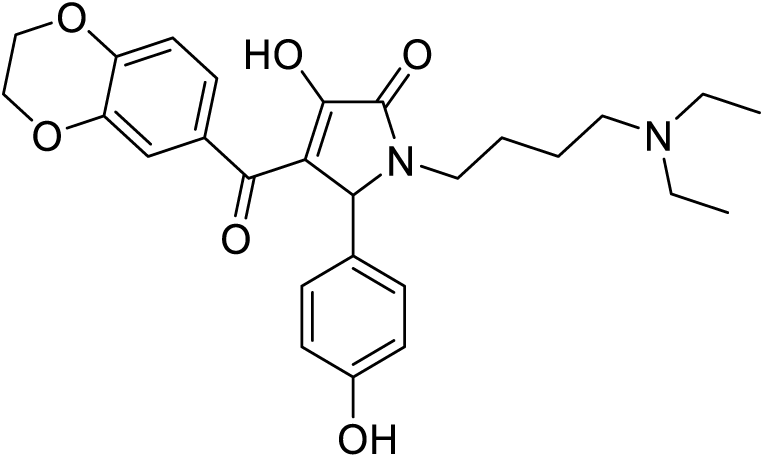

Compound **S12** was synthesized following **Method C** using 4-hydroxybenzaldehyde, *N,N*-diethylbutane-1,4-diamine and 2,4-diketo ester **2a**. The crude product was purified by preparative RP-HPLC (H_2_O + 0.1% HCOOH / ACN + 0.1% HCOOH) using linear gradient elution (5–60% ACN (0.1% HCOOH), to afford **S12** as a yellow solid (Yield: 7%); m.p.: 124–126 °C; ^1^H NMR (400 MHz, CD_3_OD) δ 8.27 (s, 1H), 7.36–7.26 (m, 2H), 7.11 (d, *J* = 8.2 Hz, 2H), 6.84 – 6.77 (m, 1H), 6.71 (d, *J* = 8.2 Hz, 2H), 5.37 (s, 1H), 4.29 – 4.20 (m, 4H), 3.66 – 3.53 (m, 1H), 3.12 (q, *J* = 7.2 Hz, 4H), 3.03 (t, *J* = 7.5 Hz, 2H), 2.98–2.89 (m, 1H), 1.64–1.42 (m, 4H), 1.25 (t, *J* = 7.2 Hz, 6H); ^13^C NMR (100 MHz, CD_3_OD) δ 189.67, 169.76, 166.83, 158.66, 148.71, 144.33, 133.68, 130.12, 129.14, 124.14, 119.63, 118.50, 117.59, 116.45, 65.97, 65.44, 62.65, 52.54, 48.13, 40.87, 26.15, 22.24, 9.03; HRMS (ESI): *m/z* [M + H]^+^ calcd for C_27_H_33_N_2_O_6_: 481.2333; found: 481.2341.

## Data availability

The coordinates and structure factors of the crystal structure are deposited to PDB with ID 9QFC. Raw diffraction data and NMR data will be available at www.fairdata.fi (https://doi.org/10.23729/0dfe3539-a7b9-4a60-ad72-a698576de760).

## Acknowledgements

The use of the facilities and expertise of the Biocenter Oulu Structural Biology core facility (a member of Biocenter Finland, Instruct-ERIC Centre Finland and FINStruct), Proteomics and Protein Analysis core facility (a member of Biocenter Finland) and Biocenter Oulu sequencing center are gratefully acknowledged. We gratefully acknowledge the staff at the Chemical Biology Platform at the Norwegian Centre for Molecular Biosciences and Medicine (formerly NCMM), especially Eirin Solberg, for performing the high-throughput screen. This work benefited from access to the Instruct-ERIC Centre Finland NMR facility at the University of Helsinki. We greatly thank Perttu Permi and Tuomas Niemi-Aro for the help with NMR data collection. Financial support was provided by Instruct-ERIC (PID 23611). We are grateful to local contacts at Diamond for providing assistance in using beamline I24.

## Author contributions

Conceptualization, Ch.B., S.T.S., L.L.; methodology, Ch.B.; investigation, Ch.B., A.G.P., A.G., Cl.B., S.A.B., F.N., V.P.; data curation, Ch.B., A.G.P., A.G., J.P.; formal analysis, Ch.B., A.G.P., A.G., Cl.B., S.A.B., J.P., V.P.; writing-original draft, Ch.B.; writing-review & editing, Ch.B., A.G.P., S.T.S., A.G., Cl.B., S.A.B., F.N., J.P., V.P., J.W., M.H.C., L.L.; visualization, Ch.B.; validation, Ch.B.; funding acquisition, L.L.; resources, L.L; supervision, L.L., J.W., M.H.C.

## Supplemental information

**Document S1**. Figure S1-S32 and Table S1-S3.

## References

1. Sedgwick SG, Smerdon SJ. The ankyrin repeat: a diversity of interactions on a common structural framework. Trends in Biochemical Sciences. 1999;24(8):311–316. doi:10.1016/S0968-0004(99)01426-7

2. Li J, Mahajan A, Tsai MD. Ankyrin Repeat: A Unique Motif Mediating Protein−Protein Interactions^†^. Biochemistry. 2006;45(51):15168–15178. doi:10.1021/bi062188q

3. Guettler S, LaRose J, Petsalaki E, et al. Structural Basis and Sequence Rules for Substrate Recognition by Tankyrase Explain the Basis for Cherubism Disease. Cell. 2011;147(6):1340–1354. doi:10.1016/j.cell.2011.10.046

4. Mariotti L, Templeton CM, Ranes M, et al. Tankyrase Requires SAM Domain-Dependent Polymerization to Support Wnt-β-Catenin Signaling. Molecular Cell. 2016;63(3):498–513. doi:10.1016/j.molcel.2016.06.019

5. DaRosa PA, Ovchinnikov S, Xu W, Klevit RE. Structural insights into SAM domain-mediated tankyrase oligomerization: Structural Insights into Tankyrase SAM Oligomerization. Protein Science. 2016;25(9):1744–1752. doi:10.1002/pro.2968

6. De Rycker M, Price CM. Tankyrase Polymerization Is Controlled by Its Sterile Alpha Motif and Poly(ADP-Ribose) Polymerase Domains. Molecular and Cellular Biology. 2004;24(22):9802–9812. doi:10.1128/MCB.24.22.9802-9812.2004

7. Riccio AA, McCauley M, Langelier MF, Pascal JM. Tankyrase Sterile α Motif Domain Polymerization Is Required for Its Role in Wnt Signaling. Structure. 2016;24(9):1573–1581. doi:10.1016/j.str.2016.06.022

8. Callow MG, Tran H, Phu L, et al. Ubiquitin Ligase RNF146 Regulates Tankyrase and Axin to Promote Wnt Signaling. Hotchin NA, ed. PLoS ONE. 2011;6(7):e22595. doi:10.1371/journal.pone.0022595

9. Kang HC, Lee YI, Shin JH, et al. Iduna is a poly(ADP-ribose) (PAR)-dependent E3 ubiquitin ligase that regulates DNA damage. Proc Natl Acad Sci USA. 2011;108(34):14103–14108. doi:10.1073/pnas.1108799108

10. Zhang Y, Liu S, Mickanin C, et al. RNF146 is a poly(ADP-ribose)-directed E3 ligase that regulates axin degradation and Wnt signalling. Nat Cell Biol. 2011;13(5):623–629. doi:10.1038/ncb2222

11. Wang Z, Michaud GA, Cheng Z, et al. Recognition of the *iso* –ADP-ribose moiety in poly(ADP-ribose) by WWE domains suggests a general mechanism for poly(ADP-ribosyl)ation-dependent ubiquitination. Genes Dev. 2012;26(3):235–240. doi:10.1101/gad.182618.111

12. Zhou Z dong, Chan CH shan, Xiao Z cheng, Tan EK. Ring finger protein 146/Iduna is a Poly(ADP-ribose) polymer binding and PARsylation dependent E3 ubiquitin ligase. Cell Adhesion & Migration. 2011;5(6):463–471. doi:10.4161/cam.5.6.18356

13. Sowa ST, Bosetti C, Galera-Prat A, Johnson MS, Lehtiö L. An Evolutionary Perspective on the Origin, Conservation and Binding Partner Acquisition of Tankyrases. Biomolecules. 2022;12(11):1688. doi:10.3390/biom12111688

14. Morrone S, Cheng Z, Moon RT, Cong F, Xu W. Crystal structure of a Tankyrase-Axin complex and its implications for Axin turnover and Tankyrase substrate recruitment. Proc Natl Acad Sci USA. 2012;109(5):1500–1505. doi:10.1073/pnas.1116618109

15. DaRosa PA, Klevit RE, Xu W. Structural basis for tankyrase-RNF146 interaction reveals noncanonical tankyrase-binding motifs. Protein Science. 2018;27(6):1057–1067. doi:10.1002/pro.3413

16. Eisemann T, McCauley M, Langelier MF, et al. Tankyrase-1 Ankyrin Repeats Form an Adaptable Binding Platform for Targets of ADP-Ribose Modification. Structure. 2016;24(10):1679–1692. doi:10.1016/j.str.2016.07.014

17. Koirala S, Klein J, Zheng Y, et al. Tissue-Specific Regulation of the Wnt/β-Catenin Pathway by PAGE4 Inhibition of Tankyrase. Cell Reports. 2020;32(3):107922. doi:10.1016/j.celrep.2020.107922

18. Xu D, Liu J, Fu T, et al. USP25 regulates Wnt signaling by controlling the stability of tankyrases. Genes Dev. 2017;31(10):1024–1035. doi:10.1101/gad.300889.117

19. Li X, Han H, Zhou MT, et al. Proteomic Analysis of the Human Tankyrase Protein Interaction Network Reveals Its Role in Pexophagy. Cell Reports. 2017;20(3):737–749. doi:10.1016/j.celrep.2017.06.077

20. Bhardwaj A, Yang Y, Ueberheide B, Smith S. Whole proteome analysis of human tankyrase knockout cells reveals targets of tankyrase-mediated degradation. Nat Commun. 2017;8(1):2214. doi:10.1038/s41467-017-02363-w

21. Liu K, Yu C, Xie M, Li K, Ding S. Chemical Modulation of Cell Fate in Stem Cell Therapeutics and Regenerative Medicine. Cell Chemical Biology. 2016;23(8):893–916. doi:10.1016/j.chembiol.2016.07.007

22. Huang SMA, Mishina YM, Liu S, et al. Tankyrase inhibition stabilizes axin and antagonizes Wnt signalling. Nature. 2009;461(7264):614–620. doi:10.1038/nature08356

23. Liu J, Xiao Q, Xiao J, et al. Wnt/β-catenin signalling: function, biological mechanisms, and therapeutic opportunities. Sig Transduct Target Ther. 2022;7(1):3. doi:10.1038/s41392-021-00762-6

24. Hayat R, Manzoor M, Hussain A. Wnt signaling pathway: A comprehensive review. Cell Biology International. 2022;46(6):863–877. doi:10.1002/cbin.11797

25. Zhan T, Rindtorff N, Boutros M. Wnt signaling in cancer. Oncogene. 2017;36(11):1461–1473. doi:10.1038/onc.2016.304

26. Hu HH, Cao G, Wu XQ, Vaziri ND, Zhao YY. Wnt signaling pathway in aging-related tissue fibrosis and therapies. Ageing Research Reviews. 2020;60:101063. doi:10.1016/j.arr.2020.101063

27. Mariotti L, Pollock K, Guettler S. Regulation of Wnt/β-catenin signalling by tankyrase-dependent poly(ADP-ribosyl)ation and scaffolding. British J Pharmacology. 2017;174(24):4611–4636. doi:10.1111/bph.14038

28. Verma A, Kumar A, Chugh A, Kumar S, Kumar P. Tankyrase inhibitors: emerging and promising therapeutics for cancer treatment. Med Chem Res. 2021;30(1):50–73. doi:10.1007/s00044-020-02657-7

29. Wang W, Li N, Li X, Tran MK, Han X, Chen J. Tankyrase Inhibitors Target YAP by Stabilizing Angiomotin Family Proteins. Cell Reports. 2015;13(3):524–532. doi:10.1016/j.celrep.2015.09.014

30. Wang H, Lu B, Castillo J, et al. Tankyrase Inhibitor Sensitizes Lung Cancer Cells to Endothelial Growth Factor Receptor (EGFR) Inhibition via Stabilizing Angiomotins and Inhibiting YAP Signaling. Journal of Biological Chemistry. 2016;291(29):15256–15266. doi:10.1074/jbc.M116.722967

31. Jia J, Qiao Y, Pilo MG, et al. Tankyrase inhibitors suppress hepatocellular carcinoma cell growth via modulating the Hippo cascade. Avila MA, ed. PLoS ONE. 2017;12(9):e0184068. doi:10.1371/journal.pone.0184068

32. Okamoto K, Ohishi T, Kuroiwa M, Iemura S ichiro, Natsume T, Seimiya H. MERIT40-dependent recruitment of tankyrase to damaged DNA and its implication for cell sensitivity to DNA-damaging anticancer drugs. Oncotarget. 2018;9(88):35844–35855. doi:10.18632/oncotarget.26312

33. Smith S, Giriat I, Schmitt A, De Lange T. Tankyrase, a Poly(ADP-Ribose) Polymerase at Human Telomeres. Science. 1998;282(5393):1484–1487. doi:10.1126/science.282.5393.1484

34. Smith S, De Lange T. Tankyrase promotes telomere elongation in human cells. Current Biology. 2000;10(20):1299–1302. doi:10.1016/S0960-9822(00)00752-1

35. Cook BD, Dynek JN, Chang W, Shostak G, Smith S. Role for the Related Poly(ADP-Ribose) Polymerases Tankyrase 1 and 2 at Human Telomeres. Molecular and Cellular Biology. 2002;22(1):332–342. doi:10.1128/MCB.22.1.332-342.2002

36. Bae J, Donigian JR, Hsueh AJW. Tankyrase 1 Interacts with Mcl-1 Proteins and Inhibits Their Regulation of Apoptosis. Journal of Biological Chemistry. 2003;278(7):5195–5204. doi:10.1074/jbc.M201988200

37. Chang W, Dynek JN, Smith S. NuMA is a major acceptor of poly(ADP-ribosyl)ation by tankyrase 1 in mitosis. Biochemical Journal. 2005;391(2):177–184. doi:10.1042/BJ20050885

38. Sbodio JI, Chi NW. Identification of a Tankyrase-binding Motif Shared by IRAP, TAB182, and Human TRF1 but Not Mouse TRF1. Journal of Biological Chemistry. 2002;277(35):31887–31892. doi:10.1074/jbc.M203916200

39. Chang P, Coughlin M, Mitchison TJ. Tankyrase-1 polymerization of poly(ADP-ribose) is required for spindle structure and function. Nat Cell Biol. 2005;7(11):1133–1139. doi:10.1038/ncb1322

40. Chi NW, Lodish HF. Tankyrase Is a Golgi-associated Mitogen-activated Protein Kinase Substrate That Interacts with IRAP in GLUT4 Vesicles. Journal of Biological Chemistry. 2000;275(49):38437–38444. doi:10.1074/jbc.M007635200

41. Zhong L, Ding Y, Bandyopadhyay G, et al. The PARsylation activity of tankyrase in adipose tissue modulates systemic glucose metabolism in mice. Diabetologia. 2016;59(3):582–591. doi:10.1007/s00125-015-3815-1

42. Su Z, Deshpande V, James DE, Stöckli J. Tankyrase modulates insulin sensitivity in skeletal muscle cells by regulating the stability of GLUT4 vesicle proteins. Journal of Biological Chemistry. 2018;293(22):8578–8587. doi:10.1074/jbc.RA117.001058

43. Sadler JBA, Lamb CA, Welburn CR, et al. The deubiquitinating enzyme USP25 binds tankyrase and regulates trafficking of the facilitative glucose transporter GLUT4 in adipocytes. Sci Rep. 2019;9(1):4710. doi:10.1038/s41598-019-40596-5

44. Lau T, Chan E, Callow M, et al. A Novel Tankyrase Small-Molecule Inhibitor Suppresses *APC* Mutation–Driven Colorectal Tumor Growth. Cancer Res. 2013;73(10):3132–3144. doi:10.1158/0008-5472.CAN-12-4562

45. Zhong Y, Katavolos P, Nguyen T, et al. Tankyrase Inhibition Causes Reversible Intestinal Toxicity in Mice with a Therapeutic Index < 1. Toxicol Pathol. 2016;44(2):267–278. doi:10.1177/0192623315621192

46. Shirai F, Mizutani A, Yashiroda Y, et al. Design and Discovery of an Orally Efficacious Spiroindolinone-Based Tankyrase Inhibitor for the Treatment of Colon Cancer. J Med Chem. 2020;63(8):4183–4204. doi:10.1021/acs.jmedchem.0c00045

47. Brinch SA, Amundsen-Isaksen E, Espada S, et al. The Tankyrase Inhibitor OM-153 Demonstrates Antitumor Efficacy and a Therapeutic Window in Mouse Models. Cancer Research Communications. 2022;2(4):233–245. doi:10.1158/2767-9764.CRC-22-0027

48. Xu W, Lau YH, Fischer G, et al. Macrocyclized Extended Peptides: Inhibiting the Substrate-Recognition Domain of Tankyrase. J Am Chem Soc. 2017;139(6):2245–2256. doi:10.1021/jacs.6b10234

49. Cheng H, Li X, Wang C, et al. Inhibition of tankyrase by a novel small molecule significantly attenuates prostate cancer cell proliferation. Cancer Letters. 2019;443:80–90. doi:10.1016/j.canlet.2018.11.013

50. Zhu H, Gao Y, Liu L, et al. A novel TNKS/USP25 inhibitor blocks the Wnt pathway to overcome multi-drug resistance in TNKS-overexpressing colorectal cancer. Acta Pharmaceutica Sinica B. 2024;14(1):207–222. doi:10.1016/j.apsb.2023.10.013

51. Pollock K, Liu M, Zaleska M, et al. Fragment-based screening identifies molecules targeting the substrate-binding ankyrin repeat domains of tankyrase. Sci Rep. 2019;9(1):19130. doi:10.1038/s41598-019-55240-5

52. Sowa ST, Vela-Rodríguez C, Galera-Prat A, et al. A FRET-based high-throughput screening platform for the discovery of chemical probes targeting the scaffolding functions of human tankyrases. Sci Rep. 2020;10(1):12357. doi:10.1038/s41598-020-69229-y

53. European chemical biology database. https://ecbd.eu/

54. Škuta C, Müller T, Voršilák M, et al. ECBD: European chemical biology database. Nucleic Acids Research. 2025;53(D1):D1383–D1392. doi:10.1093/nar/gkae904

55. Asif M, Alghamdi S. An Overview on Biological Importance of Pyrrolone and Pyrrolidinone Derivatives as Promising Scaffolds. Russ J Org Chem. 2021;57(10):1700–1718. doi:10.1134/S1070428021100201

56. Starosyla SA, Volynets GP, Lukashov SS, et al. Identification of apoptosis signal-regulating kinase 1 (ASK1) inhibitors among the derivatives of benzothiazol-2-yl-3-hydroxy-5-phenyl-1, 5-dihydro-pyrrol-2-one. Bioorganic & Medicinal Chemistry. 2015;23(10):2489–2497. doi:10.1016/j.bmc.2015.03.056

57. Ma K, Wang P, Fu W, et al. Rational design of 2-pyrrolinones as inhibitors of HIV-1 integrase. Bioorganic & medicinal chemistry letters. 2011;21(22):6724–6727. doi:10.1016/j.bmcl.2011.09.054

58. Gein V, Buldakova E, Korol A, Veikhman G, Dmitriev M. Synthesis of 5-Aryl-4-aroyl-3-hydroxy-1-cyanomethyl-3-pyrrolin-2-ones. Russian Journal of General Chemistry. 2018;88:908–911. doi:10.1134/S1070363218050110

59. Voronkov A, Holsworth DD, Waaler J, et al. Structural Basis and SAR for G007-LK, a Lead Stage 1,2,4-Triazole Based Specific Tankyrase 1/2 Inhibitor. J Med Chem. 2013;56(7):3012–3023. doi:10.1021/jm4000566

60. Bian J, Dannappel M, Wan C, Firestein R. Transcriptional Regulation of Wnt/β-Catenin Pathway in Colorectal Cancer. Cells. 2020;9(9):2125. doi:10.3390/cells9092125

61. Jeong JY, Yim HS, Ryu JY, et al. One-step sequence-and ligation-independent cloning as a rapid and versatile cloning method for functional genomics studies. Applied and environmental microbiology. 2012;78(15):5440–5443. doi:10.1128/AEM.00844-12

62. New England Biolabs. High Efficiency Transformation Protocol (C2987H/C2987I). https://www.neb.com/en/protocols/0001/01/01/high-efficiency-transformation-protocol-c2987

63. Reyrat JM, Pelicic V, Gicquel B, Rappuoli R. Counterselectable markers: untapped tools for bacterial genetics and pathogenesis. Infection and immunity. 1998;66(9):4011–4017. doi:10.1128/iai.66.9.4011-4017.1998

64. Hynes MF, Quandt J, O’Connell MP, Pühler A. Direct selection for curing and deletion of Rhizobium plasmids using transposons carrying the Bacillus subtilis sacB gene. Gene. 1989;78(1):111–120. doi:10.1016/0378-1119(89)90319-3

65. Fisher CL, Pei GK. Modification of a PCR-based site-directed mutagenesis method. Biotechniques. 1997;23(4):570–574. doi:10.2144/97234bm01

66. New England Biolabs. Transformation Protocol for BL21(DE3) Competent Cells (C2527). https://www.neb.com/en/protocols/0001/01/01/transformation-protocol-for-bl21-de3-competent-cells-c2527

67. Marley J, Lu M, Bracken C. A method for efficient isotopic labeling of recombinant proteins. Journal of biomolecular NMR. 2001;20:71–75. doi:10.1023/a:1011254402785

68. Skinner SP, Fogh RH, Boucher W, Ragan TJ, Mureddu LG, Vuister GW. CcpNmr AnalysisAssign: a flexible platform for integrated NMR analysis. Journal of biomolecular NMR. 2016;66:111–124. doi:10.1007/s10858-016-0060-y

69. Zaleska M, Pollock K, Collins I, Guettler S, Pfuhl M. Solution NMR assignment of the ARC4 domain of human tankyrase 2. Biomol NMR Assign. 2019;13(1):255–260. doi:10.1007/s12104-019-09887-w

70. Williamson MP. Using chemical shift perturbation to characterise ligand binding. Progress in Nuclear Magnetic Resonance Spectroscopy. 2013;73:1–16. 10.1016/j.pnmrs.2013.02.001

71. Kabsch W. XDS. Acta Crystallogr D Biol Crystallogr. 2010;66(2):125–132. doi:10.1107/S0907444909047337

72. McCoy AJ, Grosse-Kunstleve RW, Adams PD, Winn MD, Storoni LC, Read RJ. Phaser crystallographic software. J Appl Crystallogr. 2007;40(4):658-674. doi:10.1107/S0021889807021206

73. Emsley P, Cowtan K. *Coot*: model-building tools for molecular graphics. Acta Crystallogr D Biol Crystallogr. 2004;60(12):2126–2132. doi:10.1107/S0907444904019158

74. Murshudov GN, Skubák P, Lebedev AA, et al. *REFMAC* 5 for the refinement of macromolecular crystal structures. Acta Crystallogr D Biol Crystallogr. 2011;67(4):355–367. doi:10.1107/S0907444911001314

75. Liebschner D, Afonine PV, Baker ML, et al. Macromolecular structure determination using X-rays, neutrons and electrons: recent developments in *Phenix*. Acta Crystallogr D Struct Biol. 2019;75(10):861–877. doi:10.1107/S2059798319011471

76. American Type Culture Collection. Protocol for Wnt-3A Conditioned Medium. https://www.atcc.org/products/crl-2647

77. Masood MM, Pillalamarri VK, Irfan M, et al. Diketo acids and their amino acid/dipeptidic analogues as promising scaffolds for the development of bacterial methionine aminopeptidase inhibitors. RSC Advances. 2015;5(43):34173–34183. doi:10.1039/C5RA03354C

78. Liu Y, Wang Y, Song H, Zhou Z, Tang C. Asymmetric Organocatalytic Cascade Michael/Hemiketalization/Retro-Aldol Reaction of 2-[(E)-2-Nitrovinyl] phenols with 2, 4-Dioxo-4-arylbutanoates: A Convenient Access to Chiral α-Keto Esters. Advanced Synthesis & Catalysis. 2013;355(13):2544–2549. doi:10.1002/adsc.201300552

79. Kamal A, Tamboli JR, Ramaiah MJ, et al. Anthranilamide–Pyrazolo [1, 5-a] pyrimidine Conjugates as p53 Activators in Cervical Cancer Cells. ChemMedChem. 2012;7(8):1453–1464. doi:10.1002/cmdc.201200205

80. Chen L, Zhang Y, Tian L, et al. Noncovalent EGFR T790M/L858R inhibitors based on diphenylpyrimidine scaffold: design, synthesis, and bioactivity evaluation for the treatment of NSCLC. European Journal of Medicinal Chemistry. 2021;223:113626. doi:10.1016/j.ejmech.2021.113626

