## Supplementary information for "Discovery of Tankyrase scaffolding inhibitor specifically targeting the ARC4 peptide binding domain"

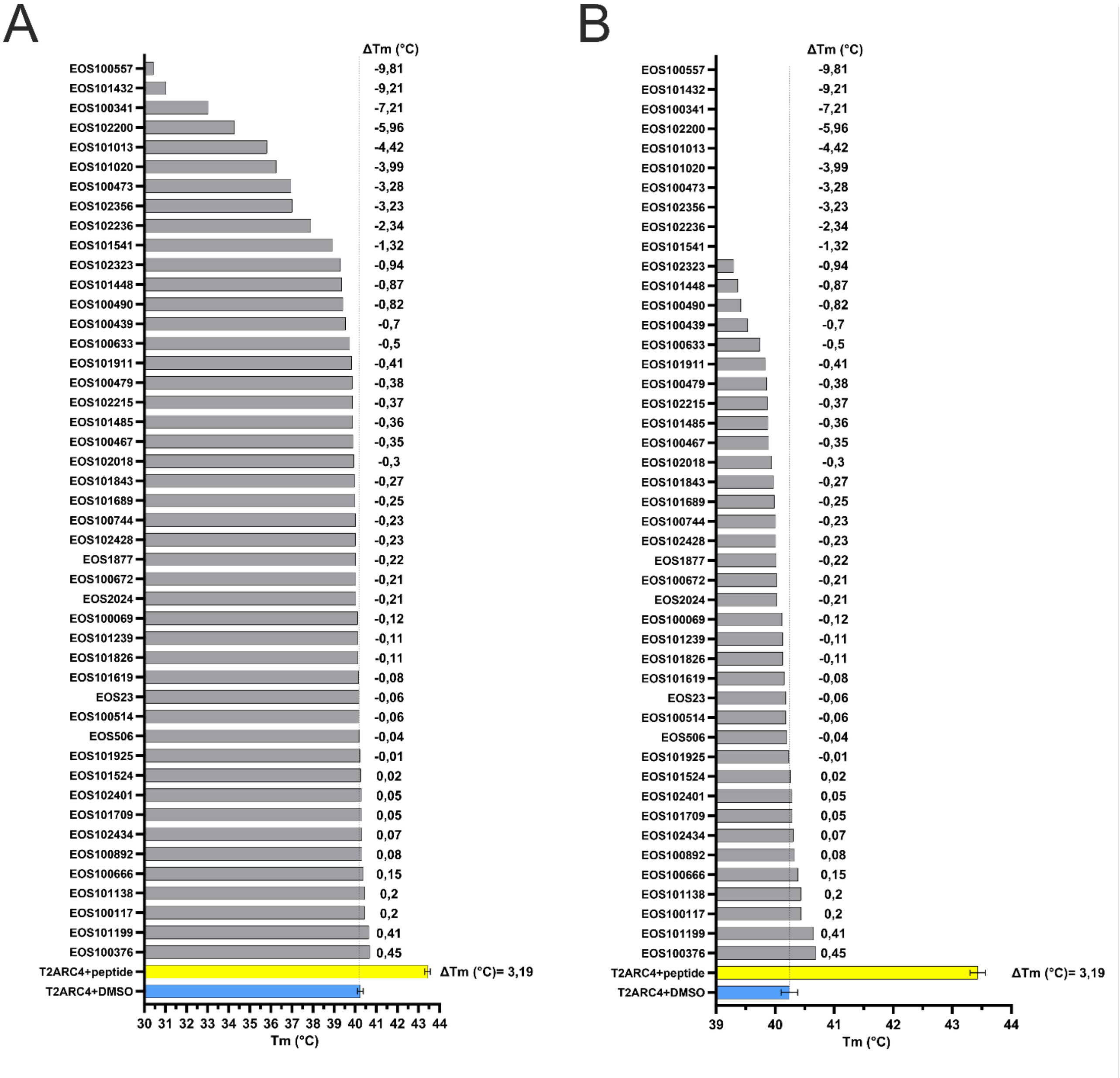

**Figure S1. DSF assay results for validation of primary hits identified in the EU-OPENSCREEN Pilot library.**

(**A**) DSF assay results indicate the level of structural stabilization of the primary hits from the Pilot Library. Compounds (studied at 100 µM) are named after their EU-OPENSCREEN identity code and are available for consultation on ECBD.[1,2] Controls are represented by TNKS2 ARC4 in presence of 100 µM DMSO, and TNKS2 ARC4 mixed with 100 µM peptide designed for optimal binding of the protein (REAGDGEE).[3] Controls were studied in at least four replicates and errors are shown, while compounds were studied with single measurements. The melting temperature (Tm) is indicated for each analyzed compound and controls. In addition, ΔTm, which is the difference between the Tm of the control consisting in TNKS2 ARC4 with DMSO and the Tm measured for the compound, is shown. The ΔTm value for the control encompassing TNKS2 ARC4 and the peptide is also reported to indicate the optimal increase in Tm. None of the primary hits was found to significantly stabilize TNKS2 ARC4. It was not possible to measure the melting curve of TNKS2 ARC4 in presence of EOS101941, EOS2202, EOS101061, EOS101771. (**B**) DSF assay measurements for the Pilot Library compounds on a shorter Tm scale (39-44 °C).

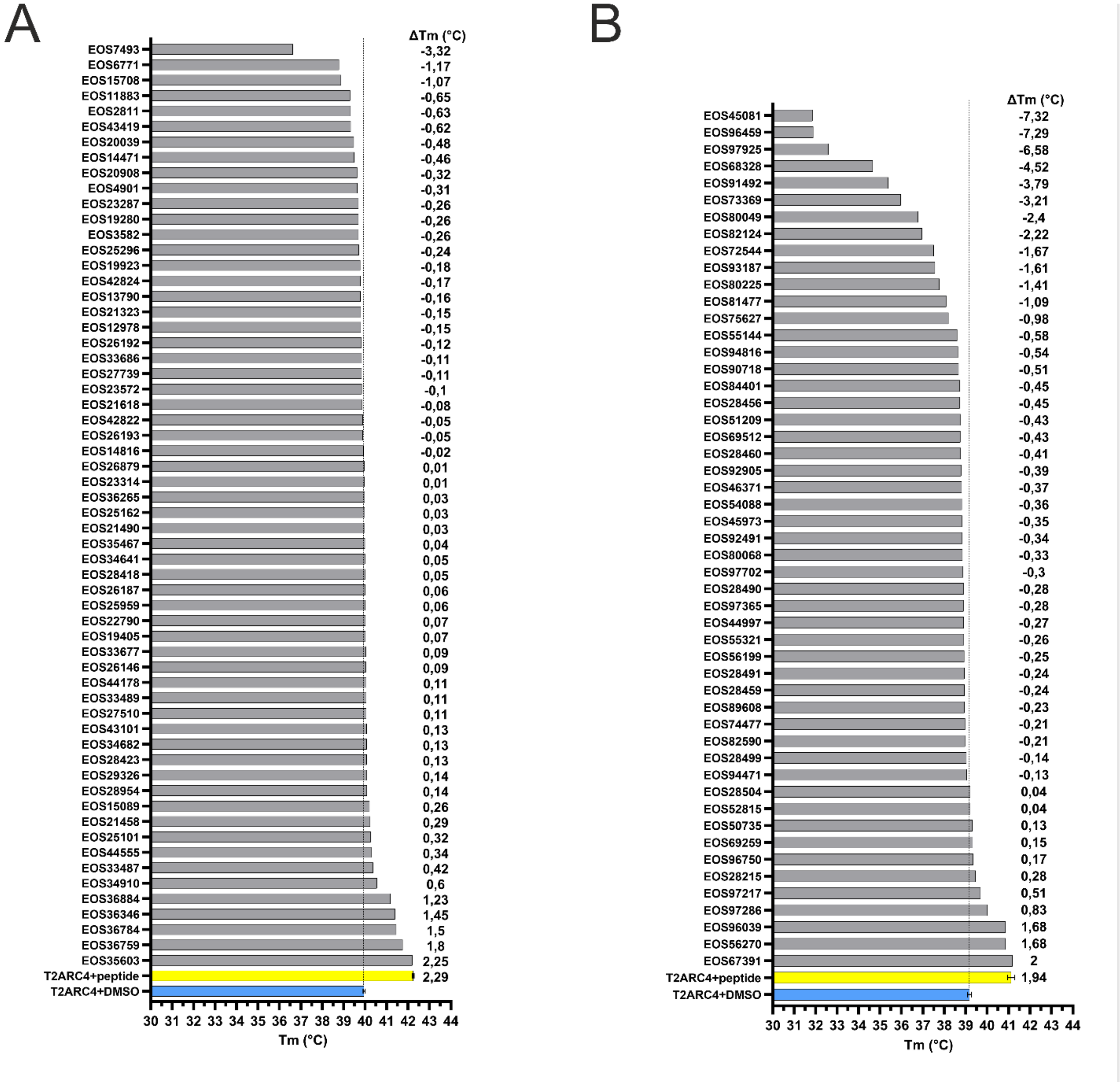

**Figure S2. DSF assay results for validation of primary hits identified in the EU-OPENSCREEN Commercials Diversity Library.**

(**A**) and (**B**) show DSF assay validation measurements of TNKS2 ARC4 with primary hits from the Commercial Diversity Library. Compounds are tested at 100 µM. They are named after their EU-OPENSCREEN identity codes and the identities are available for consultation on ECBD.[1,2] (**A**) and (**B**) show results from two independent experiments. For each experiment, each compound was tested with a single measurement, while controls (TNKS2 ARC4 with 100 µM optimized peptide, TNKS2 ARC4 with 100 µM DMSO) are studied in at least four replicates and errors are shown. Moreover, stabilization over the control (TNKS2 ARC4 with DMSO) is indicated (ΔTm °C). Several hits sharing a common pyrrolone scaffold were found to significantly stabilize TNKS2 ARC4 (EOS35603, EOS36759, EOS36784, EOS36346, EOS36884, EOS67391, EOS56270, EOS96039, EOS97286). It was not possible to measure the melting curve of TNKS2 ARC4 in presence of EOS21552, EOS54043, EOS54043.

**Table S1**. Synthesized compounds (**S1**-**S12**) are analogues of **1** and are characterized by different $R^1$, $R^2$ and $R^3$ substituents.

| Code | Structure | | |
|---|---|---|---|
| | R1 | R2 | R3 |
| 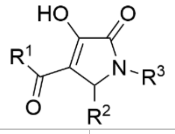 | | | |
| **S1** | 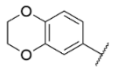 | 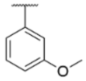 | 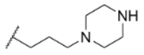 |
| **S2** | 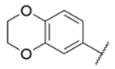 | 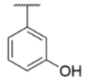 | 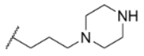 |
| **S3** | 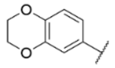 | 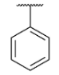 | 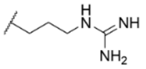 |
| **S4** | 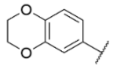 | 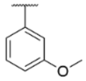 | 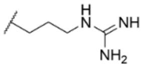 |
| **S5** | 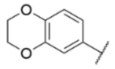 | 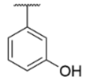 | 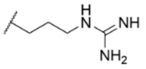 |
| **S6** | 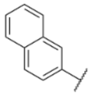 | 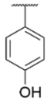 | 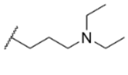 |
| **S7** | 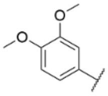 | 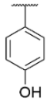 | 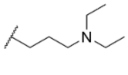 |
| **S8 (ARCher-142)** | 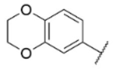 | 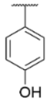 | 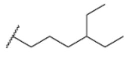 |
| **S9** | 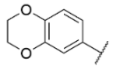 | 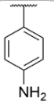 | 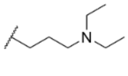 |
| **S10** | 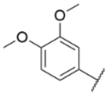 | 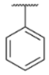 | 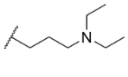 |

| | | | |
|---|---|---|---|
| **S11** | 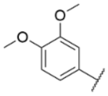 | 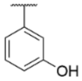 | 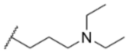 |
| **S12** | 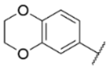 | 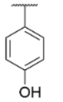 | 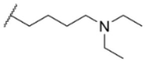 |

**Table S2**. Structures, FRET-based $IC_{50}$ and DSF measurements of **1** analogues for investigation of SARs (including compounds from **Table 1** and synthesized compounds of **Table S1**). FRET-based potency measurements are indicated as $IC_{50}$ (μM) and $pIC_{50}$ ± SEM, with number of repetitions n=3 (n=1 for compounds with $IC_{50}$ >1000 μM). Thermal stabilization (ΔTm ± SD) is indicated (n=4).

| Code | Structure | | | $IC_{50}$ (μM)<br>$pIC_{50}$ ± SEM | ΔTm ± SD (°C) |
|---|---|---|---|---|---|
| | R1 | R2 | R3 | | |
| 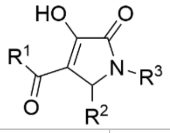 | | | | | |
| **10** | 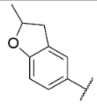 | 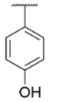 | 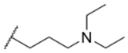 | 31<br>(4.51 ± 0.01) | 2.67 ± 0.46 |
| **27** | 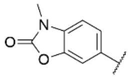 | 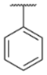 | 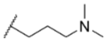 | >1000 | -0.03 ± 0.11 |
| **11** | 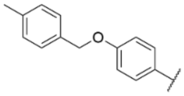 | 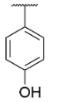 | 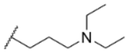 | 552<br>(3.26 ± 0.02) | 0.07 ± 0.07 |
| **28** | 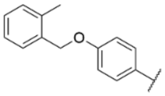 | 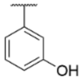 | 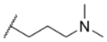 | >1000 | -0.20 ± 0.22 |
| **S7** | 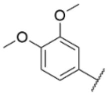 | 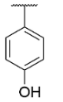 | 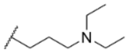 | >1000 | 0.04 ± 0.03 |
| **S11** | 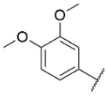 | 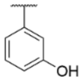 | 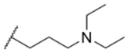 | >1000 | 0.08 ± 0.40 |
| **S10** | 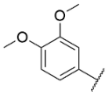 | 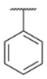 | 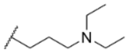 | >1000 | -0.28 ± 0.35 |
| **29** | 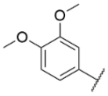 | 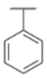 | 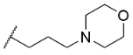 | >1000 | 0.14 ± 0.25 |
| **30** | 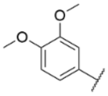 | 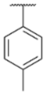 | 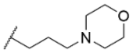 | >1000 | 0.12 ± 0.25 |

| | | | | | |
|---|---|---|---|---|---|
| **31** | 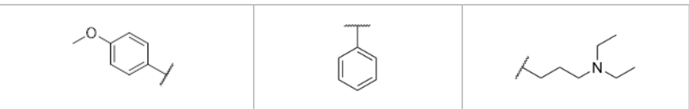 | | | 136 (3.86 ± 0.03) | 1.46 ± 0.52 |
| **32** | 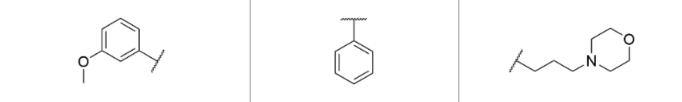 | | | >1000 | 0.23 ± 0.25 |
| **33** | 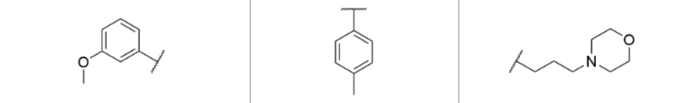 | | | >1000 | 0.36 ± 0.33 |
| **34** | 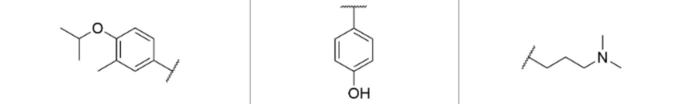 | | | >1000 | -0.08 ± 0.41 |
| **S6** | 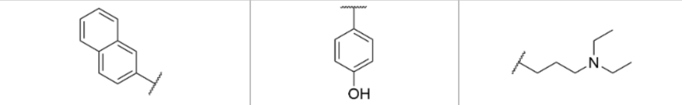 | | | >1000 | -0.06 ± 0.04 |
| **12** | 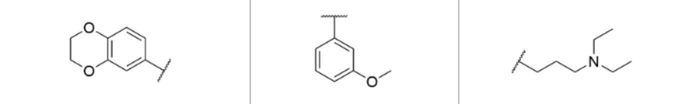 | | | 11 (4.95 ± 0.03) | 4.03 ± 0.39 |
| **S1** | 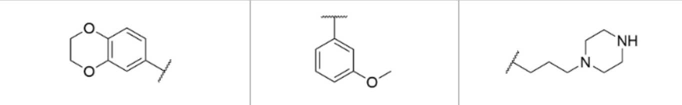 | | | 250 (3.60 ± 0.03) | 0.69 ± 0.30 |
| **S4** | 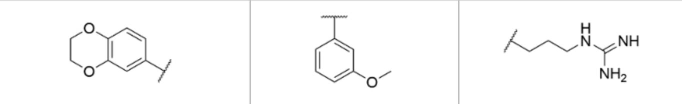 | | | 142 (3.85 ± 0.02) | 1.10 ± 0.12 |
| **13** | 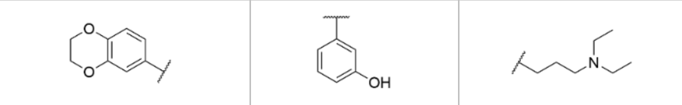 | | | 24 (4.62 ± 0.01) | 4.82 ± 0.49 |
| **14** | 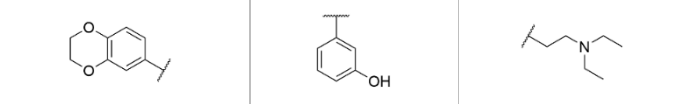 | | | 139 (3.86 ± 0.04) | 1.40 ± 0.36 |
| **S2** | 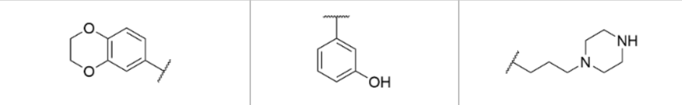 | | | 302 (3.52 ± 0.01) | 0.80 ± 0.16 |
| **15** | 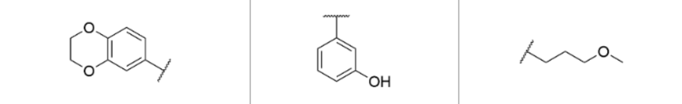 | | | 368 (3.43 ± 0.08) | 0.78 ± 0.43 |
| **35** | 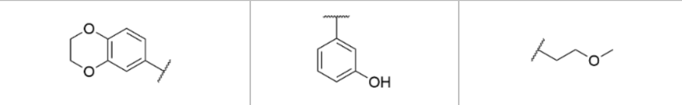 | | | 305 (3.52 ± 0.02) | 0.82 ± 0.38 |
| **36** | 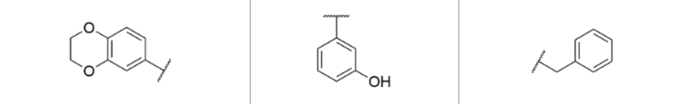 | | | 23 (4.63 ± 0.03) | 0.45 ± 0.25 |
| **16** | 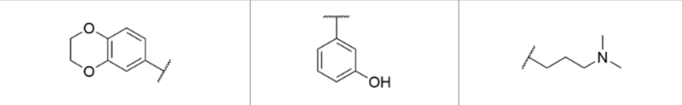 | | | 31 (4.51 ± 0.02) | 2.96 ± 0.36 |
| **17** | 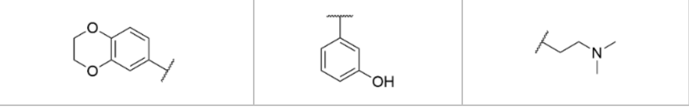 | | | 274 (3.56 ± 0.02) | 0.98 ± 0.32 |

| | | | | | |
|---|---|---|---|---|---|
| **18** | 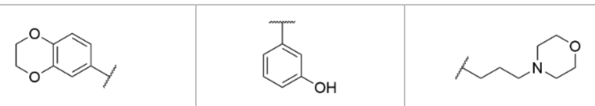 | | | 85<br>(4.07 ± 0.02) | 2.25 ± 0.48 |
| **S5** | 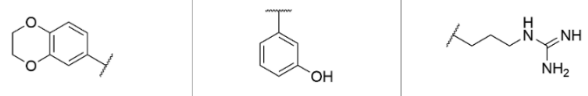 | | | 121<br>(3.92 ± 0.06) | 1.31 ± 0.17 |
| **19** | 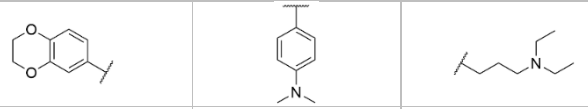 | | | 10<br>(5.00 ± 0.02) | 4.14 ± 0.12 |
| **S9** | 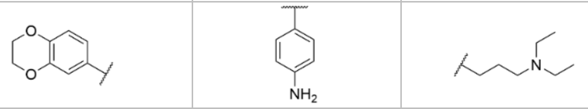 | | | 17<br>(4.76 ± 0.03) | 2.78 ± 0.18 |
| **20** | 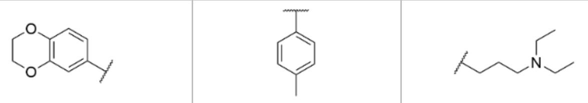 | | | 78<br>(4.11 ± 0.07) | 3.43 ± 0.66 |
| **37** | 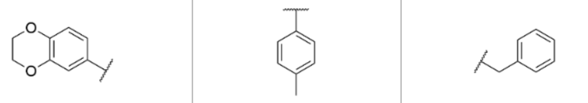 | | | 71<br>(4.15 ± 0.05) | 0.51 ± 0.31 |
| **3** | 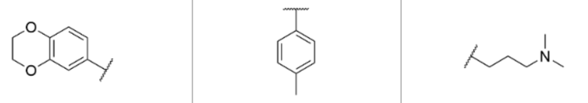 | | | 51<br>(4.29 ± 0.01) | 2.34 ± 0.30 |
| **38** | 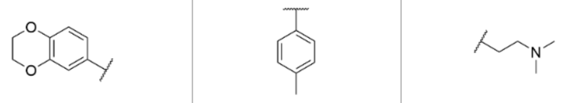 | | | 493<br>(3.31 ± 0.05) | 0.67 ± 0.28 |
| **4** | 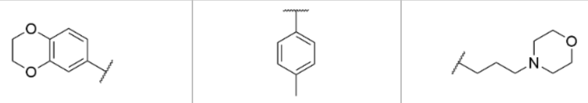 | | | 77<br>(4.11 ± 0.02) | 2.19 ± 0.31 |
| **21** | 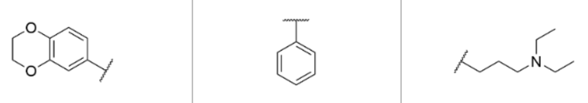 | | | 36<br>(4.44 ± 0.05) | 4.73 ± 0.29 |
| **39** | 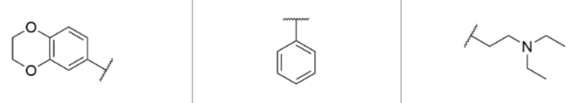 | | | 255<br>(3.59 ± 0.04) | 1.05 ± 0.36 |
| **40** | 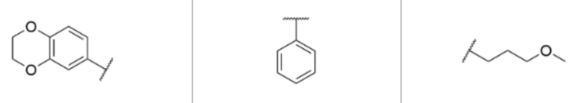 | | | 360<br>(3.44 ± 0.02) | 0.47 ± 0.31 |
| **41** | 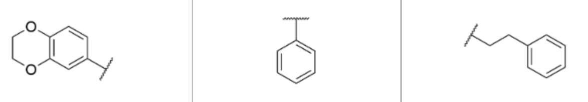 | | | 65<br>(4.18 ± 0.01) | 0.91 ± 0.31 |
| **22** | 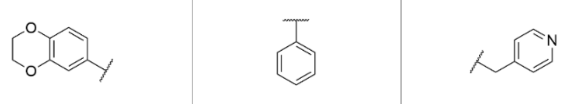 | | | 325<br>(3.49 ± 0.07) | 0.48 ± 0.27 |
| **42** | 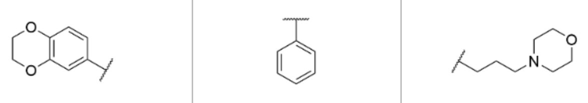 | | | 101<br>(4.00 ± 0.01) | 2.26 ± 0.34 |
| **23** | 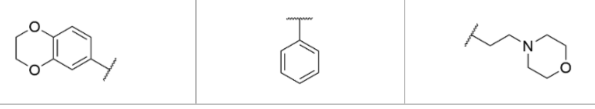 | | | 411<br>(3.39 ± 0.05) | 0.89 ± 0.33 |

| | | | | | |
|---|---|---|---|---|---|
| **S3** | 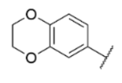 | 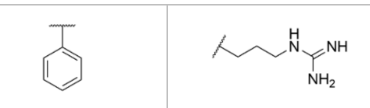 | | 133 (3.86 ± 0.01) | 1.20 ± 0.19 |
| **5** | 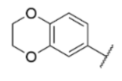 | 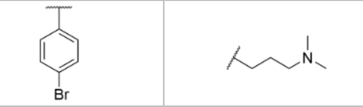 | | 70 (4.15 ± 0.04) | 1.49 ± 0.16 |
| **2** | 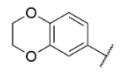 | 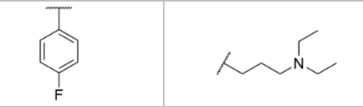 | | 32 (4.49 ± 0.04) | 2.38 ± 0.52 |
| **43** | 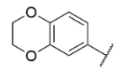 | 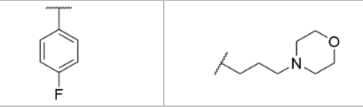 | | 114 (3.94 ± 0.02) | 1.61 ± 0.37 |
| **24** | 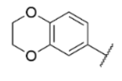 | 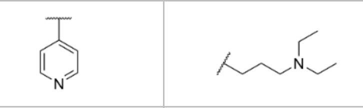 | | 56 (4.25 ± 0.002) | 2.75 ± 0.44 |
| **44** | 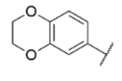 | 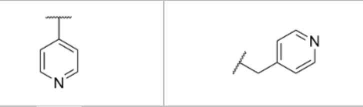 | | > 1000 | 0.21 ± 0.26 |
| **6** | 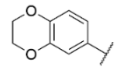 | 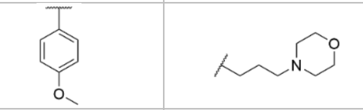 | | 82 (4.09 ± 0.01) | 1.81 ± 0.31 |
| **7** | 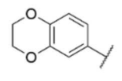 | 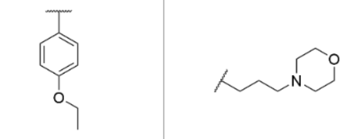 | | 117 (3.93 ± 0.09) | 1.42 ± 0.16 |
| **9** | 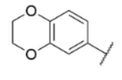 | 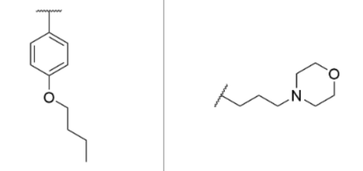 | | 81 (4.09 ± 0.05) | 1.40 ± 0.18 |
| **8** | 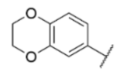 | 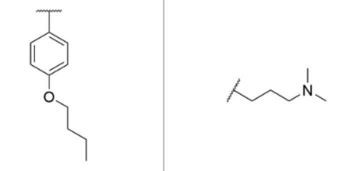 | | 69 (4.16 ± 0.08) | 1.65 ± 0.24 |
| **45** | 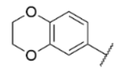 | 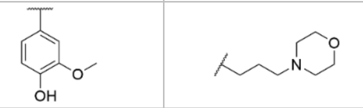 | | 78 (4.11 ± 0.07) | 2.32 ± 0.33 |
| **S8** | 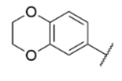 | 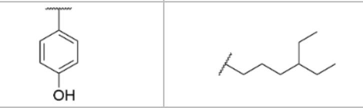 | | 8 (5.11 ± 0.04) | 2.44 ± 0.12 |
| **S12** | 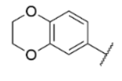 | 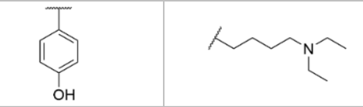 | | 74 (4.13 ± 0.03) | 1.97 ± 0.35 |
| **25** | 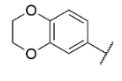 | 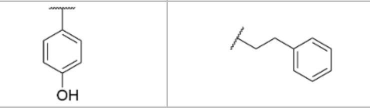 | | 56 (4.25 ± 0.01) | 1.80 ± 0.26 |

| | | | | | |
|---|---|---|---|---|---|
| **26** | 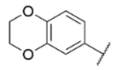 | 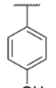 | 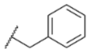 | 30<br>(4.52 ± 0.03) | 0.76 ± 0.33 |

**Table S3**. Data collection and refinement statistics of TNKS2 ARC4 in complex with compound **S8** (**ARCher-142**).

| **Protein, Inhibitor (PDB ID)** | **TNKS2 ARC4 (PDB 9QFC)** |
|---|---|
| **Data collection** | |
| Beamline | I24 (Diamond) |
| Wavelength (Å) | 0.97950 |
| Space group | P 43 21 2 |
| Unit cell dimensions<br>*a, b, c* (Å)<br>α, β, γ (°) | 95.33 95.33 85.94<br>90 90 90 |
| Resolution range(Å) (Outer cell) | 42.63 - 2.55 (2.641 - 2.55) |
| Total n. of reflections | 77082 (7159) |
| N. of unique reflections | 13414 (1304) |
| Completeness (%) | 99.76 (99.85) |
| ⟨*I*/σ(*I*)⟩ | 7.20 (1.63) |
| CC1/2 | 0.985 (0.61) |
| $R_{meas}$ | 0.2441 (1.188) |
| $R_{merge}$ | 22.2 |
| **Refinement** | |
| R-work/R-free | 0.2157/0.2602 |
| **N. of non-hydrogen atoms** | 2640 |
| Protein | 2448 |
| Ligands | 277 |
| Solvent | 49 |
| **RMSD** | |
| Bonds (Å) | 0.002 |
| Angles (°) | 0.38 |
| **Average *B* factors ($Å^2$)** | 37.78 |
| Protein | 37.56 |
| Ligands | 42.70 |
| Solvent | 34.72 |
| **Ramachandran plot** | |
| Favoured (%) | 95.89 |
| Allowed (%) | 4.11 |

*Values within parentheses refers to the highest resolution shell.

**Figure S3**. $^1$H NMR ($CDCl_3$, 400MHz) of ethyl 2,3-dihydro-α,γ-dioxo-1,4-benzodioxin-6-butanoate (**2a**).

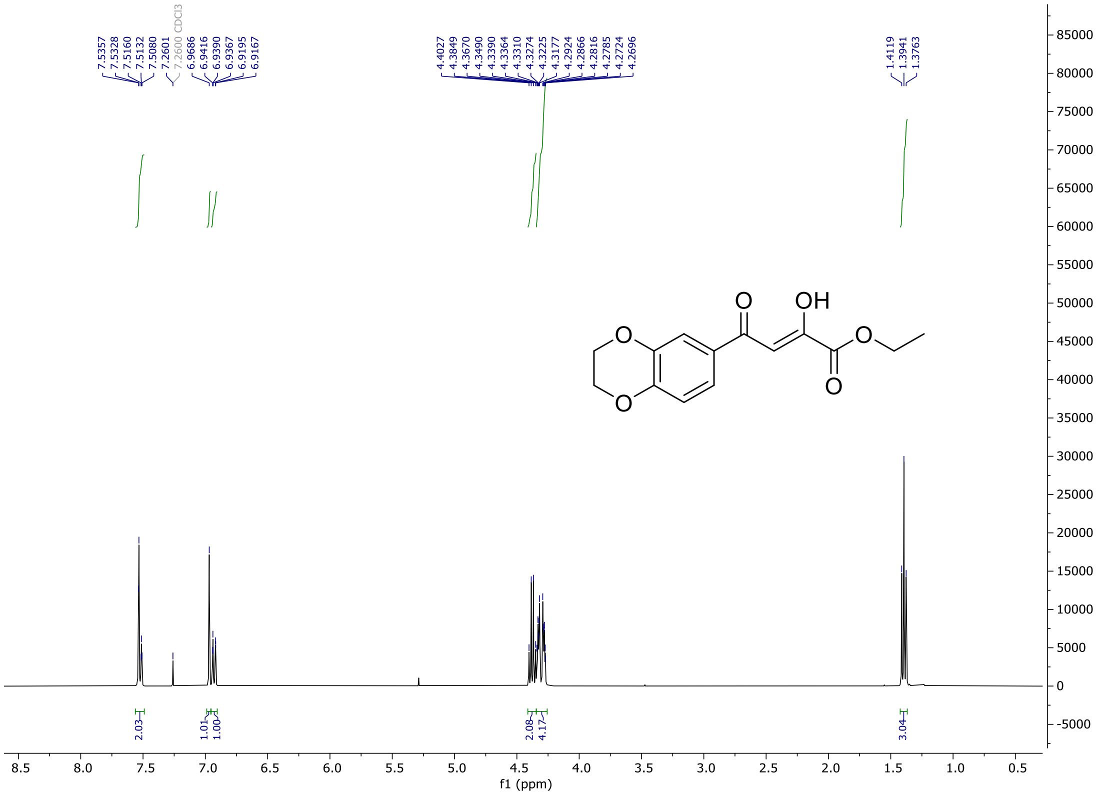

**Figure S4**. $^{13}C$ NMR ($CDCl_3$, 100MHz) of ethyl 2,3-dihydro-α,γ-dioxo-1,4-benzodioxin-6-butanoate (**2a**).

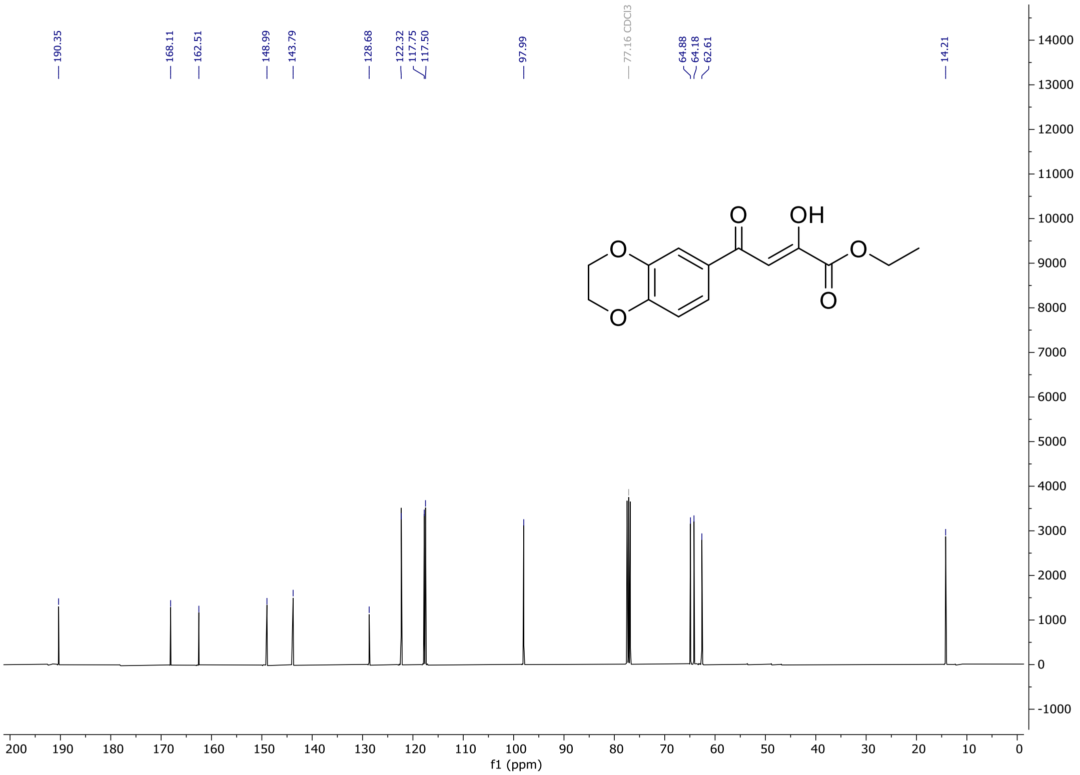

**Figure S5**. $^{1}$H NMR ($CDCl_3$, 400MHz) of methyl 4-(naphthalen-2-yl)-2,4-dioxobutanoate (**2b**).

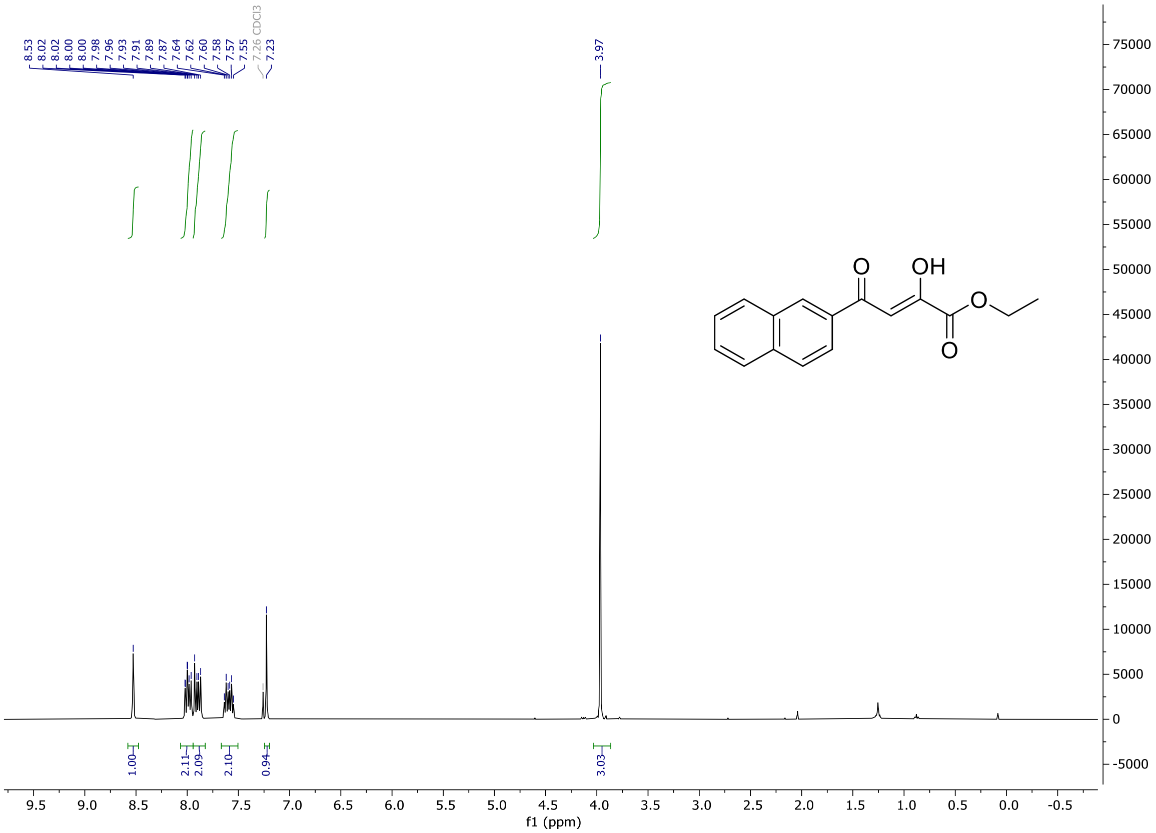

**Figure S6**. $^{13}C$ NMR ($CDCl_3$, 100MHz) of methyl 4-(naphthalen-2-yl)-2,4-dioxobutanoate (**2b**).

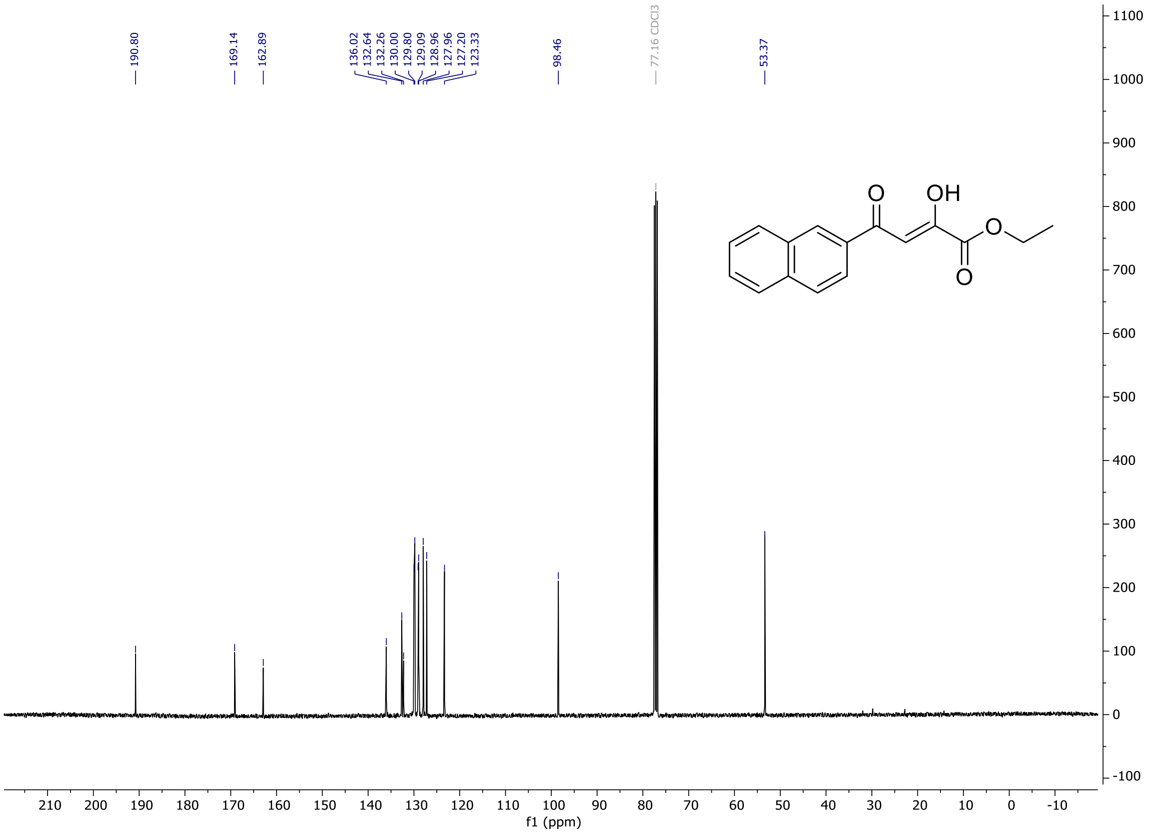

**Figure S7**. $^{1}H$ NMR ($CDCl_3$, 400MHz) of methyl 4-(3,4-dimethoxyphenyl)-2,4-dioxobutanoate (**2c**).

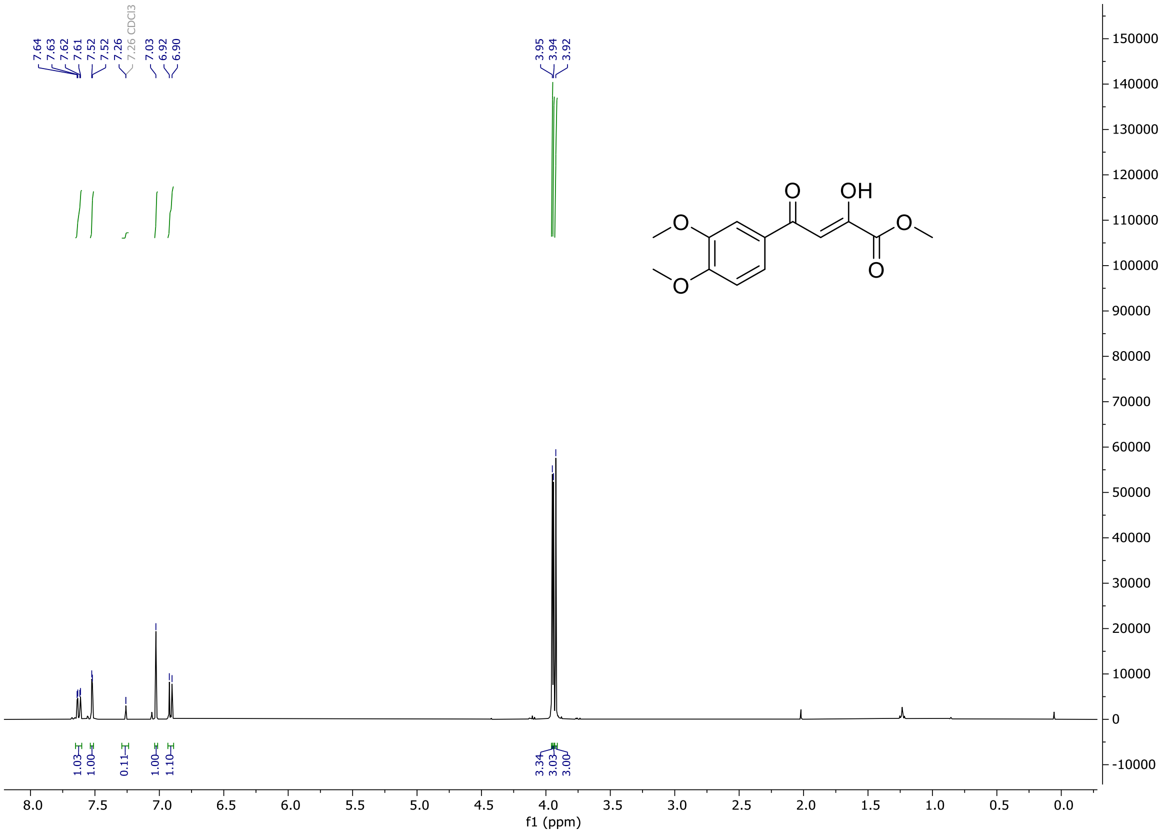

**Figure S8**. $^{13}C$ NMR ($CDCl_3$, 100MHz) of methyl 4-(3,4-dimethoxyphenyl)-2,4-dioxobutanoate (**2c**).

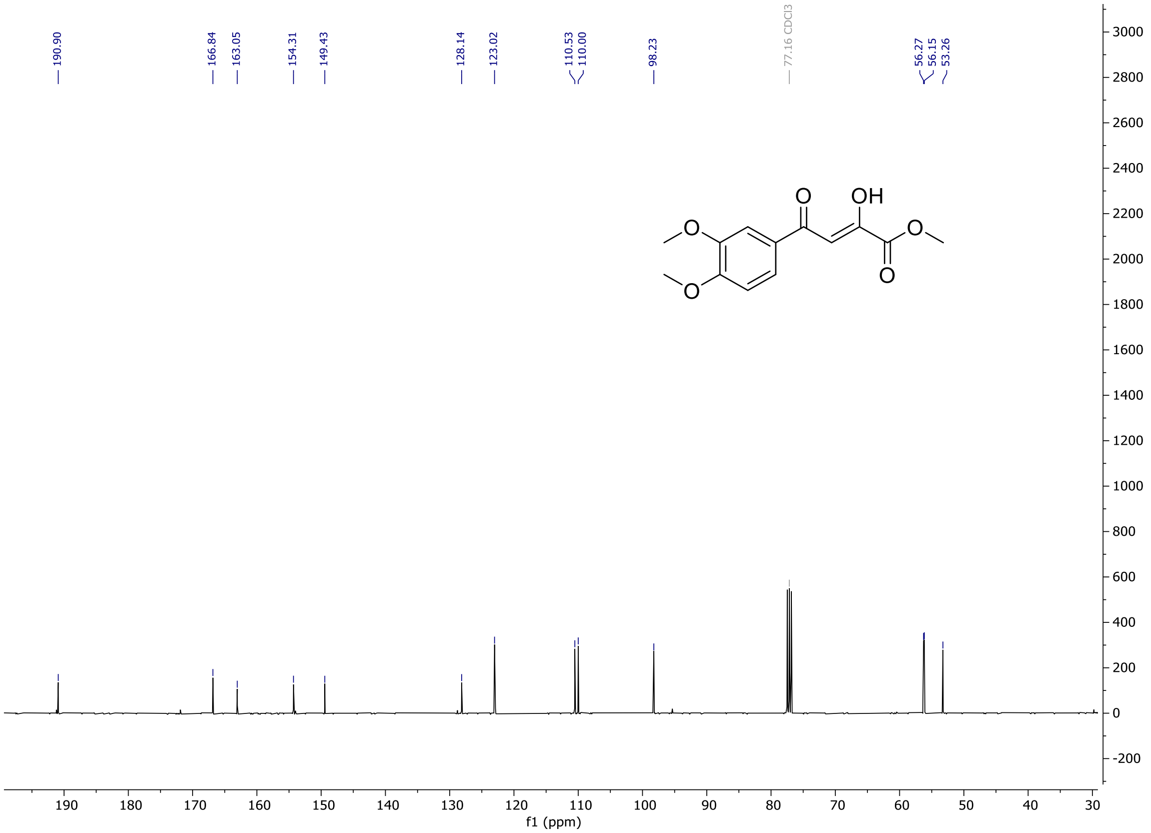

**Figure S9**. $^{1}H$ NMR ($(CD_3)_2SO$, 800 MHz) of **S1**.

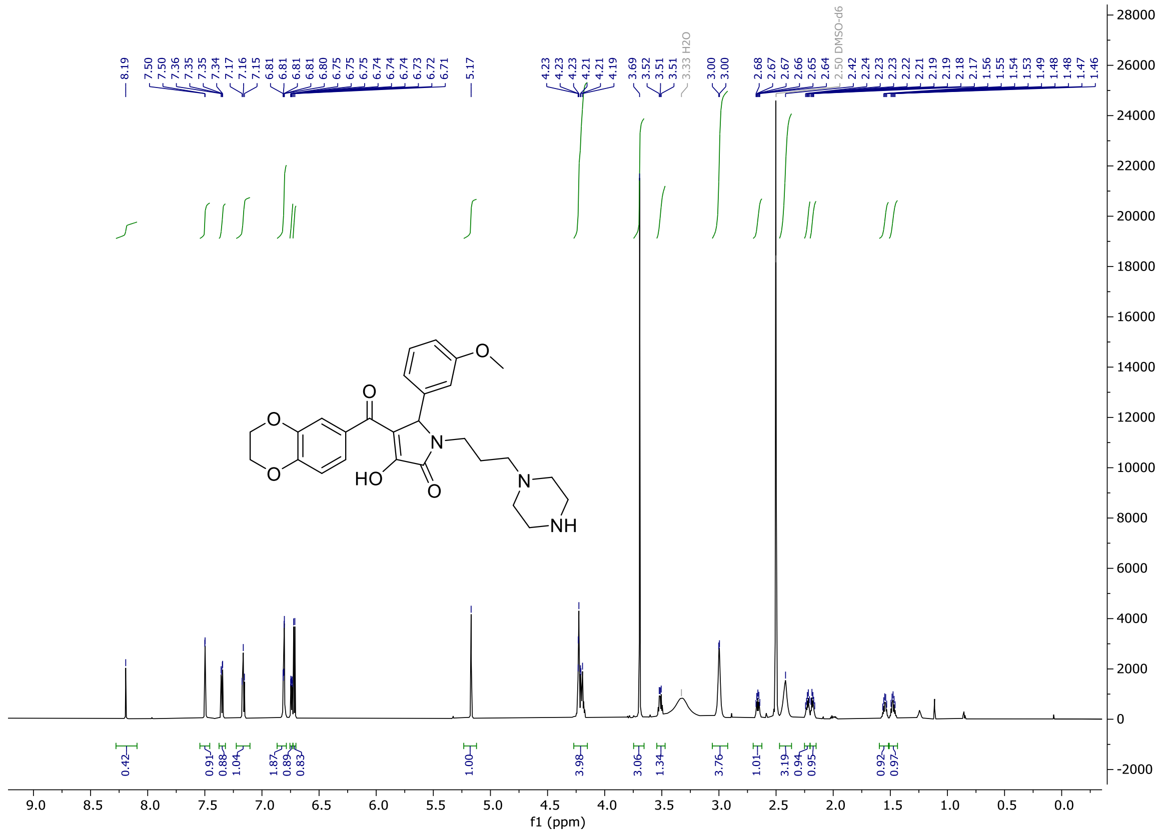

**Figure S10**. $^{13}C$ NMR ($(CD_3)_2SO$, 200 MHz) of **S1**.

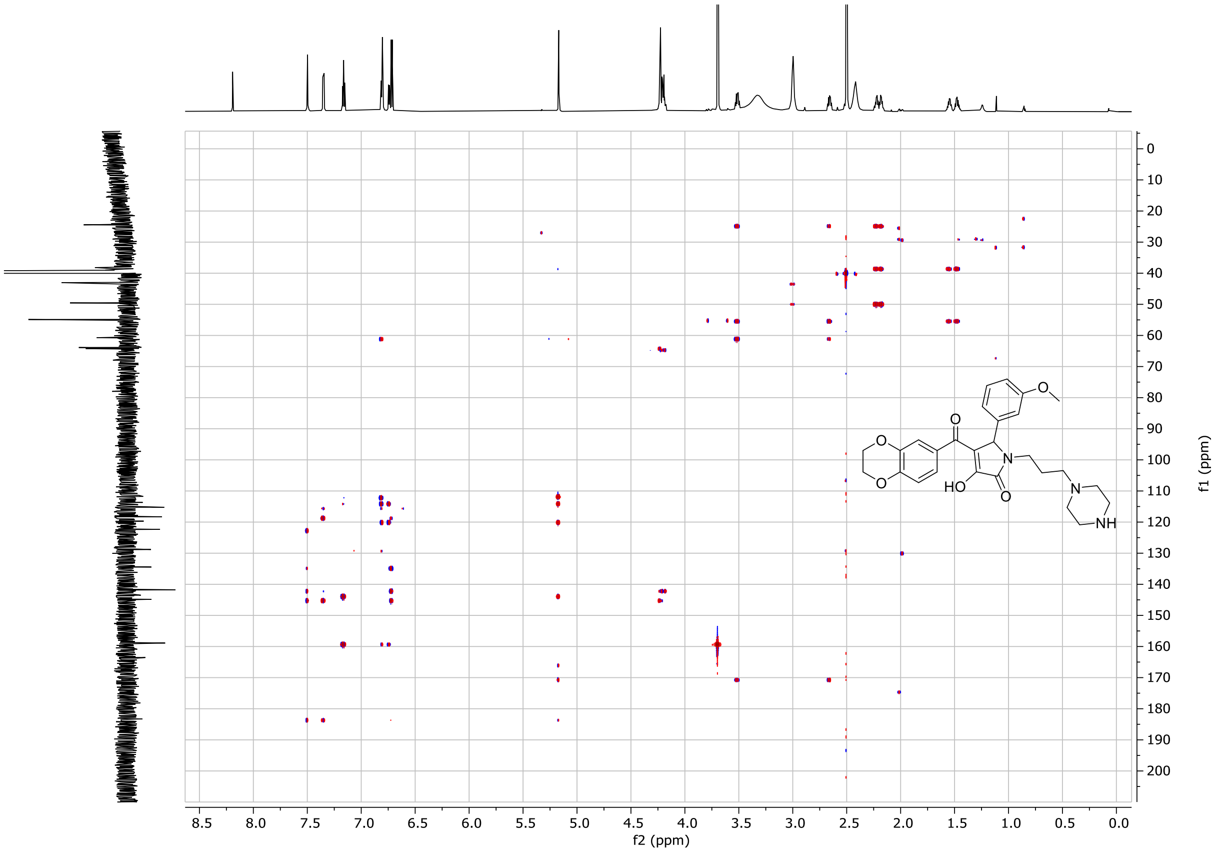

**Figure S11**. $^{1}H$ NMR ($(CD_3)_2SO$, 600MHz) of **S2**.

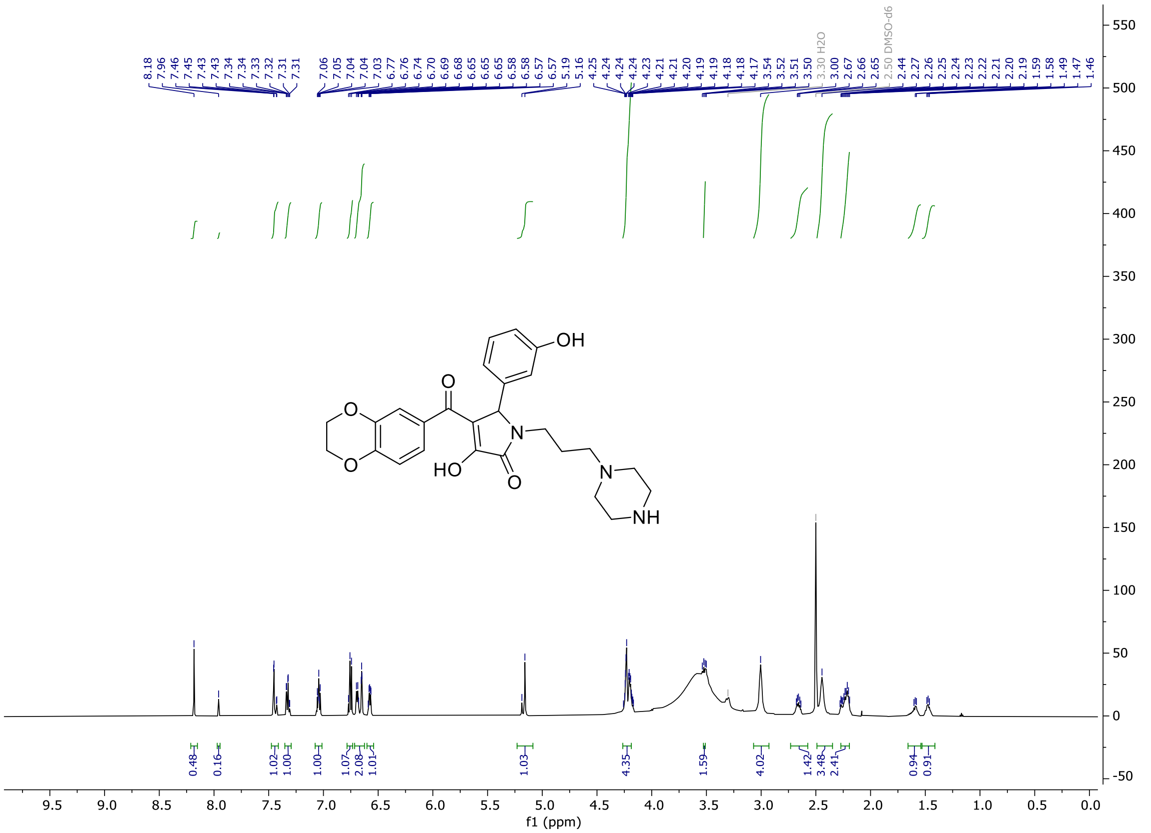

**Figure S12**. $^{13}C$ NMR ($(CD_3)_2SO$, 150MHz) of **S2**.

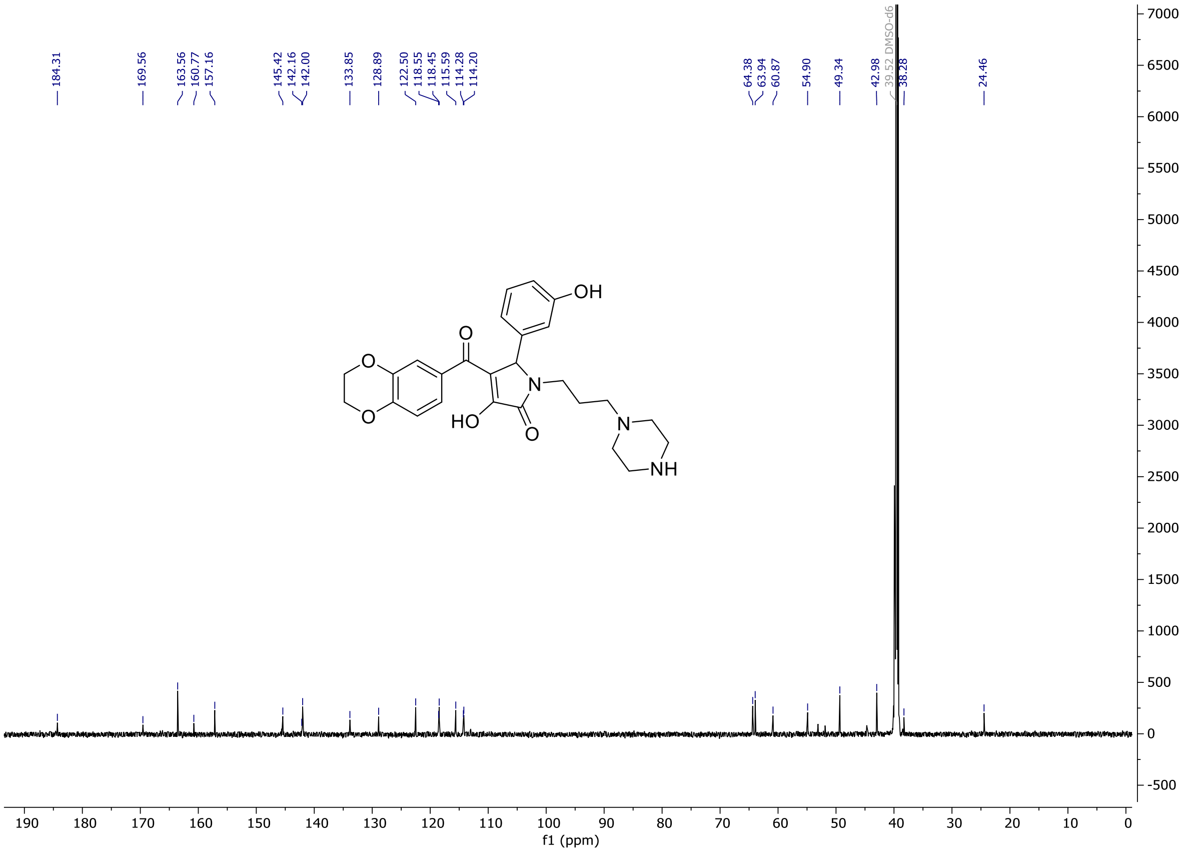

**Figure S13**. $^1$H NMR ($(CD_3)_2SO$, 800 MHz) of **S3**.

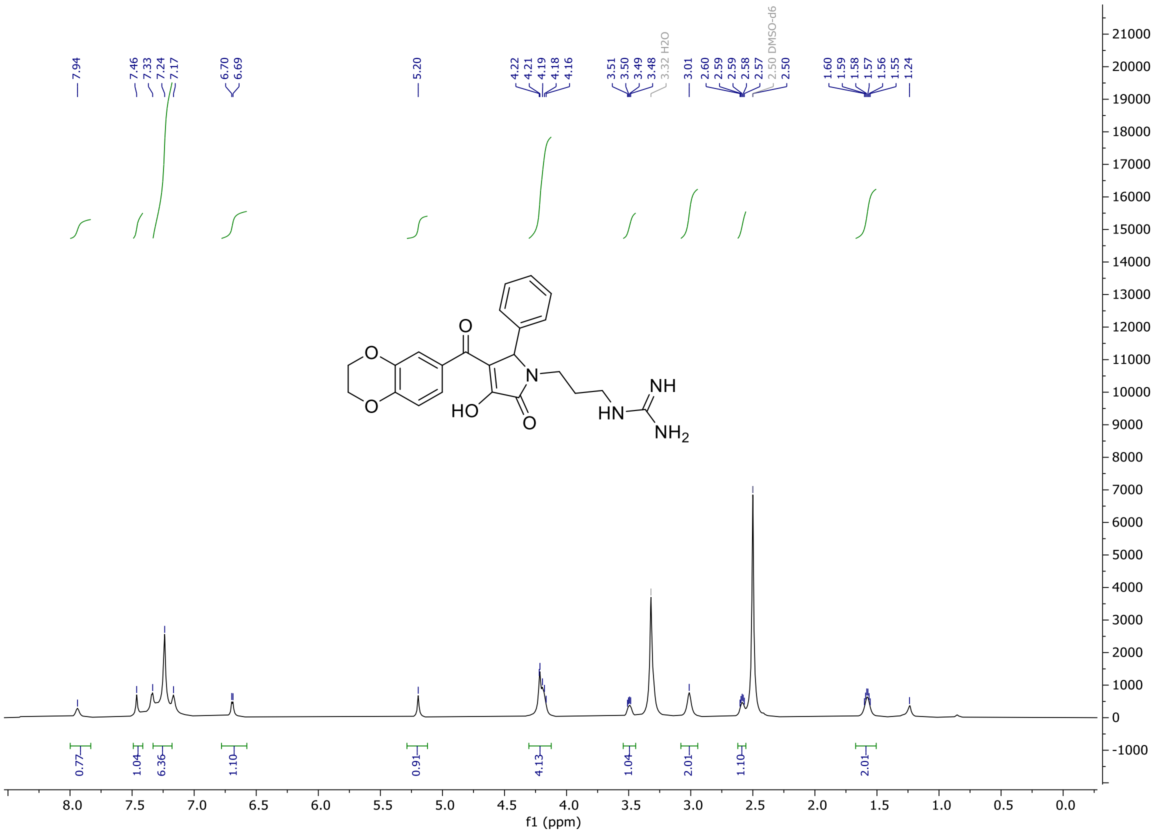

**Figure S14**. $^{13}C$ NMR ($(CD_3)_2SO$, 200 MHz) of **S3**.

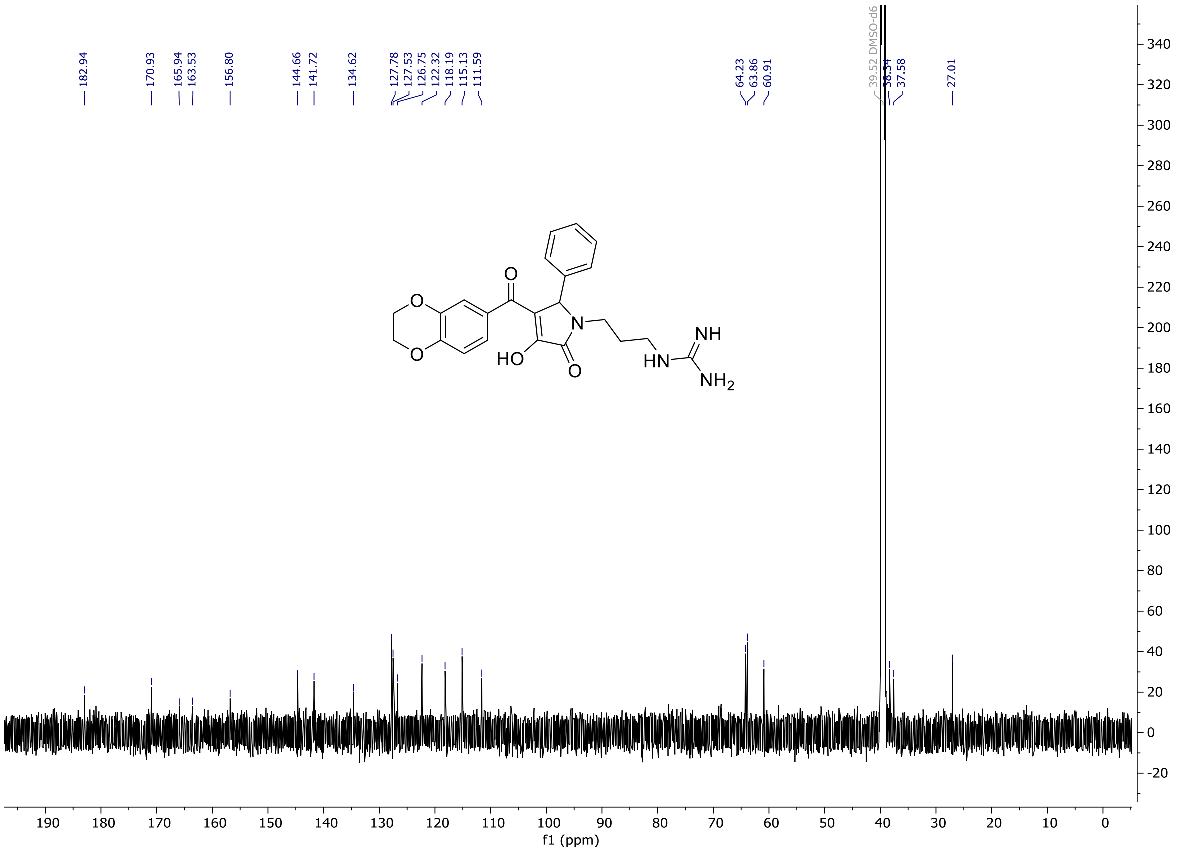

**Figure S15**. $^{1}H$ NMR ($CD_3OD$, 400 MHz) of **S4**.

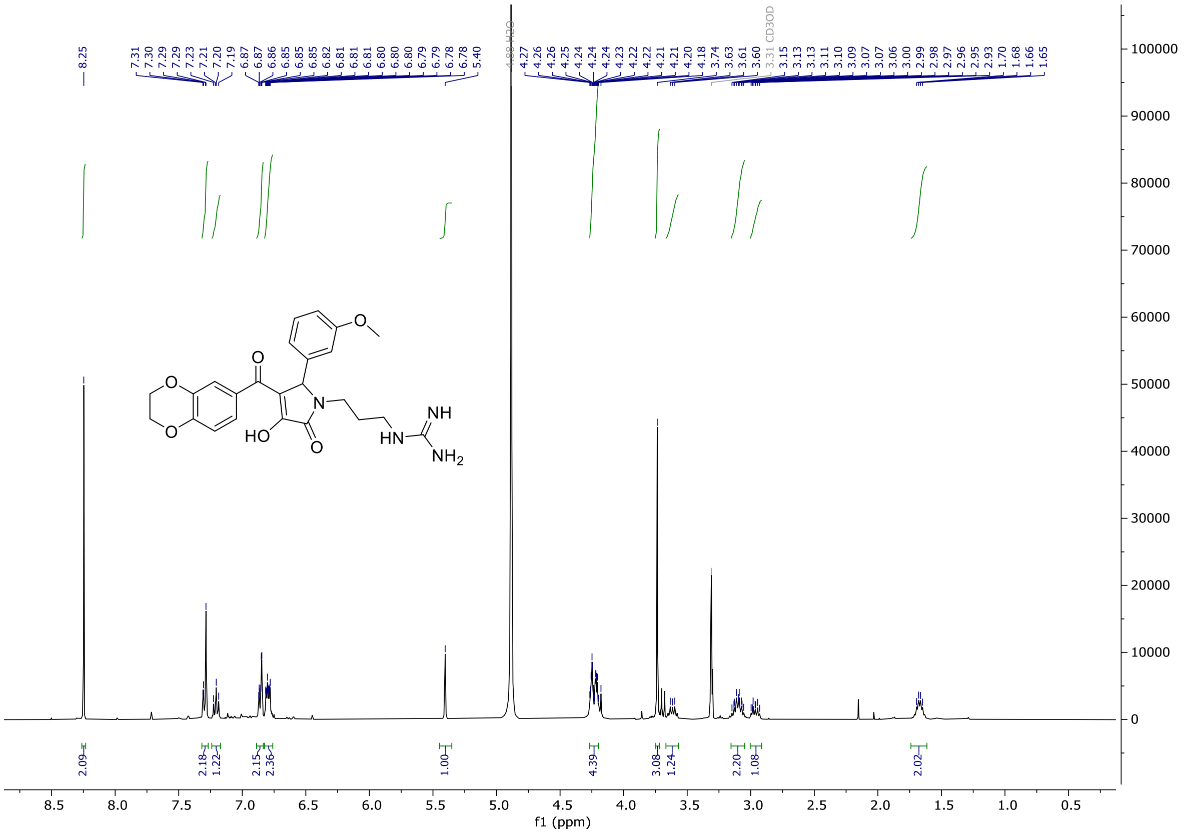

**Figure S16**. $^{13}C$ NMR ($CD_3OD$, 100 MHz) of **S4**.

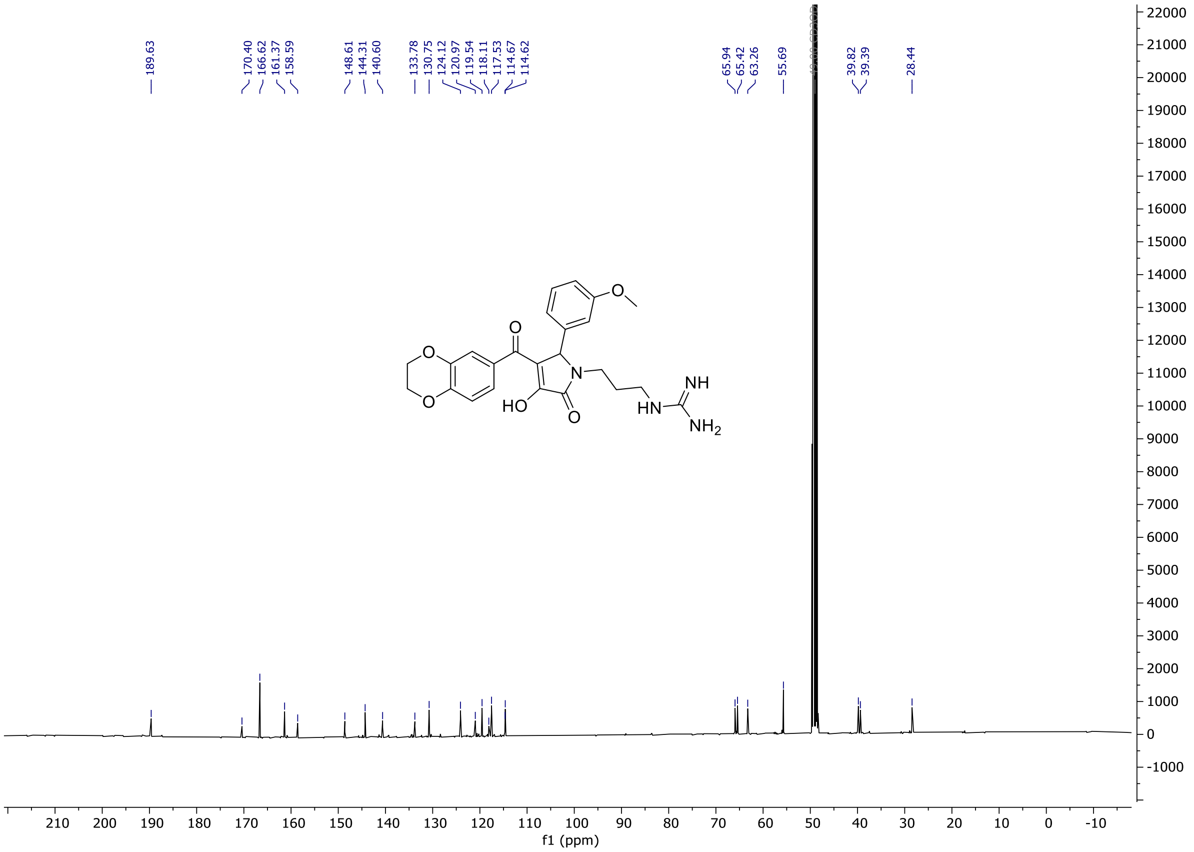

**Figure S17**. $^{1}$H NMR ($CD_3OD$, 400MHz) of **S5**.

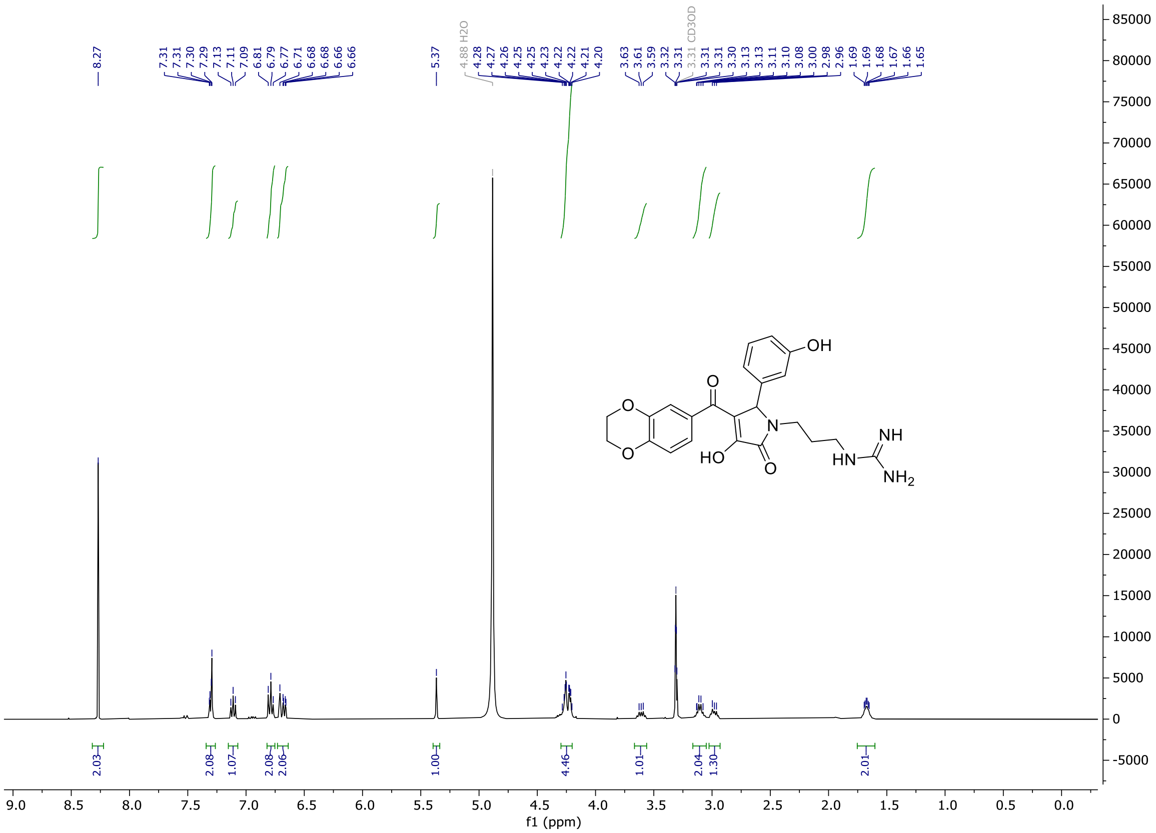

**Figure S18**. $^{1}H$ NMR ($(CD_3)_2SO$, 400 MHz) of **S6**.

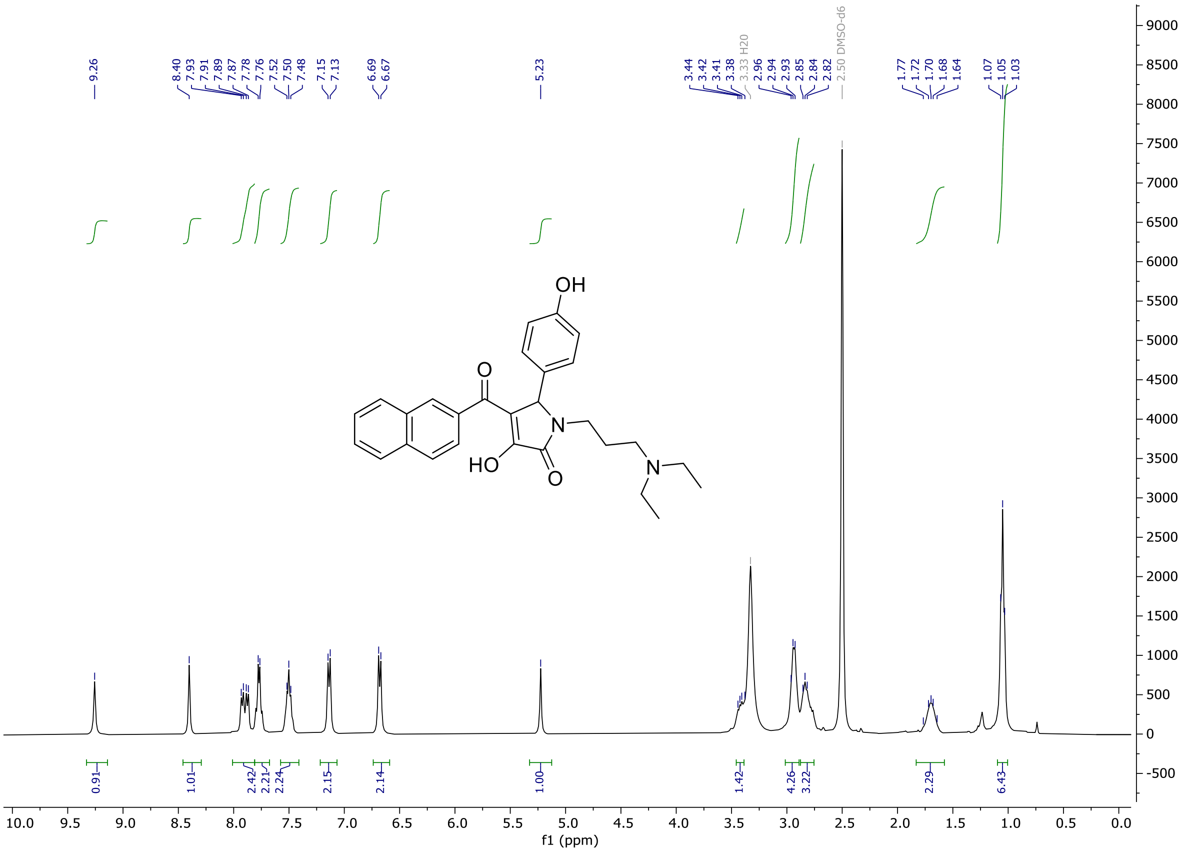

**Figure S19**. $^{13}C$ NMR ($(CD_3)_2SO$, 100 MHz) of **S6**.

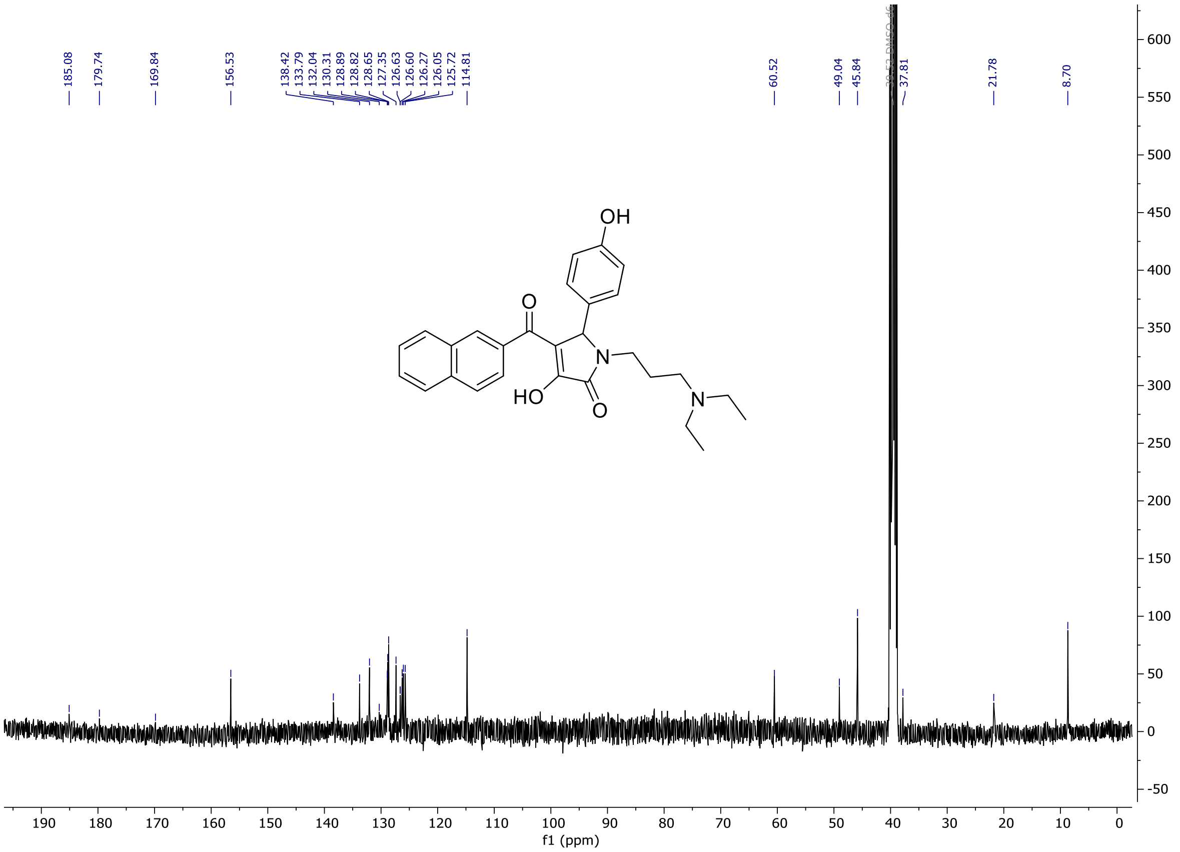

**Figure S20**. $^{1}H$ NMR ($(CD_3)_2SO$, 400 MHz) of **S7**.

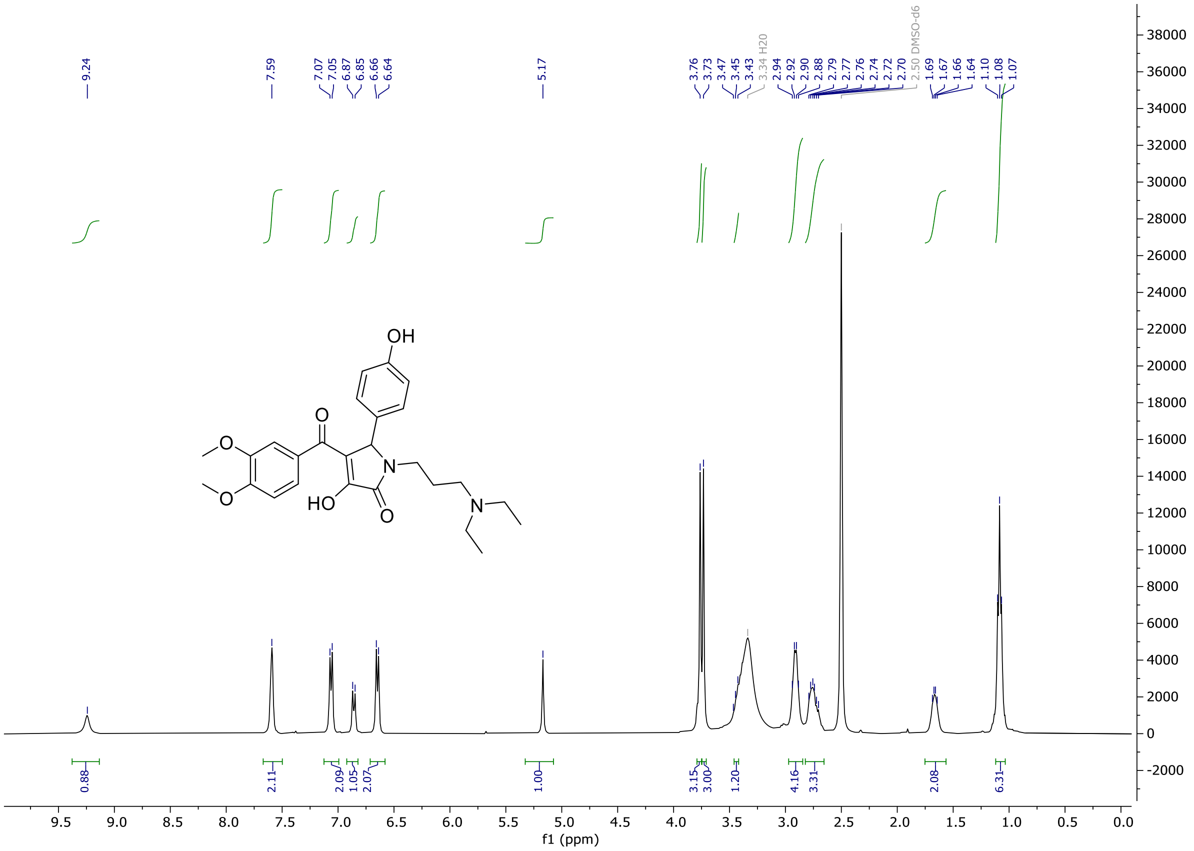

**Figure S21**. $^{13}C$ NMR ($(CD_3)_2SO$, 100 MHz) of **S7**.

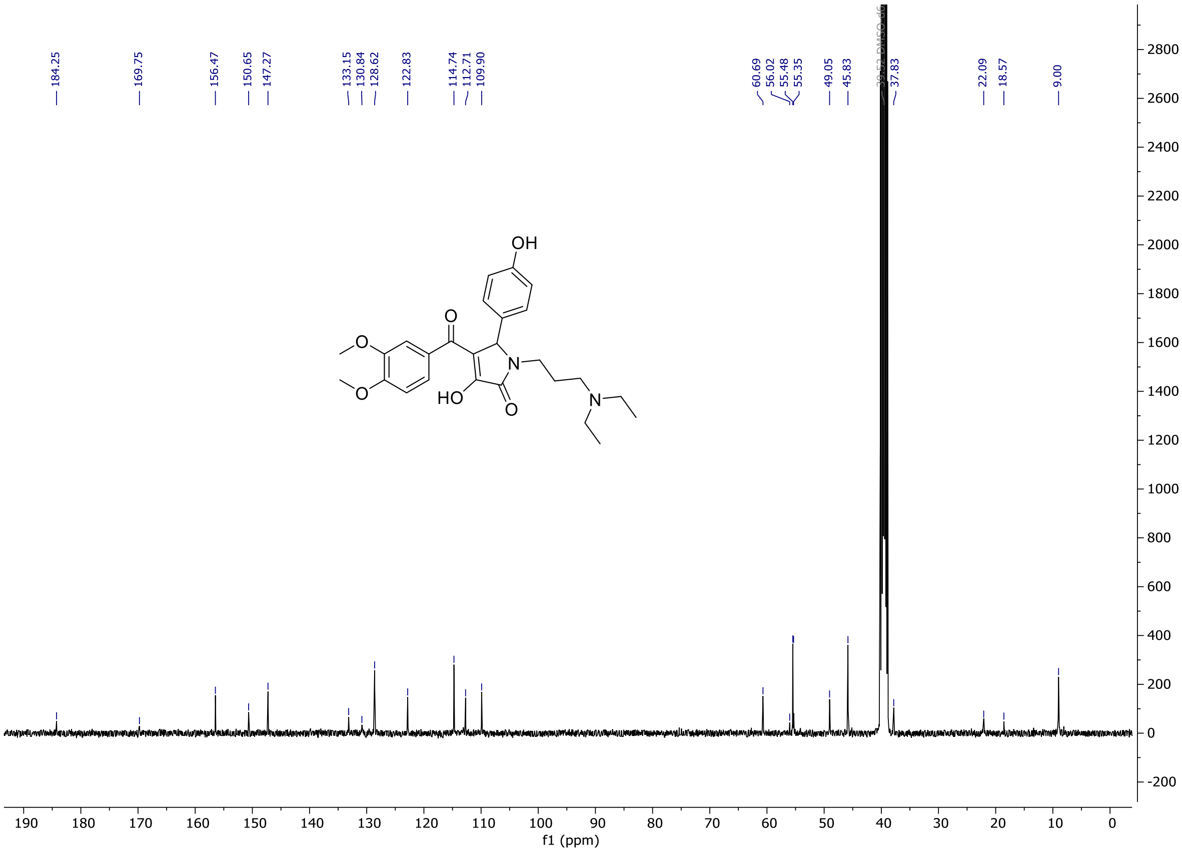

**Figure S22**. $^1H$ NMR ($CD_3OD$, 400 MHz) of **S8**.

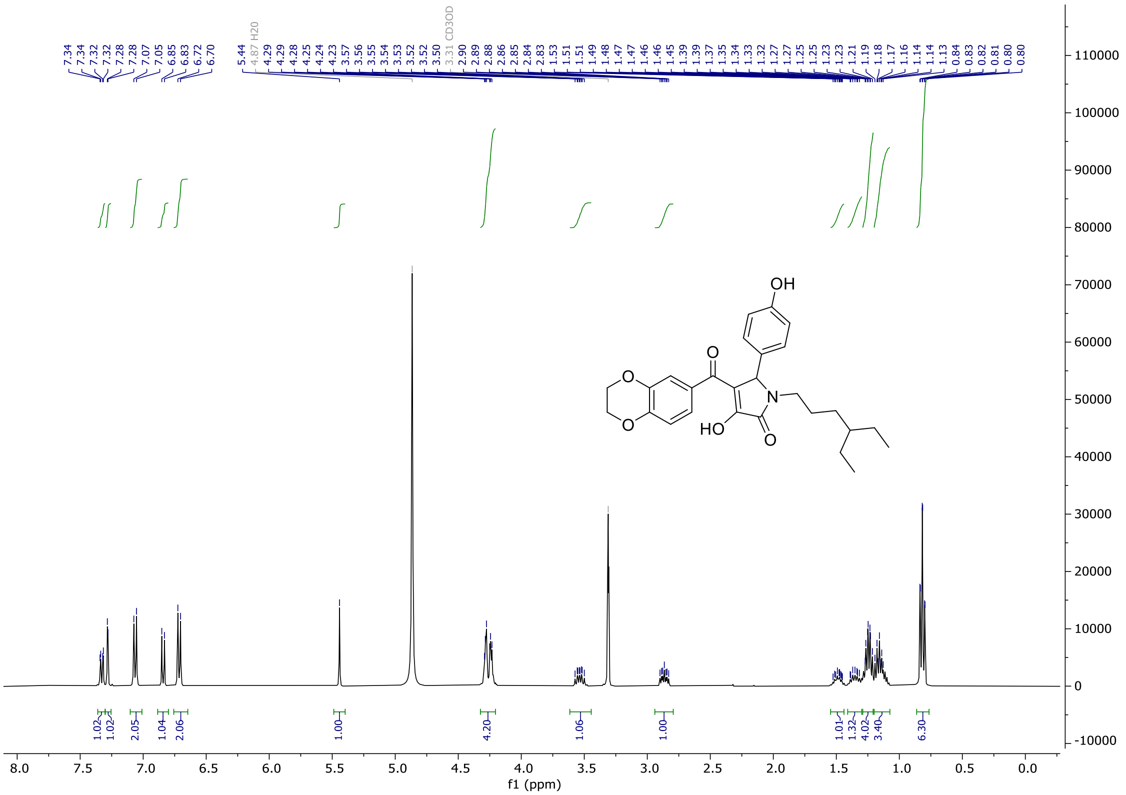

**Figure S23**. $^{13}$C NMR ($CD_3OD$, 100 MHz) of **S8**.

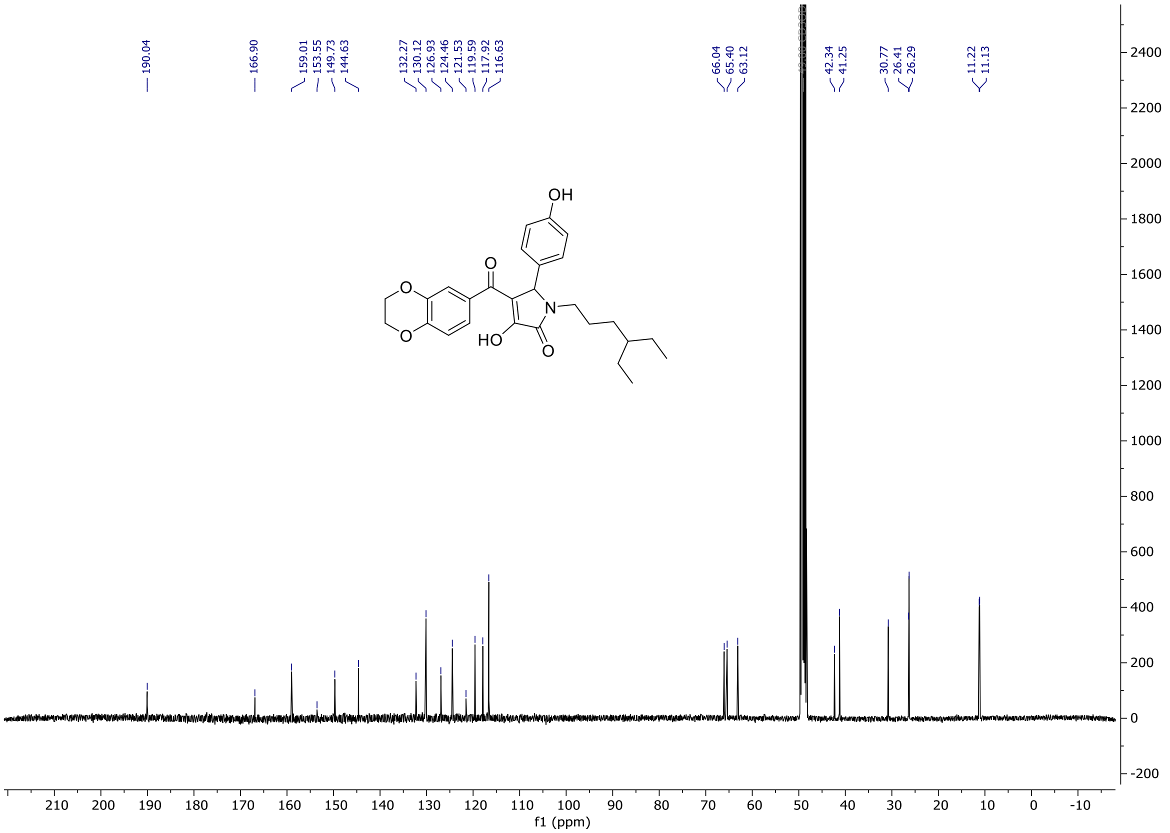

**Figure S24**. $^1H$ NMR ($(CD_3)_2SO$, 800 MHz) of **S9**.

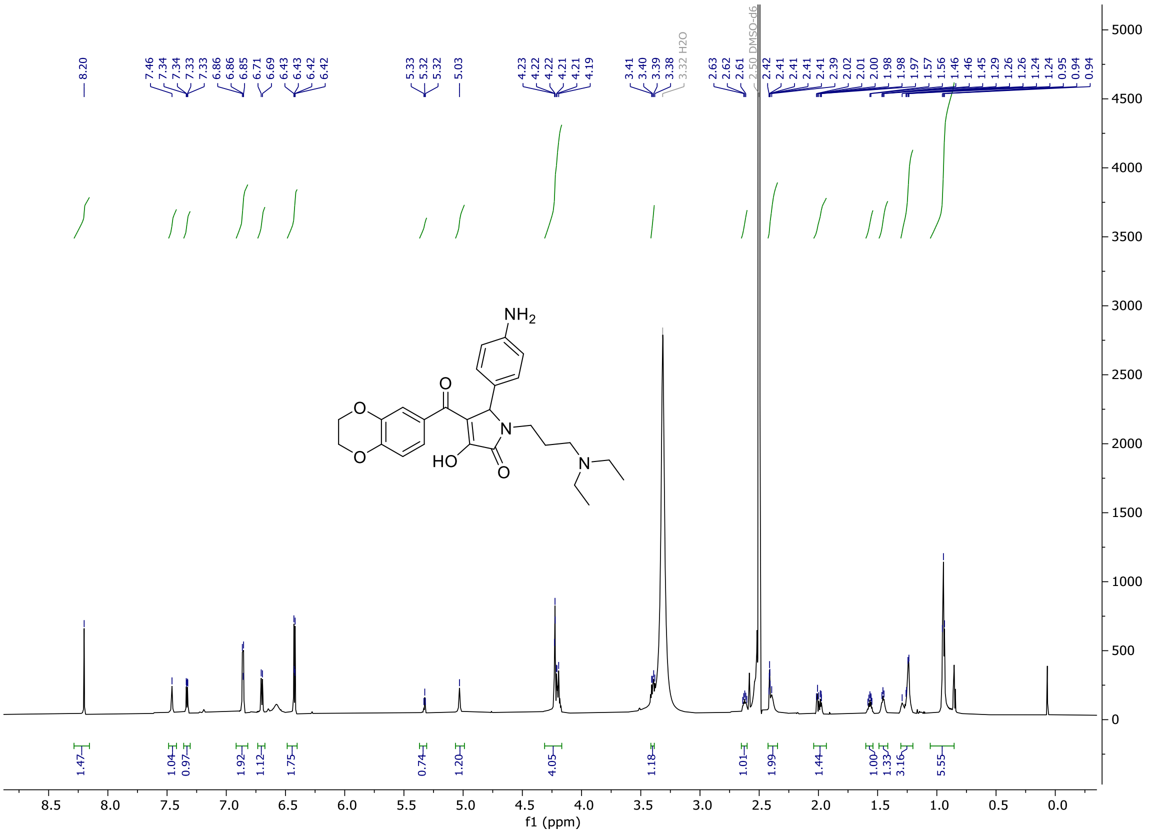

**Figure S25**. $^{1}H$-$^{13}C$ HSQC NMR ($(CD_3)_2SO$) of **S9**.

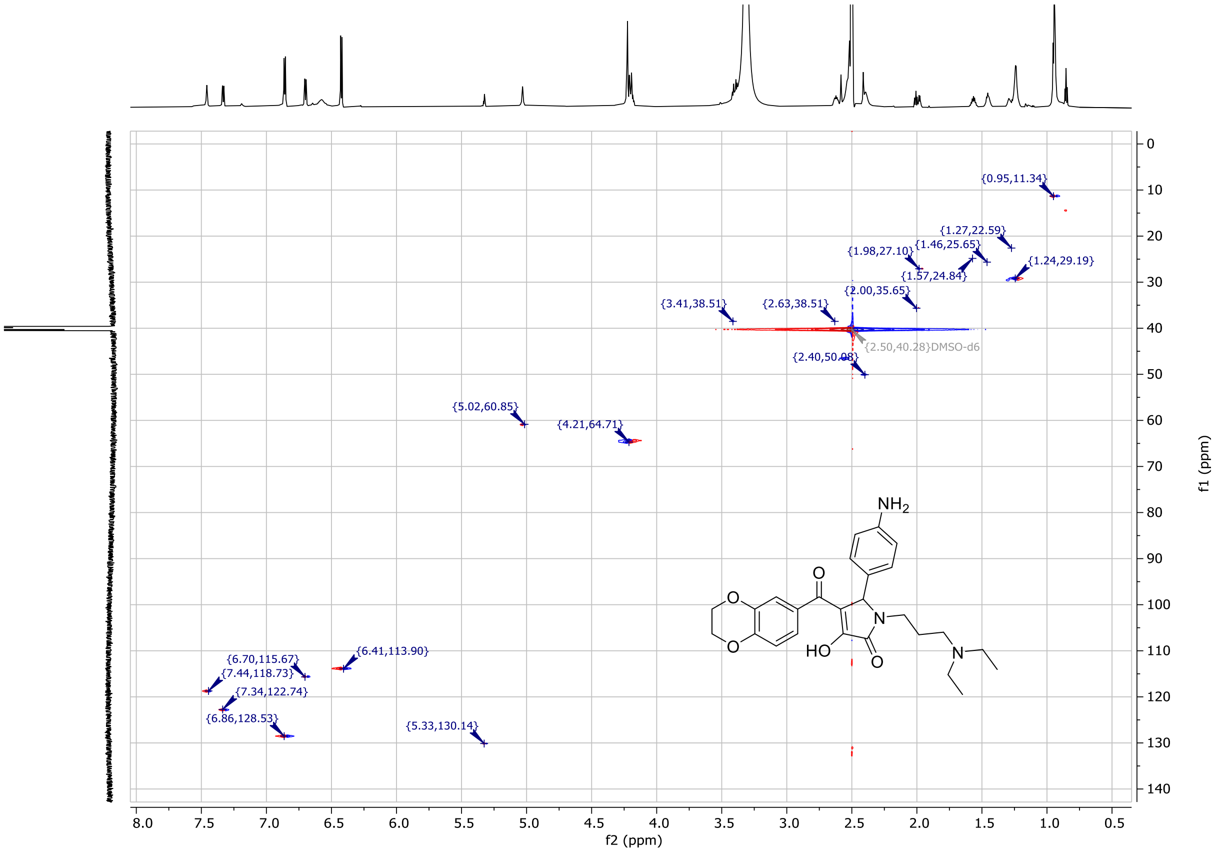

**Figure S26**. $^{1}H$-$^{13}C$ HMBC NMR ($(CD_3)_2SO$) of **S9**.

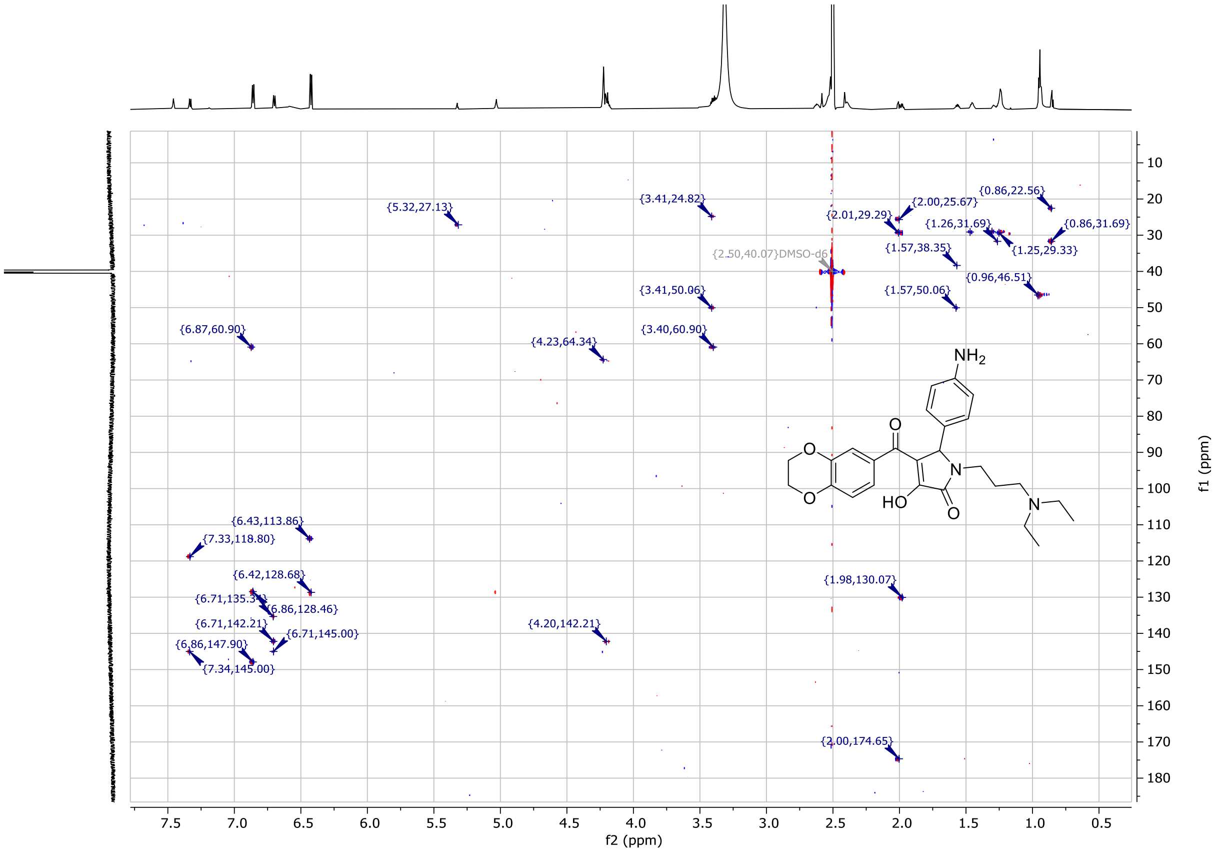

**Figure S27**. $^1$H NMR ($(CD_3)_2SO$, 400 MHz) of **S10**.

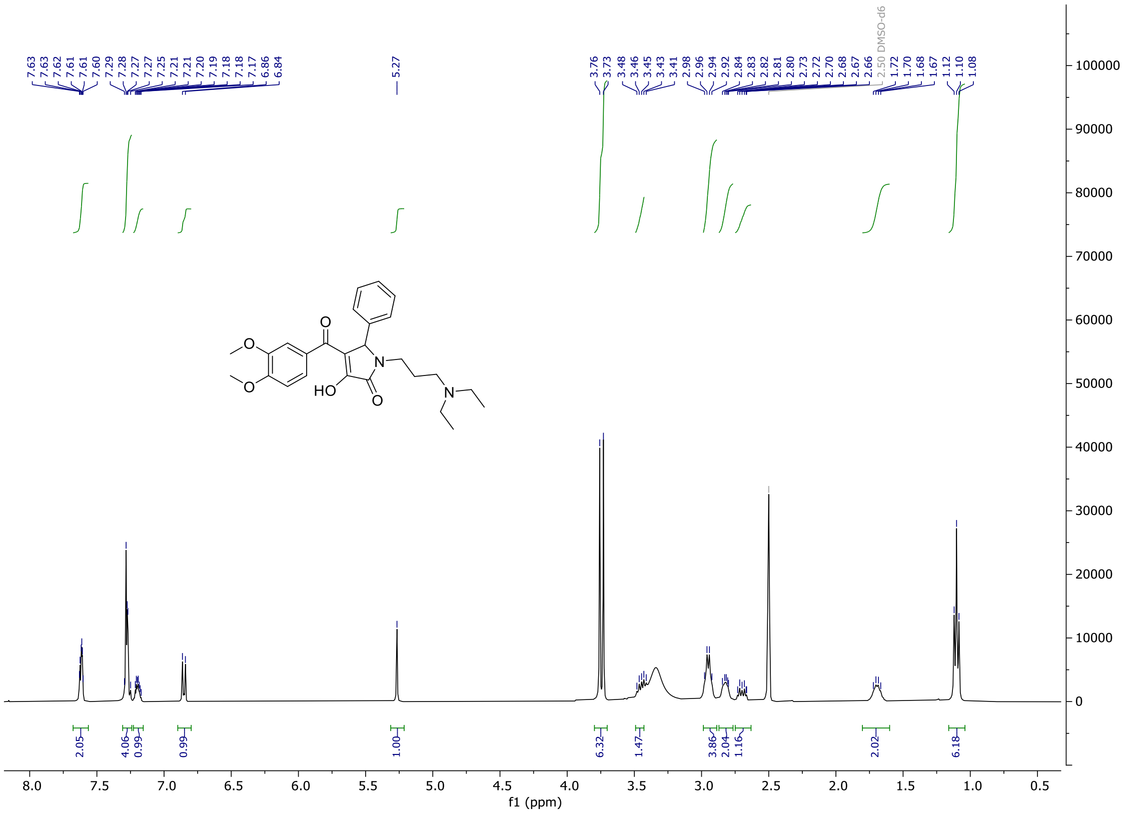

**Figure S28**. $^{13}$C NMR ($(CD_3)_2SO$, 100 MHz) of **S10**.

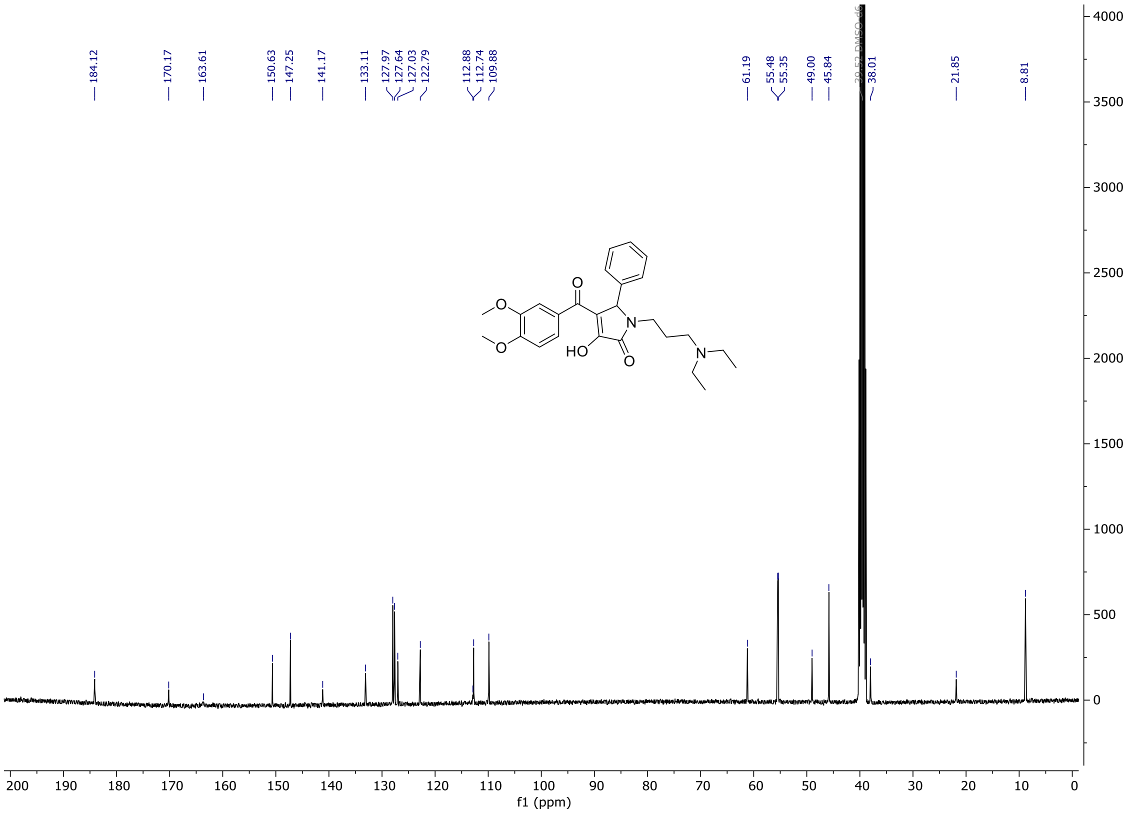

**Figure S29**. $^{1}$H NMR ($(CD_3)_2SO$, 400 MHz) of **S11**.

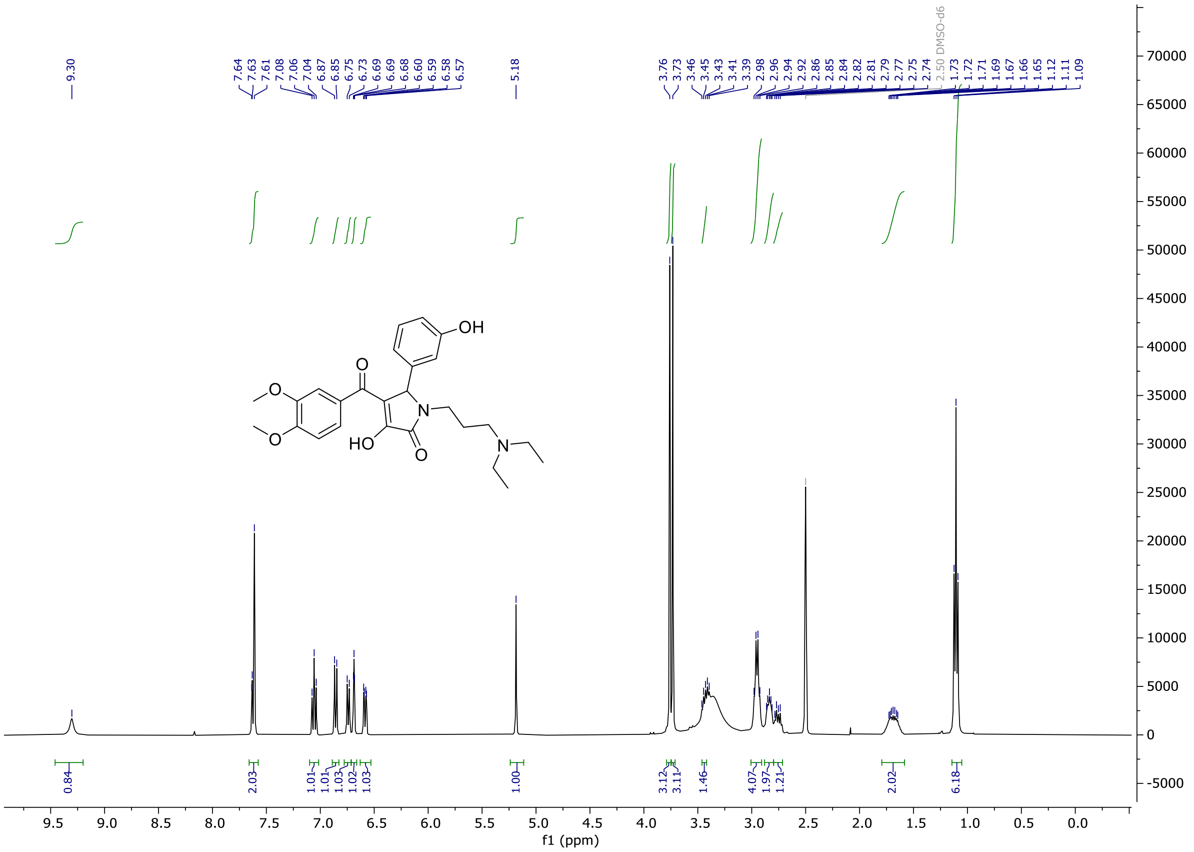

**Figure S30**. $^{13}C$ NMR ($(CD_3)_2SO$, 100 MHz) for **S11**.

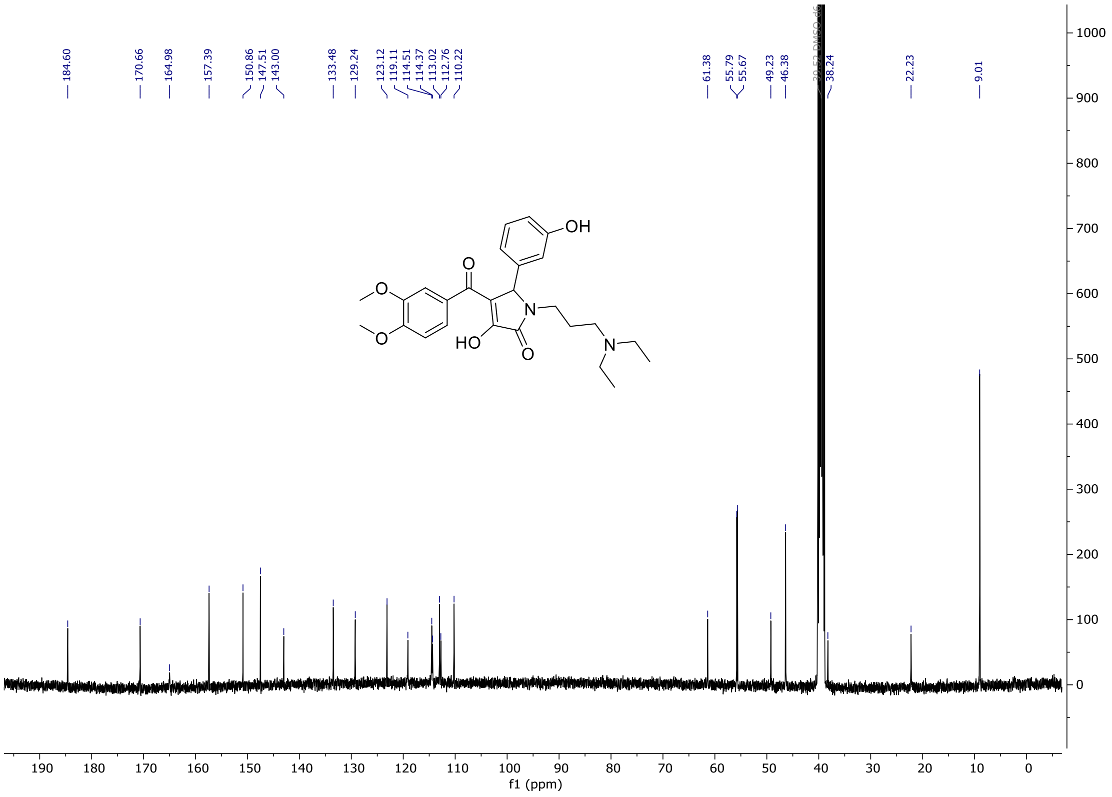

**Figure S31**. $^{1}H$ NMR ($CD_3OD$, 400 MHz) of **S12**.

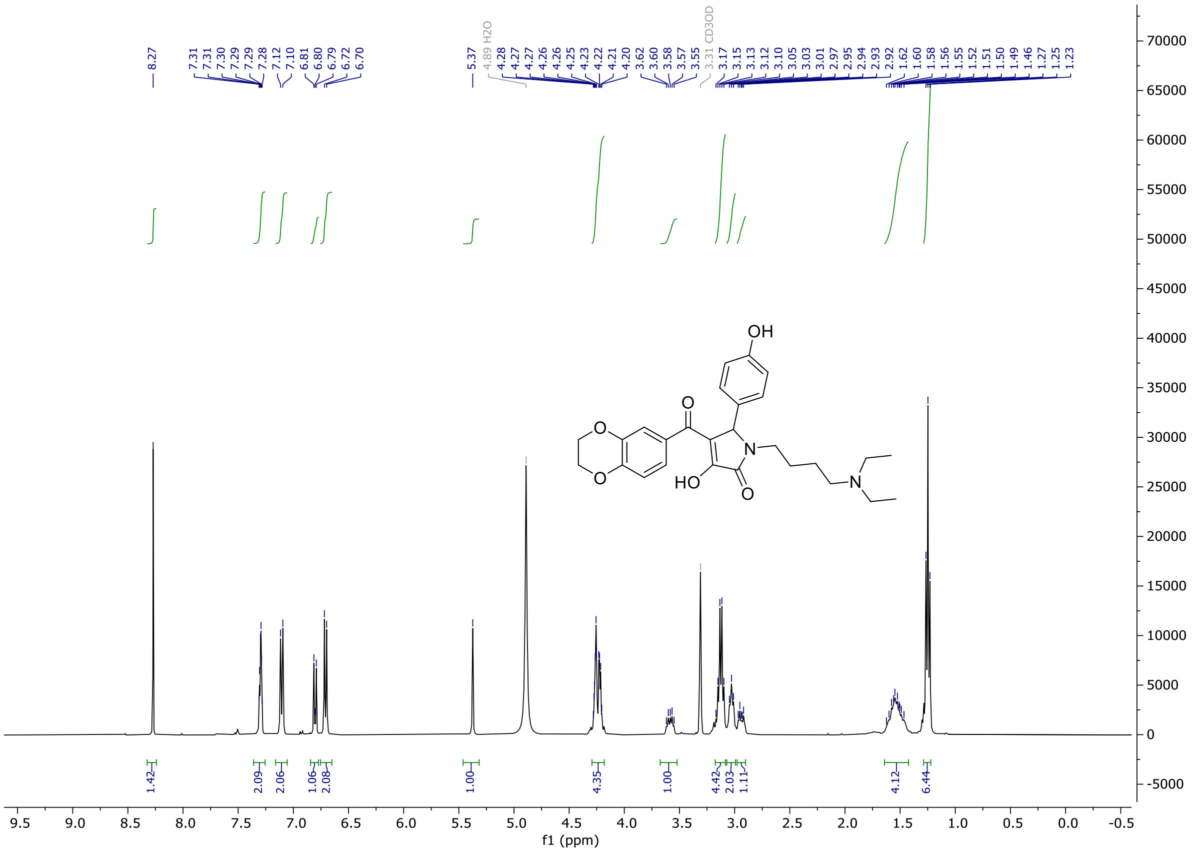

**Figure S32**. $^{13}C$ NMR ($CD_3OD$, 100 MHz) of **S12**.

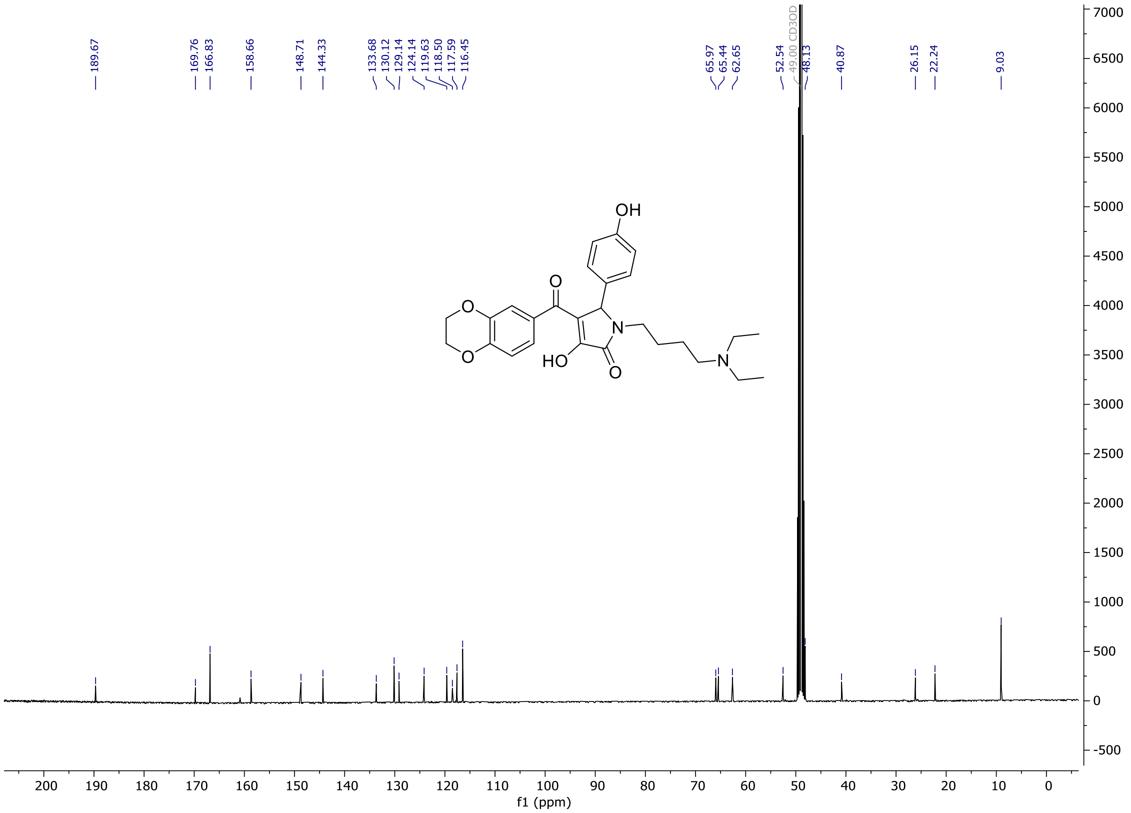

**Reference**

1. European chemical biology database. https://ecbd.eu/

2. Škuta C, Müller T, Voršilák M, et al. ECBD: European chemical biology database. *Nucleic Acids Research*. 2025;53(D1):D1383-D1392. doi:10.1093/nar/gkae904

3. Guettler S, LaRose J, Petsalaki E, et al. Structural Basis and Sequence Rules for Substrate Recognition by Tankyrase Explain the Basis for Cherubism Disease. *Cell*. 2011;147(6):1340-1354. doi:10.1016/j.cell.2011.10.046
